# Out-of-Anatolia: cultural and genetic interactions during the Neolithic expansion in the Aegean

**DOI:** 10.1101/2024.06.23.599747

**Authors:** Dilek Koptekin, Ayça Aydoğan, Cansu Karamurat, N. Ezgi Altınışık, Kıvılcım Başak Vural, D. Deniz Kazancı, Ayça Küçükakdağ Doğu, Damla Kaptan, Hasan Can Gemici, Eren Yüncü, Gülsün Umurtak, Refik Duru, Erkan Fidan, Özlem Çevik, Burçin Erdoğu, Taner Korkut, Christopher J. Knüsel, Scott Haddow, Clark Spencer Larsen, Rana Özbal, Fokke Gerritsen, Eylem Özdoğan, Uygar Ozan Usanmaz, Yasin Cemre Derici, Mine Uçmazoğlu, Flora Jay, Mehmet Özdoğan, Anders Götherström, Yılmaz Selim Erdal, Anna-Sapfo Malaspinas, Çiğdem Atakuman, Füsun Özer, Mehmet Somel

## Abstract

Western Anatolia has been a crucial yet elusive element in the Neolithic expansion from the Fertile Crescent to Europe. Using 30 new palaeogenomes from Anatolia c.8000-6000 BCE we describe the early Holocene genetic landscape of Western Anatolia, which reveals population continuity since the late Upper Pleistocene. Our findings indicate that the Neolithisation of Western Anatolia in the 7^th^ millennium BCE was a multifaceted process, characterised by the assimilation of Neolithic practices by indigenous groups and the influx of populations from the east, their admixed descendants eventually laying the foundations of Neolithic Southeast Europe. Intriguingly, the observed diversity in material culture among Aegean Early Neolithic communities correlates with their geographical distances but not their genetic differences, signifying a decoupling between cultural developments and genetic admixture processes.

## Introduction

The Neolithic transition to sedentism and food production in W Eurasia initially developed in SW Asia c.11000 BCE - 7000 BCE (Bar-Yosef, 2001; M. Özdoğan, 2024). Palaeogenomic studies suggest that this transition largely emerged through cultural experimentation by local foraging communities in different parts of the Neolithic core zones (the Levant, U Mesopotamia, Zagros, and C Anatolia) and their cultural exchanges, accompanied by limited levels of inter-regional mobility and genetic mixing (Broushaki et al., 2016; Feldman et al., 2019; Kılınç et al., 2016; Lazaridis et al., 2016). In contrast, the spread of Neolithic culture into Central and South Europe post-7000 BCE was driven by large-scale migrations, possibly originating from the Aegean region (Gamba et al., 2014; Hofmanová et al., 2016; Lipson et al., 2017; Mathieson et al., 2018; Skoglund, Sjödin, et al., 2014). Cultural interactions prevailed in other areas, such as Baltic foragers adopting pottery (Jones et al., 2017) or North African indigenous populations embracing agriculture through exchanges with Neolithic communities (Simões et al., 2023).

W Anatolia (i.e. the E Aegean coasts, the eastern Marmara, and the Pisidian Lakes District), has a unique role in this picture. The region was not among the Neolithic core zones, although it was possibly the origin of the “out-of-Anatolia” farming expansion into Europe and beyond post-7000 BCE (Hofmanová et al., 2016; Marchi et al., 2022; M. Özdoğan, 2024). Despite this key position, the sociocultural and demographic transformation of W Anatolia after 7000 BCE remains a major gap in our understanding of Neolithic development. In fact, we know little of W Anatolia during the initial Holocene, except that its coastal areas (E Aegean and Marmara littorals) harboured forager groups with cultural connections to neighbouring regions as inferred from their stone tools (Atakuman et al., 2022; Baird, 2012; Çilingiroğlu et al., 2020; Düring, 2013; Schoop, 2005). By c.9000 BCE, Aceramic/Pre-Pottery Neolithic (PPN) villages of sedentary foragers had emerged in the Fertile Crescent, including C Anatolia. Despite a lack of major geographic borders between C and W Anatolia, however, there was no visible Neolithic development in W Anatolia between c.9000-7000 BCE, apart from evidence for modest interregional contacts attested from stone tools (Aydıngün, 2009; Gatsov & Özdoğan, 1994; Nazaroff et al., 2013; Özbal & Gerritsen, 2019b). Only post-7000 BCE did fully sedentary villages start appearing in W Anatolia. By c.6500 BCE, these carried the full “Neolithic package” of permanently inhabited buildings, agriculture, domestic animals, pottery, and specific ritual elements that eventually dispersed into Europe [reviewed in (Brami, 2015; Düring, 2011; Karul, 2019; Özbal & Gerritsen, 2019b; E. Özdoğan, 2016; M. Özdoğan, 2024)].

The early Neolithic of W Anatolia (c.7000-6000 BCE) was an interesting phenomenon in itself, with villages that yielded remarkable diversity in their material culture and subsistence strategies (e.g. the simultaneous presence of round vs. rectilinear structures or the use or disregard of aquatic resources), a pattern that has been traditionally attributed to distinct regional affinities or to the variable influence of local foragers (Çevik & Vuruşkan, 2020; Düring, 2011; Erdogu & Çevik, 2020; Fidan et al., 2022; Gerritsen & Özbal, 2019; Karul & Avci, 2011; M. Özdoğan, 2024; Sari & Akyol, 2019). For instance, many NW Anatolian sites show both cultural and obsidian connections with Neolithic C Anatolia (Gerritsen & Özbal, 2019; Gemici et al., 2024), an indication that these villages were established by C Anatolian migrant farmers. Meanwhile, some NW Anatolian communities built round (rather than rectangular) buildings, a primitive type of construction long abandoned in C Anatolia; this mixed architecture has been interpreted as the outcome of integration with local W Anatolian foragers (M. Özdoğan, 2013). In CW Anatolia, villages exhibit yet other distinctive features, such as the presence of lime-plastered floors, the absence of round buildings, and the use of Melos (Greece) obsidian and maritime resources; these villages, in turn, have been suggested to be established by seafaring groups from the Levant (Horejs et al., 2015).

These models of population movement and admixture in W Anatolia have remained speculative, however, in the absence of conclusive archaeogenomic evidence. All published data from the region derive from c.6500 BCE NW Anatolia, and mostly from a single site, Barcın. The Barcın genomes have clear affinities to contemporaneous C Anatolians (e.g. Çatalhöyük), which might support the C Anatolian migration hypothesis. However, these NW Anatolian genomes have also been inferred to carry European Mesolithic-related ancestry, suggesting western admixture in Marmara by the early Holocene (Marchi et al., 2022). Without genetic or archaeological context, the verity of such western admixture has been unclear.

Overall, both the material culture and the limited palaeogenomics data remain equivocal as to the origins of the W Anatolian Neolithic. The genetic affinities of the populations who lived in the region pre-7000 BCE, whether W Anatolian Mesolithic and C Anatolian PPN communities between 9000-7000 BCE were genetically distinct or a homogeneous gene pool with distinct cultures, and how W Anatolian (E Aegean) Neolithic populations connected with the early Neolithic groups of modern-day Greece (W Aegean) are unknown. This has raised the question of whether the demographic expansion that drove European Neolithisation was originally out-of-Anatolia or rather out-of-the-Balkans/Greece (Kılınç et al., 2017). Here we address these points using new genomes from 30 individuals dated between 7800-6000 BCE, deriving from five W Anatolian settlements, as well as C Anatolian Çatalhöyük (Figure 1). Our dataset includes a human genome from Girmeler in SW Anatolia dated to 7800 BCE, the earliest in the region, with sociocultural affinities to PPN villages in C Anatolia (Takaoğlu et al., 2014). Analysing this genetic data, we describe the pre-7000 population of W Anatolia, its continuity and transformation through the Neolithic period, and its connections with the Neolithic in Greece.

**Figure 1:**
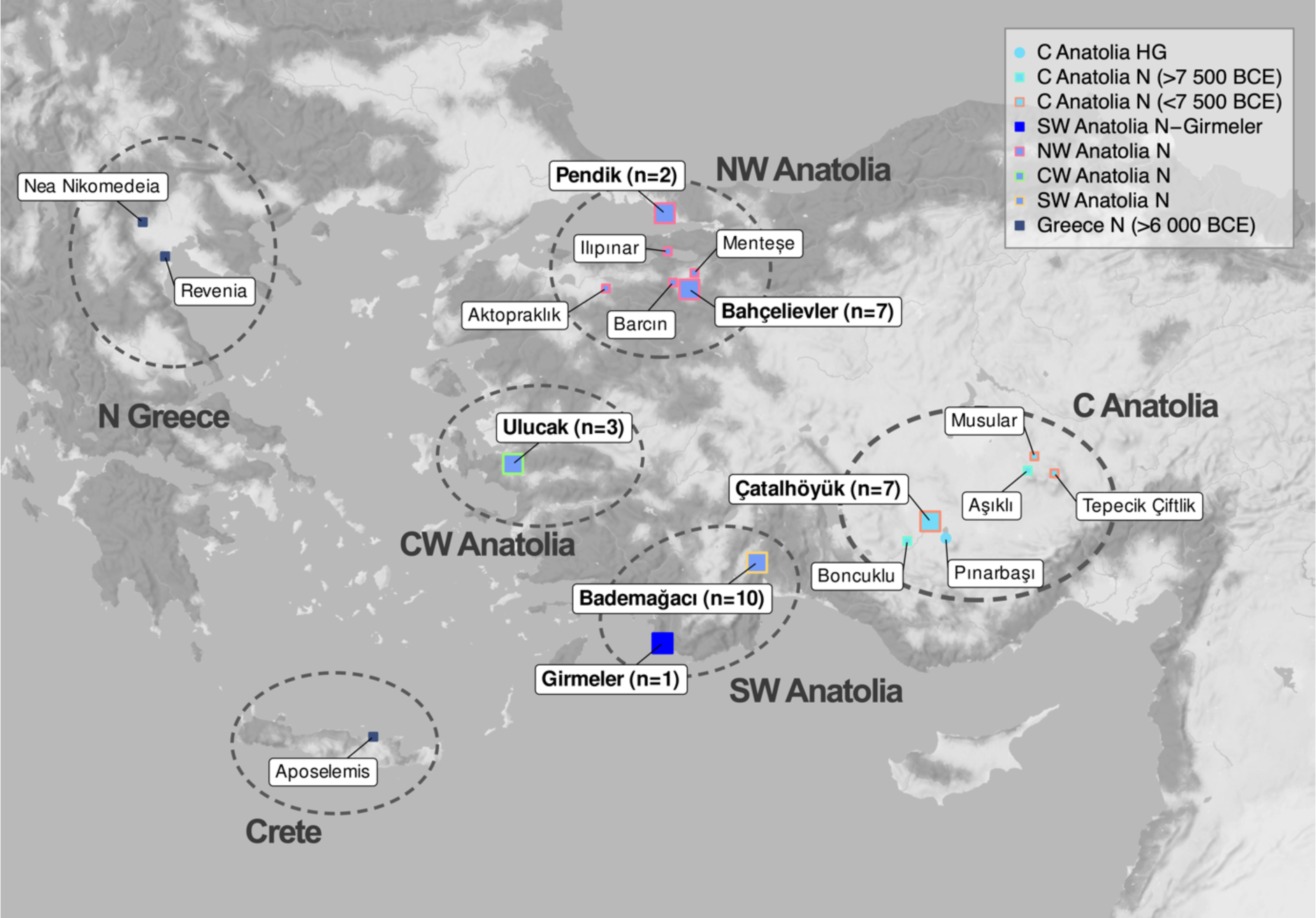
Map of Aegean and Anatolian sites >6000 BCE with palaeogenomics data analysed in this study. Smaller fonts and symbols indicate sites where only published genomic data have been included, while bold fonts and large symbols indicate sites where new palaeogenomic data has been produced, with sample sizes shown in parentheses (see also Table 1, Figure S1, Tables S1-2). We also used published data of 453 individuals from U Mesopotamia, the Balkans, Zagros, Levant, Cyprus and from European sites in our analyses, not shown in this figure. To improve visualization, some of the locations were slightly shifted (the exact site coordinates can be found in Table S1-2).

**Table 1.** Archaeological and genetic information of the ancient individuals generated in this study. The genomes have been sorted according to their archaeological dates. Genomes with sample ID shown with asterisks have been published previously but were enriched by further sequencing (Methods). The “N” suffix in the column “Region” indicates Neolithic (see also Tables S1-2). The individual bbh10 was not included in population genetic analyses because it had a close genetic kinship to bbh08 from the same site at Bahçelievler.

| Region | Sample ID | Skeletal codes | Location | Date (BCE, (2 $\sigma$)) | Average date (BCE) | Genome coverage | Genetic sex |
| --- | --- | --- | --- | --- | --- | --- | --- |
| SW Anatolia N | gir001 | S1 | Girmeler, Muğla, Turkey | 7738-7597 calBCE | 7668 | 0.11 | XX |
| C Anatolia N (<7500 BCE) | cch153 | 10527 | Çatalhöyük, Konya, Turkey | 6775-6700 BCE | 6738 | 1.06 | XY |
| C Anatolia N (<7500 BCE) | cch289* | 1885 | Çatalhöyük, Konya, Turkey | 6825-6635 calBCE | 6730 | 0.52 | XY |
| C Anatolia N (<7500 BCE) | cch294* | 2779 | Çatalhöyük, Konya, Turkey | 6700-6650 BCE | 6675 | 5.51 | XY |
| C Anatolia N (<7500 BCE) | cch251 | 2197 | Çatalhöyük, Konya, Turkey | 6600-6500 BCE | 6550 | 1.12 | XX |
| C Anatolia N (<7500 BCE) | cch311* | 8587 | Çatalhöyük, Konya, Turkey | 6600-6500 BCE | 6550 | 1.04 | XX |
| C Anatolia N (<7500 BCE) | cch348 | 2141 | Çatalhöyük, Konya, Turkey | 6600-6500 BCE | 6550 | 0.84 | XY |
| C Anatolia N (<7500 BCE) | cch245 | 12876 | Çatalhöyük, Konya, Turkey | 6400-6375 BCE | 6388 | 0.90 | XX |
| CW Anatolia N | ulu008 | B259/JDR | Ulucak Höyük, Izmir, Turkey | 6820-6597 calBCE | 6709 | 0.48 | XY |
| CW Anatolia N | ulu007 | B101 GSH/GTC | Ulucak Höyük, Izmir, Turkey | - | 6700 | 0.14 | XX |
| CW Anatolia N | ulu009 | B260J60 | Ulucak Höyük, Izmir, Turkey | - | 6700 | 0.11 | XY |
| NW Anatolia N | bbh001 | Bahçeli evler'20 iskelet No:1 | Bahçelievler, Bilecik, Turkey | 6612-6471 calBCE | 6542 | 1.40 | XY |
| NW Anatolia N | bbh007 | BAH21-A4/2-iskelet11 | Bahçelievler, Bilecik, Turkey | 6238-6391 calBCE | 6500 | 0.58 | XX |
| NW Anatolia N | bbh008 | BAH21-B5-24-iskelet7 | Bahçelievler, Bilecik, Turkey | - | 6500 | 0.15 | XX |
| NW Anatolia N | bbh009 | BAH21-B3-70/2-iskelet6 | Bahçelievler, Bilecik, Turkey | - | 6500 | 0.07 | consistent with XY but not XX |
| NW Anatolia N | bbh010 (2nd degree related with bbh008) | BAH21-C3-35-iskelet9 | Bahçelievler, Bilecik, Turkey | - | 6500 | 0.08 | XX |
| NW Anatolia N | bbh011 | BAH21-C3-38-iskelet10 | Bahçelievler, Bilecik, Turkey | - | 6500 | 0.66 | XY |
| NW Anatolia N | bbh012 | BAH21-iskelet4 | Bahçelievler, Bilecik, Turkey | - | 6500 | 0.08 | XY |
| SW Anatolia N | bad017 | BH ENÇ II/2 izole 1 | Bademağacı Höyüğü, Antalya, Turkey | 6389-6237 calBCE | 6313 | 3.67 | XX |
| SW Anatolia N | bad019 | BH'00 EN II/3 SK 13 A | Bademağacı Höyüğü, Antalya, Turkey | 6381-6222 calBCE | 6302 | 6.32 | XY |
| SW Anatolia N | bad022 | BH'00 EN II/3 SK 17 | Bademağacı Höyüğü, Antalya, Turkey | - | 6300 | 1.30 | XY |
| SW Anatolia N | bad023 | BH'00 EN II/3 SK 19 | Bademağacı Höyüğü, Antalya, Turkey | - | 6300 | 0.10 | consistent with XY but not XX |
| SW Anatolia N | bad024 | BH'02 EN 2 SK 2-4 | Bademağacı Höyüğü, Antalya, Turkey | - | 6300 | 1.54 | XX |
| SW Anatolia N | bad025 | BH'02 EN 24A House 2 | Bademağacı Höyüğü, Antalya, Turkey | - | 6300 | 1.44 | XY |
| SW Anatolia N | bad026 | BH'03 EN II/3 SK 20 | Bademağacı Höyüğü, Antalya, Turkey | - | 6300 | 2.03 | XY |
| SW Anatolia N | bad030 | BH'04 EN II/3 House 7 SK 34 | Bademağacı Höyüğü, Antalya, Turkey | - | 6300 | 1.76 | XY |
| SW Anatolia N | bad033 | BH'02 EN 2 SK 1 | Bademağacı Höyüğü, Antalya, Turkey | - | 6300 | 1.27 | XY |
| SW Anatolia N | bad034 | BH'06 EN II/1 Orta alan | Bademağacı Höyüğü, Antalya, Turkey | - | 6300 | 1.71 | XX |
| NW Anatolia N | pft004 | Pendik-Fikirtepe'81 123-6 | Pendik Höyük, Istanbul, Turkey | 6387-6230 calBCE | 6309 | 0.33 | XX |
| NW Anatolia N | pft003 | Pendik-Fikirtepe'81 2.5-10 | Pendik Höyük, Istanbul, Turkey | - | 6300 | 0.21 | XX |

## Results and Discussion

### The local population of W Anatolia before Neolithisation by 7000 BCE

The Girmeler site in SW Anatolia is located well outside the regions culturally assigned to the PPN (12^th^-8^th^ mil BCE) (Erdoğu, 2022; Takaoğlu et al., 2014) (Figure 1, Figure S1, Table 1). The mound has nevertheless revealed evidence for PPN-related activities, including sedentary structures such as a lime-plastered floor and hearths, sickle blades and glume wheats, and flexed burials, together indicating connections to late 9^th^/early 8^th^ millennium BCE PPN sites in C Anatolia (e.g. Aşıklı and Boncuklu) (Supplementary Information). Meanwhile, the lack of ovicaprid hunting or obsidian at Girmeler distinguishes it from the C Anatolian PPN (Erdoğu, 2022; Takaoğlu et al., 2014). The site also has parallels with Maroulas in the Aegean (Takaoğlu et al., 2014), suggesting interactions with Aegean foragers in addition to groups in the Neolithic core regions (Atakuman et al., 2022). However, whether the Girmeler community were locals or migrant PPN groups from C Anatolia, the Levant, Cyprus, or Greece, has remained unknown.

We genetically studied the best-preserved Girmeler skeleton. Endogenous aDNA content was only 0.7-0.9% in multiple libraries, from which we produced a 0.11x palaeogenome using shotgun sequencing (Methods). We radiocarbon-dated the individual to 7738 - 7597 calBCE (Table 1, Table S1). We then compared the Girmeler genetic profile with those from Late Upper Pleistocene (LUP) and Early Holocene (EH) populations from the broader region (Figure S1, Table S2). Here we chose to include published shotgun-sequenced genomes and not 1240K SNP capture data whenever possible; this was based on insights from our admixture simulations using a new algorithm, genoMIX (Methods), which confirmed that analyses involving shotgun and capture data types together a) can lead to false positive signals in f_4_-tests of admixture, and b) can weaken statistical power in admixture modeling using qpAdm (Methods; Figures S2-3).

In MDS, PCA and ADMIXTURE analyses the Girmeler genome consistently clustered with other LUP/EH Anatolians (shown in blue and purple) (Figure 2, Figures S4-6). Intriguingly, in MDS and PCA space Girmeler had the most similar behaviour with the 14th mil. BCE (Epipalaeolithic) C Anatolian Pınarbaşı genome (Feldman et al. 2019) and the 7^th^ mil. BCE NW Anatolian (Neolithic) Aktopraklık genome (Marchi et al. 2020) (Figure 2, Figures S4-5, Table S2). Despite the five millennia separating Girmeler and Pınarbaşı, their genetic profiles were so alike that all LUP/EH genomes from the broad region were equally distant to both in f_4_-tests (Figure S7A, Table S3). qpAdm modelling of genetic ancestry sources of the Girmeler genome further suggested that it carried mixed ancestry between Balkan- and Levant-related sources from the LUP/EH, again indistinguishable from the qpAdm model of Epipalaeolithic Pınarbaşı (Figure 3A, Figure S8A). Meanwhile, Girmeler was distinct from PPN Cyprus on the PCA (Figure S5); the Girmeler qpAdm model was also distinct from those of contemporaneous PPN C Anatolians, Aşıklı and Boncuklu (Figure 3A). In fact, the latter could instead be explained as admixed between c.65% Epipalaeolithic Pınarbaşı-related and c.35% U Mesopotamia PPN (Çayönü)-related genetic sources (Figure 3A, Table S4). Hence, this U Mesopotamia-related ancestry appears to have arrived and spread in C Anatolia sometime between 13-8k BCE, without reaching Girmeler (SW Anatolia) at detectable levels.

**Figure 2:**
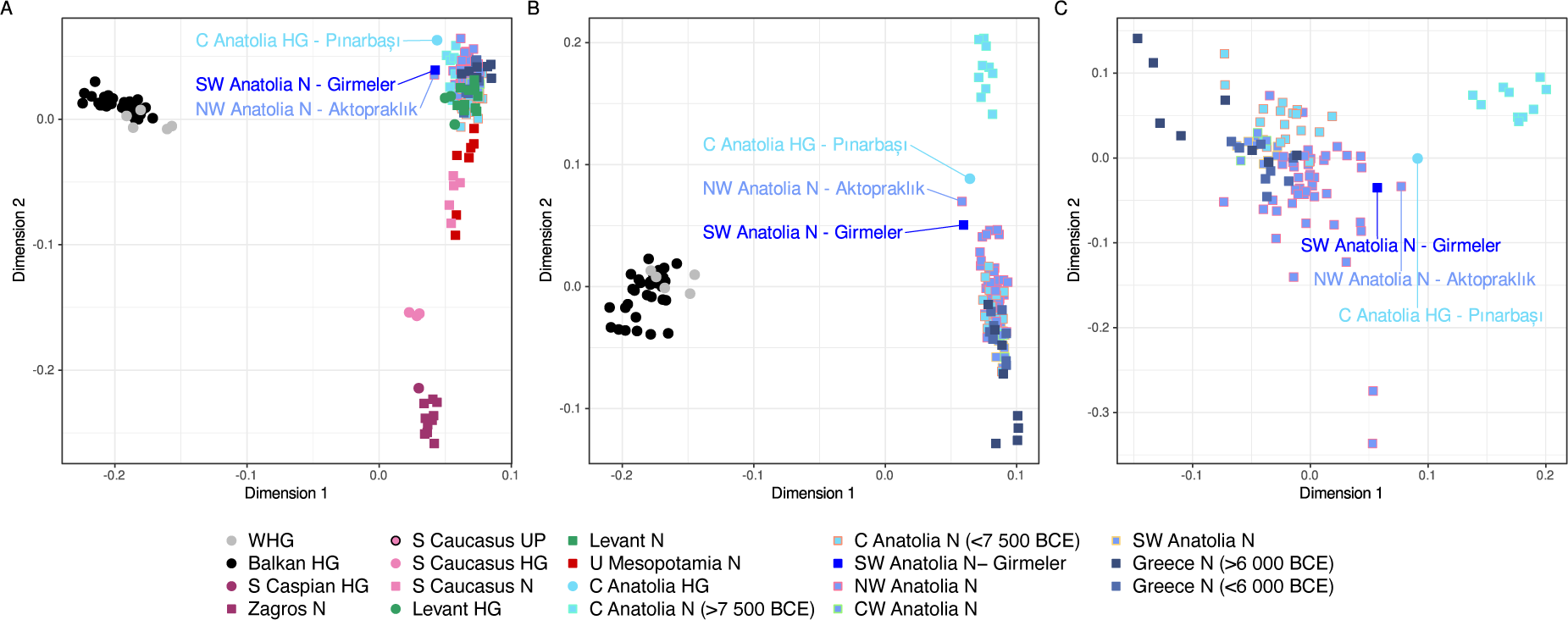
A multidimensional-scaling plot describing genetic distances among early Holocene individuals from the broader region. Distances were calculated using the outgroup-f_3_-statistic (Methods). Panels A to D have been drawn using progressively fewer populations to reveal broad-scale to fine-scale affinity patterns. “UP”, “HG” and “N” suffixes indicate genomes of individuals from “Upper Palaeolithic”, (Mesolithic) “hunter-gatherer” and “Neolithic” contexts, respectively. EHG and WHG stand for genomes from “East European” and “West European” Mesolithic contexts. Note that genomes with low coverage (e.g. Cyprus) could not be included in this analysis but are included in the projection-based PCA (Figure S5).

**Figure 3:**
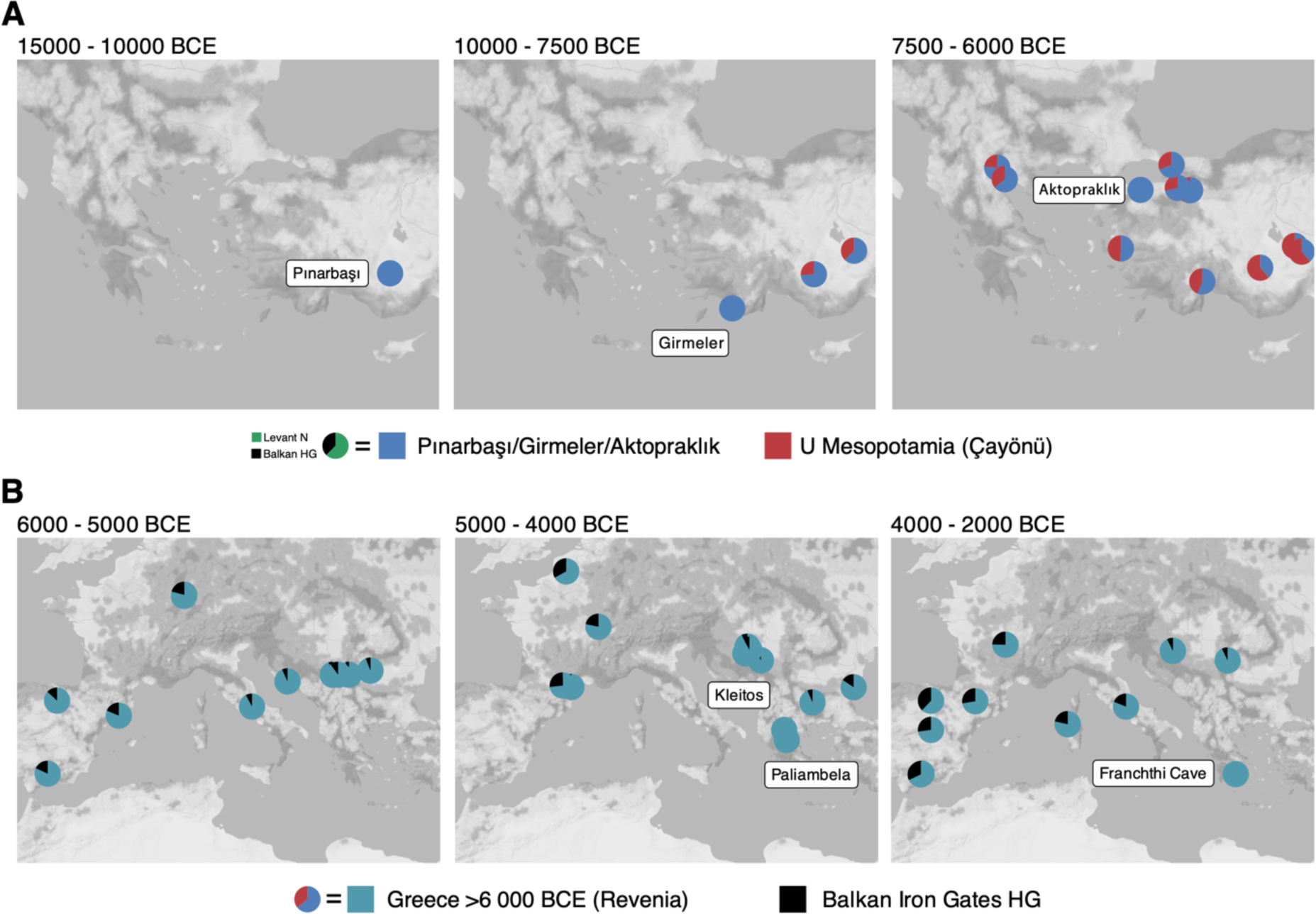
qpAdm models of genomes from Anatolia and the Aegean (A), and S/C Europe (B). The pie-charts show ancestry proportions of genomes from archaeological sites estimated using statistically “feasible” qpAdm models (Methods). The time windows of the genomes included are indicated on top of the panels. The ancestry sources used in the models are shown in the key. The key also shows the qpAdm models for the genetic profiles of Pınarbaşı/Girmeler/Aktopraklık (with Levant N and Balkan HG sources) and of pre-6000 BCE Greece represented by Revenia and Nea Nikomedeia with Pınarbaşı/Girmeler/Aktopraklık and U Mesopotamia sources) (see Table S4). qpAdm models of Neolithic Aposelemis (Crete) did not yield significant results and are therefore not shown. To improve visualization, some of the locations were slightly shifted (the exact site coordinates can be found in Table S1-2).

Even though Girmeler is a single genome, its genomic profile is conspicuously distinct from any of its contemporaries from the core regions of the PPN. This implies that the Girmeler individual was not a migrant, but a descendant of a local population that was experimenting with sedentism and Neolithic-related cultures, in cultural connections with its eastern neighbours albeit without visible genetic interaction.

### The mixed origins of W Anatolian Neolithic villages

The mid-8th mil. BCE was a demographic turning point in the Neolithic development of Anatolia, when a second wave of eastern mobility possibly of U Mesopotamian origin (Altınışık et al., 2022) arrived in C Anatolia, as captured in Musular genomes (Koptekin et al., 2023) (Figure 3A). By the early 7th mil. BCE, villages with pottery, agriculture and animal husbandry were being established across the Fertile Crescent. But this time, they continued to spread beyond the core zones, into the W Anatolia and the wider Aegean. The simultaneous emergence of the so-called “Neolithic package” elements in the Aegean has been interpreted as indicative of emigration from the Neolithic core (M. Özdoğan, 2011). Analysis of archaeological data has suggested various possible routes of human mobility/interaction that linked the emergence of W Anatolian Neolithic with its neighbours: one originating in C Anatolia and/or U Mesopotamia and reaching CW Anatolia via SW Anatolia, another reaching NW Anatolia directly from C Anatolia and/or U Mesopotamia, and also a maritime route connecting the Levant with SW/CW Anatolia (Horejs et al., 2015; M. Özdoğan, 2013, 2024). Hitherto these models could not be tested genetically due to the scarcity of representative palaeogenomes.

Together with 29 new genomes post-7000 BCE with coverages 0.08-6.32x (median=0.9x) (Figure 1, Table 1, Table S1, Supplementary Information), our dataset now covers W and C Anatolia between 7500-6000 BCE with 104 genomes and comprises 11 settlements: Bademağacı in SW Anatolia, Ulucak in CW Anatolia, Bahçelievler, Pendik, Barcın, Aktopraklık, Menteşe and Ilıpınar in NW Anatolia, Musular, Çatalhöyük, and Tepecik-Çiftlik in C Anatolia. We compared this data with partial genomes from earlier or contemporary sites, which revealed four notable observations.

We found that W Anatolian genomes are overall similar to contemporaneous (post-7500 BCE) genomes from C Anatolia, as previously noted (Kılınç et al., 2017). C and W Anatolian genetic affinity post-7500 BCE can be observed both in MDS and PCA plots (Figure 2, Figures S4-5) and also in qpAdm models, where nearly all C and W Anatolians studied can be modelled as admixed between Pınarbaşı/Girmeler and Çayönü (U Mesopotamia) (Figure 3A). A considerable fraction of the ancestors of W Anatolian Neolithic villages may thus have immigrated some centuries ago from C Anatolia and/or genetically unsampled regions with possibly similar genetic profiles, such as the U Euphrates or N Levant. In fact, the presence of domestic pigs or pressure flaking in CW Anatolian sites such as Ulucak, when these were absent in C Anatolia (Supplemental Information) (Guilbeau et al. 2019; M. Özdoğan, 2024), may support an U Euphrates contribution instead of C Anatolian origins. Meanwhile, a S Levantine origin that could have arrived via seafaring is highly unlikely given their distinct genetic profiles (Figures S4-5). We also found that Cyprus PPN (Heraclides et al., 2024) had an equal affinity to SW, CW and NW Anatolians in f_4_-tests. Given the expectation that a maritime expansion from the Cyprus/Levant might more strongly impact CW/SW Anatolia rather than the physically and culturally more distant NW Anatolia, the f_4_ results cast doubt on the idea that W Anatolia was settled via maritime expansion (Figure S9).

A surprising exception to the general affinity between W and C Anatolia involved the Aktopraklık individual coded AKT16, the oldest NW Anatolian in our dataset (6658-6578 calBCE) (Marchi et al., 2022). AKT16 not only stood out in MDS and PCA analyses as being different from other W Anatolians (Figure 2) but was indistinguishable from Pınarbaşı (C Anatolia - 13642-13073 calBCE) and Girmeler (SW Anatolia - 7738-7597 calBCE) in qpAdm models and f_4_-tests (Figure 3, Figures S7-8); it also clustered with Pınarbaşı in haplotype-sharing analyses with FineSTRUCTURE (Lawson et al., 2012) (Methods) (Figures S10-11). Low-coverage genome-wide capture data from three other 7^th^ millennium BCE Aktopraklık individuals (Hofmanová et al., 2022) also indicated high AKT16-like ancestry in these individuals, although Late Neolithic (post-6000 BCE) Aktopraklık capture genomes did not carry this additional AKT16 affinity (Figure S12). These results suggest that AKT16 was not the product of recent admixture between European and Anatolian Early Holocene groups, as previously proposed (Marchi et al., 2022). Instead, the Early Neolithic Aktopraklık population likely descended from the same pre-Neolithic gene pool as Girmeler, which may have inhabited all of W Anatolia into the 7^th^ mil. BCE.

We found that NW and SW Anatolian 7th mil. Neolithic genomes (apart from Aktopraklık and Ilıpınar) could be modelled using qpAdm with indigenous W Anatolians (Girmeler/Aktopraklık) (28-53%) and post-7500 BCE C Anatolia (Çatalhöyük) (47-72%) as sources (Figure S8B). We also found Pınarbaşı/Girmeler/Aktopraklık genomes to show higher affinity to W Anatolians than to contemporary C Anatolians: all 126 tests of the form *f_4_(Yoruba, Pınarbaşı/Girmeler/Aktopraklık; <7500 BCE C Anatolia, <7500 BCE W Anatolia)* were positive, with 56% of these significant at Z>3 (Figure 4, see also Figures S10-11 and Figures S13-14). These results strongly suggest that indigenous W Anatolians admixed with incomers from C Anatolia and/or from its east, to eventually create the Neolithic W Anatolian gene pool identified in villages such as Bademağacı, Ulucak, Barcın, Bahçelievler, or Pendik.

**Figure 4:**
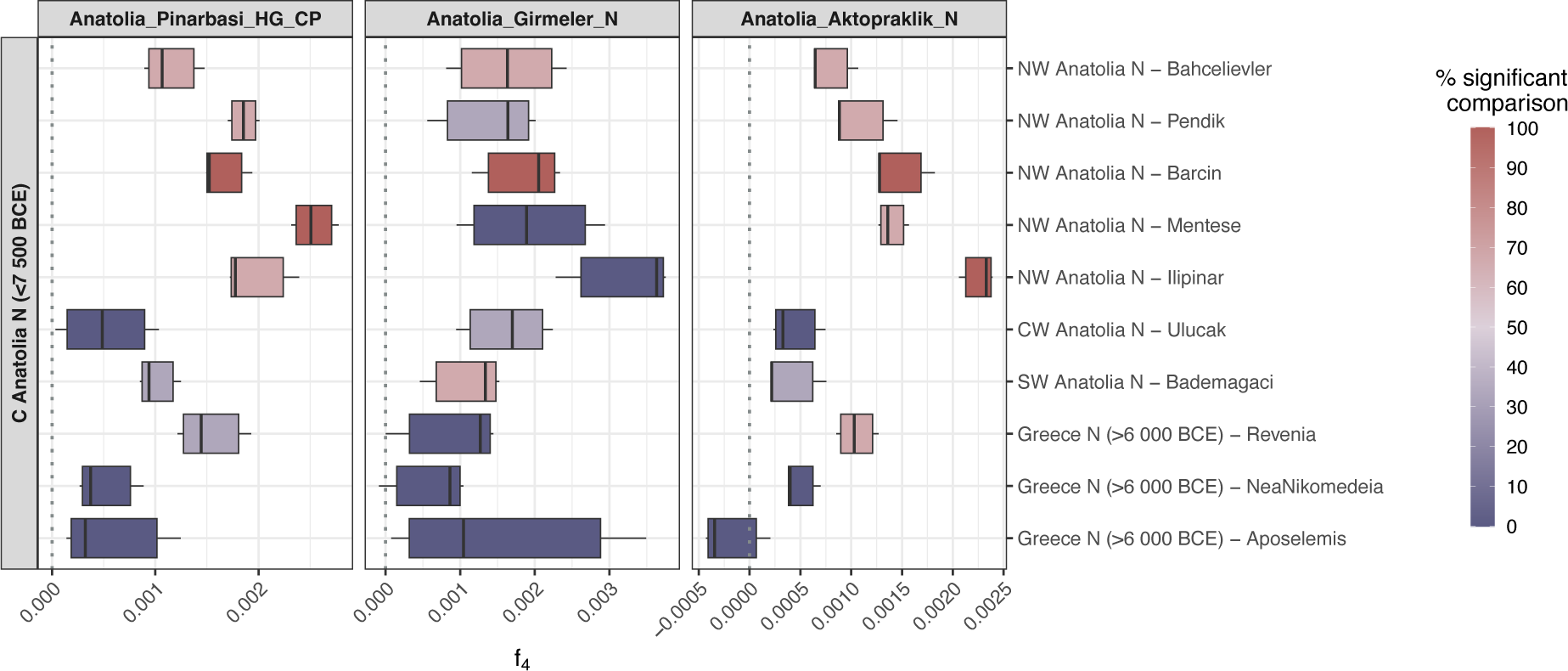
The affinities of Epipalaeolithic Pınarbaşı, Girmeler and Aktopraklık genomes to C Anatolia versus W Anatolia/Greece analysed using f_4_-tests. The populations on the left side are C Anatolians post-7500 BCE (Çatalhöyük, Musular, Tepecik-Çiftlik) while those on the right side are Neolithic groups from W Anatolia or Greece. The boxplots show the distribution of the f_4_-statistics performed using groups of genomes per site; e.g. the top left comparison involves three comparisons of the form *f_4_(Yoruba, Pınarbaşı/Girmeler/Aktopraklık; Çatalhöyük/Musular/Tepecik-Çiftlik, W Anatolia/Greece)*, one including Çatalhöyük, one Musular, and one Tepecik-Çiftlik. Colour coding indicates the proportion of nominally significant tests out of all comparisons (at |Z|>3). The “_HG” and “_N” suffixes indicate “hunter-gatherer” and “Neolithic”-related populations, respectively. The “CP” suffix stands for aDNA data produced using capture technologies (instead of shotgun).

We further observed considerable regional and intra-site heterogeneity in these indigenous admixture patterns. Both f_4_-tests, qpAdm modeling, f_3_- and haplotype-sharing analyses indicated higher Pınarbaşı/Girmeler/Aktopraklık ancestry in NW Anatolia than in SW/CW Anatolia, on average (Figure 4, Figures S15-19). The Ulucak genomes (c.6700 BCE), the oldest W Anatolians after Girmeler in our dataset, had lower indigenous affinity than most of their NW Anatolian contemporaries (Figures S16-17). Among the analysed NW Anatolian shotgun palaeogenomes, some showed higher indigenous ancestry than others, with heterogeneity even within the same site (Figure S17), although indigenous ancestry levels were not correlated with the archaeological date of the individuals (Figure S20). These observations overall mark the variability of the admixture process, both in space and time.

Our results have major implications. First, the Aktopraklık data suggests that a Pınarbaşı/Girmeler-like population persisted in W Anatolia till the mid-7th mil. BCE, with little or no demographic influence from its east, despite cultural contacts (Özbal & Gerritsen, 2019b). Second, between c.7500 and c.6500 BCE W Anatolia received gene flow from C Anatolia or from more eastern regions (M. Özdoğan, 2024), either gradually or by mass mobility, and these incoming groups were a major demographic contributor to the first Neolithic villages in the Aegean. Third, both the Aktopraklık genomes and the Pınarbaşı/Girmeler/Aktopraklık-like ancestry in W Anatolia attests to indigenous involvement in the development of the W Anatolian Neolithic (Karul, 2017), as well as admixture between incoming groups and locals. Fourth, local groups may have fully mixed with incoming groups within a few centuries, such that no genomes from post-6500 BCE W Anatolia matches the distinct autochthonous profile represented by AKT16 (c.6600 BCE). Finally, we find no apparent connection between material culture variation among W Anatolian villages and their genetic history, a point we develop later.

### The out-of-Anatolia expansion and lack of indigenous admixture in Greece

Our data further provides new insights into the Neolithisation of the W Aegean, i.e. present-day Greece. Based on material culture analyses, some scholars suggested that Greece was colonised by Levantine agriculturalists moving along the coast (Perlès, 2005), while others pointed to Anatolian origins (E. Özdoğan, 2016). The observation that European Neolithic-associated genetic profiles were highly similar to those of Neolithic W Anatolians, in turn, pointed towards a Neolithic out-of-Anatolia event (Hofmanová et al., 2016; Marchi et al., 2022; Mathieson et al., 2015). However, given the lack of pre-Neolithic genomes from the Aegean, we had previously speculated whether early Neolithic groups in W Anatolia and Greece could have been acculturated descendants of indigenous foragers with that same genetic profile (Kılınç et al., 2017). This would imply that the demographic origins of the European Neolithic expansion lay in SE Europe instead of Anatolia. We can now reject this model: the European Neolithic-like genetic profile emerged only post-7000 BCE in W Anatolia through eastern migration (Figure 3) and is unlikely to have existed in Greece pre-7000 BCE, strongly supporting Anatolian demographic origins for the European Neolithic expansion (Figures S21-22).

The question then becomes the exact sources of those Anatolian populations and the routes they used. Genomes from >6000 BCE from Greece are limited to a few sites, Nea Nikomedeia and Revenia in N Greece (Hofmanová et al., 2016; Marchi et al., 2022) and Aposelemis in Crete (Skourtanioti et al., 2023) (Figure 1). The latter was produced using capture technology and we could not model it in qpAdm (Methods), while qpAdm models of the former two sites were overall similar to those of W Anatolia, as noted earlier (Marchi et al., 2022) (Figure 3 and Figures S16-17). Outgroup-f_3_ and f_4_-statistics also indicated higher similarity between Greece and W Anatolia relative to C Anatolia (Figure 4; Figures S13, S18, S21). To increase our resolution for studying demographic connections, we imputed >0.25x shotgun genomes and >1x capture genomes and used this data to study haplotype sharing patterns (Methods). Using identical-by-descent (IBD) haplotype-sharing with ancIBD (Ringbauer et al., 2023), we found shared segments of size 8-12 cM representing distant relationships (e.g. >20 generations apart) between pre-6000 BCE N Greece and both NW and CW Anatolia (Figure 5, Figure S22). Clustering based on haplotype-sharing statistics, in turn, suggested strongest affinities to NW and SW Anatolia (Figures S10-11, S14, S23). These observations do not pinpoint a single Anatolian region as the origin of westward expansion into Greece; we either lack the resolution to identify it, or the origins may have been dispersed, including both pan-Aegean sea travel and possible land routes through Thrace.

**Figure 5:**
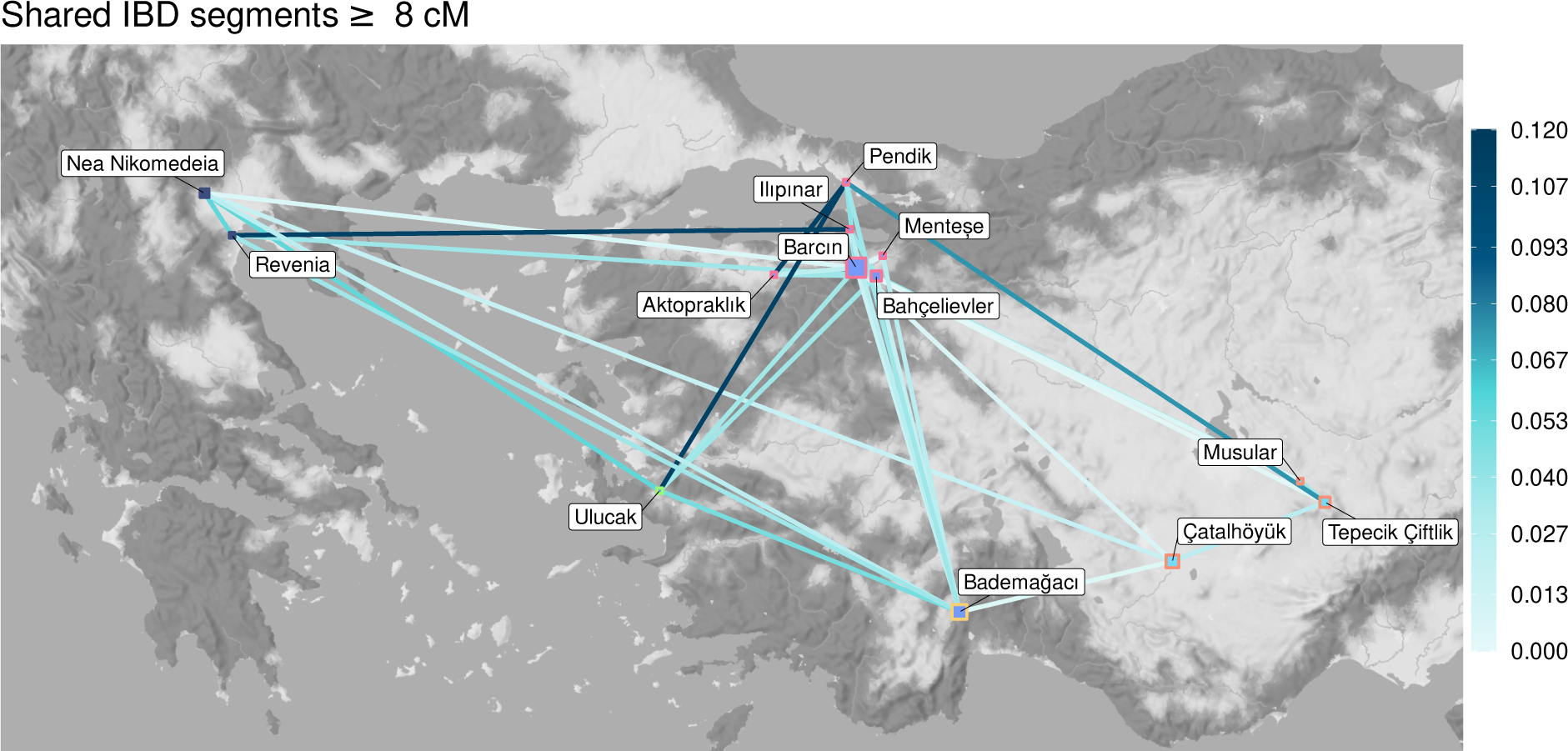
IBD-sharing network across the Aegean showing distant genetic relatedness. The analysis was performed using imputed ancient genomes from the study region and includes ≥8 cM segments (the majority of these were 8-12 cM, and the only IBD-segments >16cM were found between Nea Nikeamedia-Revenia in Greece, and between Aktopraklık-Bahçelievler in NW Anatolia). The colours shown in the key indicate the strength of connections between pairs of genomes in any two sites. We calculate the strength given the number of comparisons and the maximum IBD sharing observed in the dataset. For example, if regions X and Y are represented by 3 and 5 genomes, if any two genomes share a maximum of 4 segments in the full dataset, and if across the 15 X-Y comparisons there are 7 segments in total shared, the X-Y connection strength is estimated as 7/(4×15) = 0.12 (see also Table S5 and Figure S22).

The incomers must have encountered Mesolithic communities on the Aegean islands and in mainland Greece, with whom they were likely already in contact given the presence of Greek (Melos) obsidian in Anatolia by the early 7th mil. BCE and possibly earlier (Gemici et al., 2022; Guilbeau et al., 2019; Horejs et al., 2015). These Mesolithic communities have not yet been genetically sampled and whether they were closer to the pre-Neolithic W Anatolian or the Balkan Mesolithic (“Balkan HG”, i.e. Iron Gates) gene pool is unknown. In either case, any admixture between local groups into incoming Neolithic communities from Anatolia should be detectable in Greece, similar to the admixture signatures identified in W Anatolia. However, using the W Anatolian genetic profile as a reference, we did not detect any additional Balkan HG-related or Girmeler-related admixture signal in Greece. This was true both in qpAdm models and f_4_-tests, and irrespective of whether we used pre-6000 BCE or post-6000 BCE genomes (Figures 3-4, Figures S24).

We asked whether we might be missing low-level admixture signals in our data from Greece. Using simulations of Balkan HG and Anatolian/Balkan Neolithic genomes and by limiting the analysis to the same data type (shotgun or 1240k SNP capture), we found that ≥10% of Balkan HG admixture can be confidently detected in qpAdm models (Figures S2-3). In addition, we could construct reliable qpAdm models for genomic data from 30 published European Neolithic sites across SE, S, and C Europe (6000-2000 BCE) that indicated admixture between an early Neolithic Greece-related source (Greece >6000 BCE) and a Balkan HG-related source, with 4-38% contribution from the latter (Figure S25), a finding also supported by f_4_-tests (Figures S26-27). In contrast, genomes from four later Neolithic (post-6000 BCE) sites in Greece were simply modelled as early Neolithic Greece (>6000 BCE) without additional admixture (Figure 3, Figure S24), and showed high similarity to pre-6000 BCE genomes from Greece and W Anatolia (Figure S28).

The lack of indigenous (Mesolithic-related) admixture signatures in Greece is even more surprising given genetic evidence for early genetic interactions between incoming Neolithic communities and Balkan Mesolithic groups. This is seen in genetically W Anatolia/Greece N-related individuals buried in the Balkan Iron Gates Mesolithic site at Lepenski Vir between c.6200-5900 BCE, and evidence for genetic admixture between Anatolian and local groups in both Lepenski Vir and neighbouring Padina in this period (Brami et al., 2022), where we estimated 24% and 47% Anatolian ancestry in two published genomes (Figure S8C).

A possible explanation for this curious lack of local admixture signatures in Greece could be a large immigrant population swamping the local population in Greece but only small agriculturalist populations expanding across the rest of Europe; this would make admixture signals more visible in the latter. However, studying runs of homozygosity (ROH) or f_3_-based diversity we found no overall elevation in homozygosity in Neolithic genomes from the Balkans and S Europe compared to those in Neolithic genomes from Greece, which would be indicative of a bottleneck in the former (Figures S29-32). The difference between admixture dynamics in Greece and the rest of Europe might therefore be explained by differences in local forager-migrant farmer interaction dynamics, or by subsequent immigration from Anatolia into Greece that diluted local admixture signatures, which would be consistent with IBD-sharing patterns (Figure S22).

### The evolution of material culture uncoupled from demographic dynamics in the Aegean Neolithisation process

Our genetic data support a scenario where the W Anatolian Neolithic arose by both C Anatolian migration and acculturation of indigenous forager groups and their eventual admixture. This appears to align with previous proposals based on the material culture evidence that invoked immigration and admixture with locals (M. Özdoğan, 2013). This, and the fact that sites across W Anatolia were both culturally and genetically heterogeneous, motivated us to investigate whether sociocultural affinities might be correlated with genetic affinity patterns among sites. Such a result would arise if cultural patterns were shaped by mobility and admixture.

A qualitative evaluation indicates little, if any, relationship between sociocultural and genetic similarities among settlements. For instance, CW Anatolian sites (e.g. Ulucak) have predominantly Melos obsidian from Greece (Guilbeau et al. 2019), while obsidian in NW Anatolian sites (e.g. Aktopraklık and Barcın) was mainly of C Anatolian origin (Milic, 2014). Naively, this might be interpreted as Ulucak having inherited indigenous maritime culture/connections while NW Anatolia having more connections to C Anatolia. Genetically, however, Ulucak carries the weakest indigenous ancestry signals in W Anatolia; nor is it connected with the Levant or Cyprus. In NW Anatolian Barcın, both architecture and subsistence data indicate close sociocultural connections to C Anatolia, with little indication of Mesolithic sociocultural presence (Gerritsen & Özbal, 2019) (Figure 4). Meanwhile, in other NW Anatolian sites such as Aktopraklık, Bahçelievler and Pendik, semisubterranean round huts have been considered an indication of Mesolithic influence, and the subsistence patterns of Pendik and neighbouring villages (the “Fikirtepe culture”) interpreted as an amalgamation of Mesolithic and C Anatolian Neolithic traditions (Balcı et al., 2022; Karul, 2017; M. Özdoğan, 2014, 2024). Genetically, however, only in early Aktopraklık do we find strong evidence for local ancestry, while local admixture signatures in “culturally C Anatolian” Barcın are equally prevalent as in “culturally Mesolithic-influenced” Pendik and Bahçelievler (Figure 3).

We then investigated these patterns systematically, calculating correlations between sociocultural and population genetic affinities among sites while controlling for geographic proximity (Methods). For this, we collected and binary coded material culture traits across 89 sites from SW Asia and SE Europe between 9500-5800 calBCE. The information comprised 58 traits including burial and ritual elements, architecture, pottery, lithics, and obsidian sources collected from the literature (Methods; Table S6). Using this data matrix we measured sociocultural distances using Jaccard dissimilarity among all pairs of sites and compared these with their geodesic distances (geographical distances representing the shortest path), separating the data into three periods: 9500-8500 BCE (early PPN), 8500-7000 BCE (PPN), and 7000-5800 BCE (Pottery Neolithic) (Methods). As expected, sociocultural similarities could be explained by geographical proximity among sites in all periods (Spearman r>0.21, Mantel test p<0.01). Moreover, we saw a decreasing correlation through the Neolithic (from r=0.48 to r=0.21; Figure S33). This may reflect the growing intensity of interregional cultural exchanges in parallel with interregional genetic admixture through the Neolithic (Koptekin et al., 2023).

Next, we limited the dataset to 16 sites from 7000-5800 BCE which had both cultural and genetic data. We again found a positive correlation between geodesic and cultural distances (Spearman r=0.49, p=0.001), as between geodesic and genetic distances (r=0.58, p=0.001). Genetic affinities were also correlated with cultural affinities when tested directly (r=0.26, p=0.033). However, after controlling for the confounding effect of geography, genetic affinities did not explain any significant variation in cultural affinities (p>0.10) (Figure 6). Repeating the analysis using only Aegean and Anatolian sites, testing each trait group separately, or using Girmeler affinity differences as genetic distances did not change the results (Figures S34-36). Admittedly, our analysis does not have high resolution as it combines data from multiple centuries of a settlement’s occupation in single-digit records; the genetic information may also vary as new evidence becomes available. Still, it is worthwhile that both qualitative and quantitative analyses point to the mismatch between genetic background and cultural affinities. We hypothesise that at the regional level, sociocultural affinity patterns may be evolving more rapidly and plastically than genetic affinities captured by ancient DNA data; hence the lack of correspondance.

**Figure 6:**
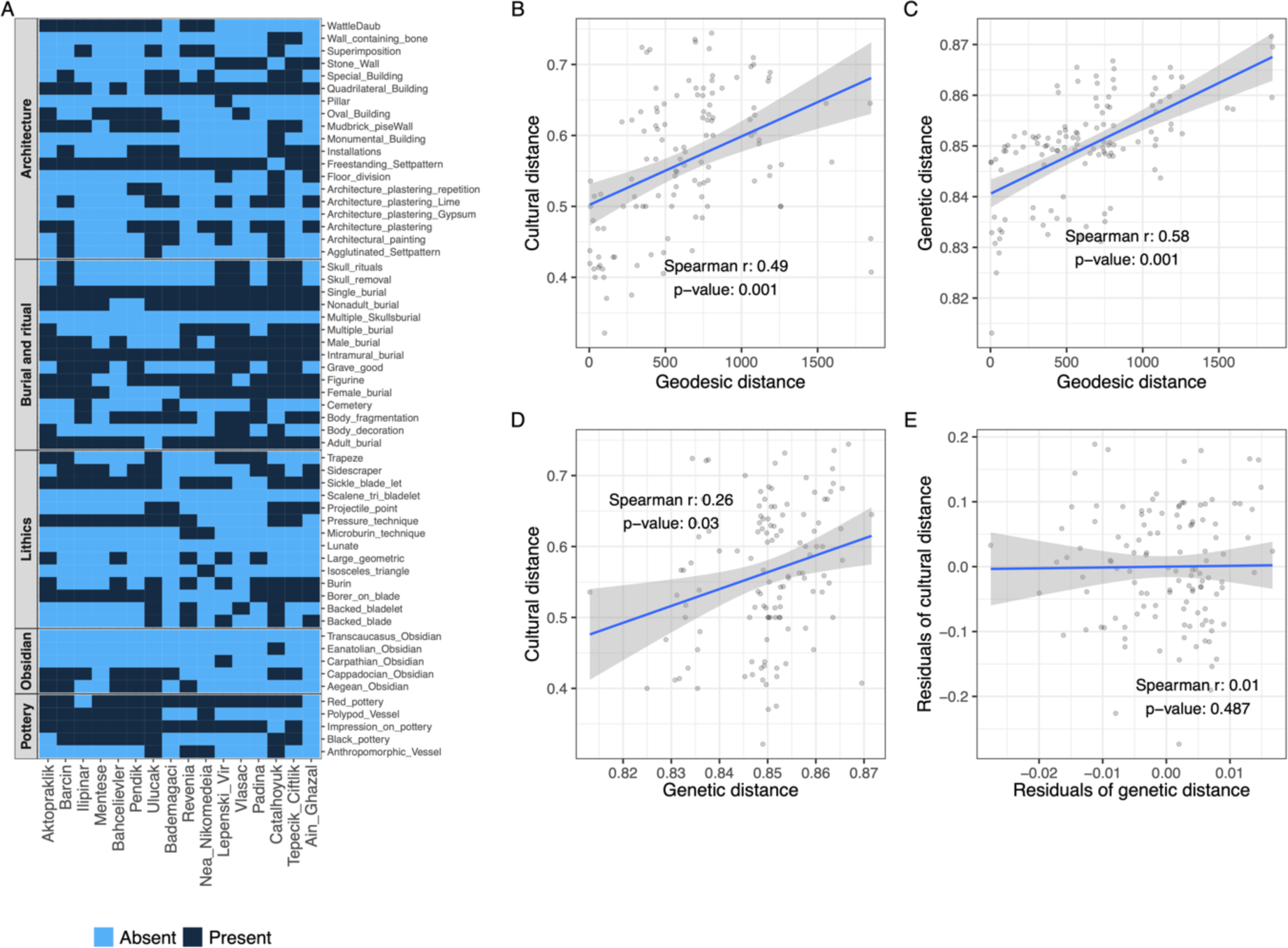
The influence of geographic proximity and genetic similarity on material culture similarities among 7^th^ millennium BCE sites across SW Asia and the Aegean. A: Presence/absence records of 58 material culture traits compiled from the literature for 16 SW Asian/Aegean sites covering 7000-5800 BCE and that have genetic data (see Table S6 for the full dataset). B-D: Correlations between pairwise distances among sites in material culture (Jaccard dissimilarity), geodesic distances (geographic shortest path estimates) and genetic distances (1-f_3_) across the 16 sites (Methods). E: Correlation between the residuals of sociocultural and genetic distances after each was regressed on geodesic distances using linear regression. The Spearman correlation coefficient and the Mantel test p-values are shown inside the panels. Partial Mantel tests between cultural and genetic distances controlling for the effect of geography were also non-significant (r=-0.03, p=0.56) (see also Figures S33-36).

## Conclusion

Our expanded genomic dataset has helped resolve long-standing questions on the earliest westward steps of the Neolithic expansion beyond the Fertile Crescent. We first describe the indigenous gene pool of W Anatolia prior to Neolithisation, represented by the 7800 BCE Girmeler genome, which was closely related to C Anatolians from the PPN period but distinct in its lack of U Mesopotamian ancestry. The presence of PPN-related cultural elements in Girmeler provides another case of cultural interaction among regional communities without major genetic admixture. The fact that we find U Mesopotamia-related admixture during the PPN in C Anatolia but not in Girmeler is also interesting, raising the possibility that eastern gene flow into C Anatolia >8k BCE may have involved the introduction of certain Neolithic elements such as mudbrick technology or animal management practices, which either did not reach Girmeler or were simply not adopted there.

Moving forward in time, we use the Neolithic Aktopraklık genome from c.6600 BCE (Marchi et al. 2020) to infer that the Pınarbaşı/Girmeler-related indigenous foragers persisted in W Anatolia at least between c.13ky-6.6ky BCE. We further infer that Neolithisation in the region involved both large-scale mobility from C Anatolia and cultural adoption by local groups, as well as admixture between the two. The patterns we observe resemble those observed in N Africa (Simões et al., 2023), where the expansion of Neolithic communities appears to have triggered cultural change among locals. Through comparative analysis of material culture traits and ancient genomes for the first time to our knowledge, we test the hypothesis that the observed cultural heterogeneity in regions of Neolithic expansion can be explained by heterogeneity in admixture patterns, but find no support for this. Instead, sociocultural affinity patterns, even in ritual elements such as burial practices (Figure S35), may have been more fluid than genetic admixture processes, at least during the Aegean Neolithisation.

Our results raise new questions. One is the sources of gene flow into W Anatolia before 7000 BCE. A second question is the reason for the apparent lack of indigenous genetic contribution in the Neolithic populations of Greece, which appears unusual given what is observed in the rest of Europe. Another question pertains to the lack of correlations between sociocultural and population genetic similarities at the local level: how universal is this conclusion, and at what levels and historical depth do mobility and admixture cease to shape cultural affinities? Or may correlations become visible only with more refined classifications of material culture assemblages and more comprehensive genetic data? Through further integration of material culture and palaeogenomic data, we are thus starting to disentangle the diverse sources of cultural shifts in prehistoric societies.

## Methods

### Sample preparation

All aDNA samples were processed at the aDNA facilities of the Middle East Technical University and Hacettepe University in Ankara. Before DNA extraction, the surfaces of bones and/or teeth were decontaminated using 0.5% sodium hypochlorite and irradiated with UV-light in a cross-linker. The samples were ground using a SPEX 6770 freezer mill or drilled using a Dremel hand motor to obtain approximately 80-120 mg of fine powder. DNA extraction was performed by following the Dabney protocol (Dabney et al., 2013) with an additional pretreatment step of a brief wash with 1% sodium hypochlorite. Single- or dual-indexed, double-stranded, and blunt-end Illumina sequencing libraries were prepared from 20 ul of DNA extracts using the Meyer and Kircher protocol (Meyer & Kircher, 2010; Kircher et al., 2012). qPCR was performed to determine each library’s optimal number of PCR cycles. Negative controls were included at every step of indexed library preparation. After amplification, the libraries were purified using AMPure XP Beads (Beckman Coulter). Library quality and concentration check was made by Biyoanalyser 2100 (Aqilent). Libraries were mixed in a final pool of equimolar concentrations and sequenced at SciLife, Sweden. The sequencing platform used for producing each library is listed in Table S1.

### Sequence data processing

Residual adapter sequences in raw FASTQ files for each library were removed using “*Adapter Removal (v 2.3.1)*” software (Schubert et al., 2016) with options “*--qualitybase 33 -- gzip --trimns*” and a minimum 11 bp overlap between pairs “-*-collapse --minalignmentlength 11*”. The merged reads were aligned to the human reference genome (*hs37d5*) with “*BWA aln/samse (v 0.7.15)*” (Li & Durbin, 2009) with parameters *“-n 0.01, -o 2*” and seed disabled with “*-l 16500*”. Multiple libraries from the same individual were merged using “*samtools merge (v 1.9)*” (Danecek et al., 2021), and PCR duplicates with identical start and end coordinates were deleted with the code “*FilterUniqueSAMCons.py*” (Kircher, 2012). Reads with >10% mismatches to the human reference genome, mapping quality < 30, and base pair count < 35 were discarded.

Importantly, samples with both UDG and non-UDG libraries were merged after the read-end trimming process (see below). We computed average genome coverage using “*genomeCoverageBed*” implemented in “*bedtools2*” (Quinlan, 2014) tool. This included only reads with mapping quality >30. To eliminate biases, previously reported ancient genomic data were remapped and filtered through the same preprocessing steps (see Table S1).

We also generated trimmed versions of bam files by trimming the ends of reads (a) removing 10 bases at the ends of each read from libraries obtained with double-stranded non-UDG libraries, (b) removing 2 bases at the ends of each read from libraries obtained with double-stranded UDG libraries, c) removing 10 bases from the 5’ end and 2 bases from the 3’ end for single-stranded non-UDG libraries. We used the trimmed bam files for genotyping and mtDNA contamination estimation (see below).

### Contamination estimation and quality control

After removing reads with minimum base quality and mapping quality <30 for each library, we used three ways to determine the authenticity of the remaining reads. First, we examined postmortem deamination patterns generated by cytosine deamination in all samples using “*PMDtools (v 0.60)*” (Skoglund, Northoff, et al., 2014) with the “*--deamination*” parameter. Second, we used “*contamMix (v 1.0-10)*” (Fu et al., 2013), which uses the rate of consensus mitochondrial sequence mismatches. To detect non-endogenous reads *contamMix* computes a contamination probability based on a reference panel of 311 mitochondrial genomes. For this approach, consensus mitochondrial sequences were produced using “*ANGSD (v 0.941*)” (Korneliussen et al., 2014) with the options “*-doFasta 2 -doCounts 1 - minQ 30 -minMapQ 30 -setMinDepth 3 -rMT”.* Finally, for the male individuals, contamination based on the X chromosome was evaluated using *ANGSD* with the command “*angsd -i BAMFILE -r X:5000000 -154900000 -doCounts 1 -iCounts 1 -minMapQ 30 -minQ 30*” for X chromosome-mapped reads. The probability of heterozygosity on the X chromosome was subsequently estimated using the R script “*contamination.R*” and the reference files included in the *ANGSD* package (Korneliussen et al., 2014).

The molecular sex of all samples was established using the *“Ry*” method (Skoglund et al., 2013), which used reads with a minimum base quality and a mapping quality of 30. Mitochondrial DNA haplogroups were determined as follows; first, consensus mitochondrial sequences were created from sequence alignment files using “*ANGSD (v 0.941)*” with parameters “*-doFasta 2 -doCounts 1 -minQ 30 -minMapQ 30 -setMinDepth 3*”, then haplogroups were assigned to individuals using the “*HaploGrep2*” (Weissensteiner et al., 2016) tool. Y chromosome VCF files were created from bam files obtained by extracting reads aligned to the Y chromosome from whole genome alignment data using “*samtools mpileup (v 1.9)*” and “*bcftools call (v 1.9)*” (Danecek et al., 2021). Here, base and map quality scores < 30 were filtered. Then, Y chromosome haplogroups were assigned for all male samples by using “*Yhaplo (v 1.1.2)*” (Poznik, 2016) based on Y-DNA Haplogroup Tree 2019-2020 of ISOGG (https://isogg.org/) (see Table S1).

### Genetic kinship analyses

We ran the software “*READv2*” (Alaçamlı et al., 2024) to estimate close kin up to third-degree relatedness between each pair of individuals from the same population using Dataset 1 (see below). *READv2* calculates the kinship coefficient (θ) between pseudo-haploid genome pairs. Because for Pendik we had only n=2, we computed the θ values for the analysis of the Pendik pair using all unpublished and published shotgun-sequenced genomes from Neolithic NW Anatolia as background (for P0 estimation). We only found one second-degree related pair in Bahçelievler (bbh008 and bbh010) and removed bbh008 from the analysis as it had lower coverage (see Table S1).

### Datasets

We prepared four datasets based on different SNP panels and which were used in different population genetics analyses. We only used autosomal variants in these datasets.

<u>Dataset 1:</u> The 1000 Genomes sub-Saharan African dataset includes 4,771,930 autosomal SNPs ascertained in sub-Saharan African populations in Phase 3 of the 1000 Genomes Project (Koptekin et al., 2023), merged with 300 present-day samples from Mallick et al (Mallick et al., 2016) (downloaded from https://reichdata.hms.harvard.edu/pub/datasets/sgdp/ as “*PLINK format*” on 30 Aug 2020) and ancient individuals from SW Asia and S Europe.

<u>Dataset 2:</u> The 1240K Capture Array dataset includes ancient individuals from SW Asia and S Europe called across a total of 1,151,161 autosomal SNPs (Mathieson et al., 2015)

<u>Dataset 3:</u> This dataset includes the Human Origins SNP Array genotypes (2,583 present-day individuals genotyped on the Affymetrix Human Origins Array) merged with ancient individuals from SW Asia and S Europe called across a total of 616,938 autosomal SNPs (Lazaridis et al., 2014, 2016).

Dataset 4: This dataset includes imputed genomes, created using all shotgun-produced ancient genomes with >0.25x coverage and capture-produced ancient genomes with >1x coverage (n=187), and using the 1000 Genome Phase3 autosomal dataset as reference panel (see Genotype imputation).

We employed the Human Origins SNP Array dataset (Dataset 3) to conduct PCA and *ADMIXTURE* analyses; the 1240K Capture Array dataset (Dataset 2) to estimate Runs of Homozygosity through *hapROH* and estimate admixture time by using *DATES*; a subset of imputed data containing 1240K SNPs used for haplotype-based analysis, ancIBD and FineSTRUCTURE, and the 1000 Genomes sub-Saharan African dataset (Dataset 1) for all remaining analyses.

### Ancient genome sample selection

In our dataset, we included all published Late Upper Pleistocene / Early Holocene (25,000-6000 BCE) palaeogenomes (as of May 2024) from Anatolia (Feldman et al., 2019; Hofmanová et al., 2016, 2022; Kılınç et al., 2016; Koptekin et al., 2023; Lazaridis et al., 2016, 2022; Marchi et al., 2022; Mathieson et al., 2015; Yaka et al., 2021), Greece (Hofmanová et al., 2016; Lazaridis et al., 2017; Marchi et al., 2022; Mathieson et al., 2018; Skourtanioti et al., 2023), the Balkans (Allentoft et al., 2024; Freilich et al., 2021; González-Fortes et al., 2017; Lazaridis et al., 2022; Marchi et al., 2022; Mathieson et al., 2018), the Levant (Feldman et al., 2019; Lazaridis et al., 2016, 2022; Wang et al., 2023), S Caucasus (Allentoft et al., 2024; Jones et al., 2015; Lazaridis et al., 2022; Skourtanioti et al., 2020) and the Zagros/S Caspian regions (Allentoft et al., 2024; Broushaki et al., 2016; Gallego-Llorente et al., 2016; Lazaridis et al., 2016, 2022; Narasimhan et al., 2019).

We further chose representative genomes from Neolithic S Europe (Allentoft et al., 2024; Antonio et al., 2019; Brunel et al., 2020; González-Fortes et al., 2019; Olalde et al., 2015; Valdiosera et al., 2018), as well as European Hunter-Gatherer (HG)-related genomes (de Barros Damgaard et al., 2018; Gamba et al., 2014; González-Fortes et al., 2017; Jones et al., 2015; Lazaridis et al., 2014; Mathieson et al., 2015; Olalde et al., 2014).

In addition to these regions, we included *Ust_Ishim* (Fu et al., 2014)*, Kostenki14* (Seguin-Orlando et al., 2014)*, AfontovaGora3* (Fu et al., 2016)*, MA1* (Raghavan et al., 2014) genomes as well as the N African *Morocco_Taforalt* genomes (van de Loosdrecht et al., 2018) for use as the “right populations (outgroups)” in qpAdm modelling.

If there were either first- or second-degree related individuals from the same site, we retained the highest coverage genome and excluded the rest from the dataset. We labelled all removed relatives with the “*_rel*” suffix in Table S2.

### Pseudo-haploid genotyping

We generated pseudo-haploid datasets by randomly selecting one allele for each targeted SNP position using the genotype caller “*pileupCaller (version 1.5.3.1)*” (https://github.com/stschiff/sequenceTools) on output from “*samtools mpileup*” (with base quality > 30 and MAPQ > 30) (Li, 2011) on trimmed bam files (see above). Genotyping was performed independently for each of the three SNP panels, thus creating three datasets.

### Genotype imputation

We used the imputation and phasing tool “*GLIMPSE2*” (Rubinacci et al., 2023). We first generated genotype probabilities using the “*bcftools mpileup (v 1.18)*” command with parameters *“-I -E -a ‘FORMAT/DP’ -T -q 30 -Q 30”*’, followed by ‘*bcftools call -Aim -C alleles*’ (Li, 2011). As reference panel we used the 1000 Genomes Phase 3 Dataset (Auton et al., 2015) generated following the steps outlined in the *GLIMPSE2* tutorial (step 2.2 in https://odelaneau.github.io/GLIMPSE/docs/tutorials/getting_started/#2-reference-panel-preparation). We similarly followed this GLIMPSE2 tutorial in the next steps. We generated imputed regions using “*GLIMPSE2_chunk”*. Then, the reference panel was converted to binary format using the “*GLIMPSE2_split_reference”* tool with default parameters. Input data for this step included the reference panel of haplotypes, the genetic map, and the imputed regions created in previous steps. We then imputed each chunk with “*GLIMPSE2_phase”* with default parameters, employing the binary reference panel obtained in the preceding step. Finally, the imputed chunks belonging to the same chromosome were merged with the “*GLIMPSE2_ligate”* tool.

We imputed all shotgun-produced ancient genomes with >0.25x coverage and all capture-produced ancient genomes with >1x coverage (see Table S2).

### Analyses of population structure

We performed principal components analysis by using the “*smartpca (v 18140)*” program of “*EIGENSOFT (v 7.2.0)*” (Patterson et al., 2006) with “*lsqproject:YES, numoutlieriter: 0*” parameters to construct the components of present-day West Eurasian populations (49 populations, 946 individuals) from the Human Origins SNP Array dataset (Dataset 3). Ancient individuals were projected onto the first two principal components of present-day genetic variance (see Figure S5).

We performed multidimensional scaling with the “cmdscale” function of R on a pairwise genetic distance matrix. Distances among all pairs of individuals were estimated using the measure 1-f_3_ (see below). Dataset 1 was used in this analysis (see Figure 2 and Figure S4). We filtered the genome set to avoid any pairs that did not share <2000 overlapping SNPs, to minimize noise in the estimates.

### Allele frequency correlation statistics

We calculated outgroup-f_3_- and f_4_-statistics using “*AdmixTools (version 7.0.2)*” (Patterson et al., 2012) using Dataset 1.

Outgroup-f_3_ statistics representing the genome-wide similarity between a pair of populations was calculated for all pairs with the “*qp3Pop (v 651)*” algorithm, the 1000 Genomes phase3 Yoruba population (n=108) serving as an outgroup. Genetic distances between pairs of populations were measured using the 1-f_3_-statistic in Figure S29. A threshold of ≥2,000 overlapping SNPs was used for reporting f_3_-based statistics.

To estimate gene flow between Population X and Population Y, f_4_-statistics were calculated with the “*qpDstat (v 980)*” algorithm. Tests of the form f_4_(Test, Outgroup; PopX, PopY) were conducted again using Yoruba as an outgroup, with the “f4mode: YES” option. A threshold of more than 10,000 overlapping SNPs was used for reporting f_4_-test calculations (see Figure 4; Figures S7, S9, S15, S21, S24, S26, S27 and Table S3).

### Model-based clustering

We performed unsupervised model-based cluster analysis using “*ADMIXTURE (v 1.3.0)*” (Alexander et al., 2009). To do this, we first selected Western Eurasian present-day populations (49 populations, 946 individuals) from Dataset 3. Then we pruned SNPs for linkage disequilibrium, and filtered out sites with a minor allele frequency (MAF) less than 5% in “*PLINK (v 1.9)*’ (Chang et al., 2015), using the parameters “*--indep-pairwise 200 25 0.4*” and “-*-maf 0.05*,” retaining 137,137 SNPs. After pruning and filtering, we combined W Eurasian present-day populations with ancient individuals (n =337) and estimated the average genomic ancestry proportions for the 946 individuals. Clustering was performed from K = 2 to K = 5 using default fivefold cross-validation (“--cv = 5”) and 10 replicate runs with different random seeds. The cross-validation procedure of *ADMIXTURE* determined the optimal K value. The LargeKGreedy algorithm of “*CLUMPP*” (Jakobsson & Rosenberg, 2007) was employed to identify common signals across each independent run (see Figure S6).

### Coancestry estimation with FineSTRUCTURE

We used *FineSTRUCTURE v4* (Lawson et al., 2012) for clustering analyses based on cumulative lengths of shared haplotypes between individuals. Here, we used a subset of imputed data containing 1240K SNPs, and included individuals represented by either >0.5x coverage shotgun genomes or >2x coverage capture genomes (see “Genotype imputation”, Table S2). Prior to clustering, we performed *ChromoPainter* analyses from the same software and calculated chunk counts and lengths for each pair of individuals. In the first run, we estimated switch (“*-n*”) and emission (“*-M*”) rates on chromosomes 1,3,5,9 of 10 individuals. We then performed *ChromoPainter* on all autosomal chromosomes with pre-calculated parameters. Using *FineSTRUCTURE*, we clustered individuals by analyzing the number of shared haplotypes. The number of burn-in iterations (“*-x*”), the number of sample iterations (“*-y*”) and the thin interval (“*-z*”) for the *mcmc* run was 2,000,000, 100,000 and 10,000 respectively (see Figure S10).

We also calculated haplotype-sharing statistics (HSS) by utilizing cumulative lengths of shared autosomal haplotypes as explained in (Flegontov et al., 2019) (see Figures S11, S14, S22 and S28).

### Ancestry proportion estimation with qpAdm

We estimated ancestry proportions of source genomes in target genomes using “*qpAdm (v 1520)*” implemented in “*AdmixTools (v 7.0.2)*”, with “*allsnps: YES, details:YES*” parameters. For all runs, we used a base set of “Right” populations (outgroups) composed of “*Mbuti, Han, Papuan, Mixe, Ust_Ishim, Kostenki14, MA1, WHG, Levant_RaqefetCave_HG_CP, Morocco_Taforalt_CP, AfontovaGora3, Iran_N*” (see Figure 3; Figures S8, S16, S17, S19, S20, S25 and Table S4).

We generated input files for qpAdm analysis that included all combinations of target, source and right populations by using “*qpAdm-wrapper (v 1.0.1)*” (https://github.com/dkoptekin/qpAdm-wrapper).

### Runs of homozygosity (ROH)

ROH were called with *PLINK* (Chang et al., 2015) on shotgun genomes with >5x coverage (n=31). We first called diploid genotypes for Dataset 1 SNPs using “*bcftools mpileup/call (v 1.9)*” with “*-B --ignore-RG -q 30 -Q 30 -R*” parameters (Li, 2011). Then, ROH were identified using by *PLINK (v 1.9)* (Chang et al., 2015) with options “*--homozyg, --homozyg-snp 30, -- homozyg-kb 500, --homozyg-density 30, --homozyg-window-snp 30, --homozyg-gap 1000, -- homozyg-window-het 3, --homozyg-window-missing 5, --homozyg-window-threshold 0.05*”. To avoid possible bias introduced due to different ranges of genome coverage, we also estimated ROH with the same individuals after downsampling all of them to 5x coverage using “*Picard DownsampleSam (v 2.26.2)*” following the same steps (see Figures S30-31).

We additionally estimated ROH using “*hapROH (v 0.64)*” (Ringbauer et al., 2021) with default parameters that were optimized for 1240K capture SNPs. The default genetic map of hapROH and 5,008 global haplotypes from the 1000 Genomes Project were used. We used 186 of the ancient genomes, which were covered by at least 300K SNPs of the 1240K Capture Array dataset (Dataset 2) (see Figure S32).

### Detecting Identical-by-Descent (IBD) segments

We estimated IBD haplotype-sharing analysis with “*ancIBD (v 0.6)*” (Ringbauer et al., 2023) that estimates long haploid blocks along the genomes of two individuals that are identical by descent (IBD). We used a subset of imputed data containing 1240K SNPs, and included individuals with >0.25x shotgun genomes and >1x capture genomes (see Genotype Imputation). We used the “*hapBLOCK_chroms*” function of ancIBD with default parameters to estimate shared blocks of more than 8 cM for each chromosome. We filtered segments with <220 SNPs per cM, following the original publication.

We then calculated IBD-sharing frequency among genomes from different archaeological sites. We thus produced IBD-sharing networks for sharing ≥8 cM segments. We summed the total shared IBD segments among sites. Since sample sizes were heterogeneous among sites, we normalized the total number of shared IBD segments by dividing them by the maximum number of shared IBD segments among pairs of individuals of the same length range (i.e. by normalizing sharing). We estimated IBD-sharing frequency between a pair of sites by dividing the proportion of IBD-sharing calculated among sites by the total number of possible pairs (e.g. between two different sites of n1 and n2 individuals there would be n1*n2 comparisons) (see Figure 5).

We also calculated IBD-sharing frequency among sites to produce the IBD-sharing heatmaps using 8-12 cM, 12-16 cM, 16-20 cM and >20 cM segments from all chromosomes following (Ringbauer et al., 2023). Here we summed the total shared IBD segments among regions and divided them by the total number of possible pairs (e.i. between two different sites of n1 and n2 individuals there are n1*n2 comparisons) (see Figure S22).

### Estimating the time of admixture

To estimate admixture dates between putative ancestral sources for W Anatolian genomes, we used “*DATES (v 3600)*” (Chintalapati et al., 2022) with default parameters. We considered models feasible if they produced a positive mean of less than 200 generations, a normalized root-mean-square deviation (NRMSD) below 1, and a Z-score above 2. A generation length of 28 years was assumed. Dataset 2 was used in the analysis.

We thus ran the algorithm to estimate the dates of possible Balkan and Levant-related admixture that produced W Anatolian local genomes, and the dates of possible W Anatolian local (Girmeler/Aktopraklık) and C Anatolian (post-7500 BCE) admixture that produced the W Anatolian Neolithic genomes. However, none of the models yielded feasible results (see Table S4).

### Generating artificial admixed individuals with genoMIX

To generate artificial admixed individuals from empirical data we used a new algorithm we present here, “*genoMIX*” (https://github.com/dkoptekin/genoMIX). *genoMIX* takes a YAML file as input and mixes the given source populations in specific (user-selected) proportions using a dataset in Eigenstrat format. To do this, first, the tool splits the “input.snp” file into chunks with a fixed number of SNPs in each chunk. It then creates an artificial admixed genome by randomly selecting chunks from one individual of given source populations based on the chosen admixture proportions. The tool can generate admixed individuals as a 2-, 3- and/or 4-way mixture using parameters specified in the input YAML file.

Here, we generated artificial admixed individuals to evaluate the performance of f_4_-tests and qpAdm models using different data types. We mixed Anatolian Barcın Neolithic individuals (I0707, I0708, I0709) with Balkan Iron Gates hunter-gatherers (I4873, I4874, I4877) in proportions of 5%, 10%, 15%, 20%, 25%, 30%, 35%, 40%, 45%, and 50%, using either shotgun-sequenced or 1240K-captured genomes and using 5,000 SNP-chunks.

We then performed f_4_-tests in the form of *f_4_(Yoruba, Balkan HG; Barcın N, Admixed-genome)*, employing different data types (capture vs. shotgun) for Balkan HG and Barcın N each time. Then, we applied qpAdm to model the admixed genomes as a two-way mixture of Balkan HG and Barcın N, again using various data type combinations for Balkan HG and Barcın N. We used the same right populations for qpAdm as we used with real data (see above). Results are shown in Figure S2.

We also generated artificial admixed individuals using Balkan N individuals (I1131, I4877, I4880) instead of Barcın Neolithic, as it is known that Balkan N genomes contain some degree of Balkan HG ancestry, to examine how this would affect the results. Results are shown in Figure S3.

Note that when performing f_4_ and qpAdm analyses, we used different genomes as reference populations than those used to generate the admixed individuals.

### Material culture dataset compilation

To investigate the correlations between the genetic and cultural affinities among sites, we collected and binary coded (present (1)/absent (0)) material culture traits across 89 sites from SW Asia and SE Europe between 9500-5800 calBCE (Table S6). The information was collected using various academic literature sources, including indexed journals, Google Scholar searches, as well as local publications (such as the yearly published results of Archaeological Excavations in Turkey). Our preliminary evaluation of these literature sources indicated that the data on architecture, pottery, lithics, obsidian and ritual elements were more systematically recorded than the subsistence and environmental data. Therefore, for the purposes of this study, we decided to concentrate on five major categories which comprised 58 traits (Figure 6A). These traits were selected in relation to the description of general typological features (such as circular or rectangular architecture, use of plaster, source of obsidian, lithic techno-typology, shape, colour and decoration on pottery, etc.) and were recorded in a spreadsheet, using binary coding of presence or absence of each element, for each site.

We note that the coding of “absence” does not necessarily mean that those material culture elements were actually/fully “absent” at these sites; rather, the excavators, given the excavation scale or the site function, may not have observed them. Nevertheless, if such instances of apparent absence (or equally, presence) were shared by other sites within a similar time and space context, then a pattern can be assumed.

Also, based on our understanding of the major transformations in material culture, architecture, and subsistence patterns, already defined in the literature, we divided the long-time span of our focus into 3 periods; these are represented by a total of 89 sites (Table S6). If a site was occupied for a long span of time, then this site’s information was included for multiple time periods, however, only the data pertaining to that time period as attested from respective layers of the site was recorded for different time periods.

### Quantitative analysis of material culture

We computed pairwise Jaccard dissimilarity among the archaeological sites on the binary coded traits using the “*vegdist*” function of the *“vegan”* (Oksanen et al., 2022) package in *R*. The Jaccard indices were compared with pairwise geographic (geodesic distance between two sites calculated using the “geodist” R package) and genetic distances (1-outgroup-f_3_), as well as pairwise differences among sites in Girmeler ancestral proportion estimates. We measured the correlations between distance matrices using the Mantel test with the “*mantel*” function of the same package with default settings (1000 iterations). We used the non-parametric Spearman correlation to avoid the effect of possible outliers. We further studied the correlation between cultural and genetic distances while controlling the effect of geographic distance on both, a) using the “mantel.partial” function of the same package as *“mantel.partial(cultural_distanceMatrix, genetic_distanceMatrix, geodesic_distanceMatrix, method=“s”, permutations=999)”*, and b) by computing residuals from regressions of culture on geography and genetics on geography using the “*lm*” function in R, and then calculating the correlation between the residuals using the Mantel test (see Figure 6 and Figures S33-36).

### Visualisation

We produced all figures in *R* (R Core Team, 2023) after reading and/or modifying data using “*tidyverse*” (Wickham et al., 2019), “*reshape2*” (Wickham, 2007) and “*gsheet*” (Conway, 2020) packages. All figures produced by using “*ggplot2*” (Wickham, 2016) and its extension packages such as “*ggmap*” (Kahle & Wickham, 2013), “*ggcorrplot*” (Kassambara, 2023a), “*ggh4x*” (Brand, 2024), “*PieGlyph*” (Vishwakarma et al., 2023), “*ggpubr*” (Kassambara, 2023b), and “*ggtree*” (Yu, 2020). The multiple panel figures are combined by using the “*patchwork*” (Pedersen, 2024) package.

### Data and code availability

The raw FASTQ files and aligned sequence data (BAM format, without filtering for mapping quality) reported in this paper can be accessed and downloaded from the European Nucleotide Archive (ENA) under the following study accession number: PRJEBNNNN.

## Acknowledgments

We thank all members of the CompEvo (METU) group, Eva-Maria Geigl, Pavlos Pavlidis, Stefanos Papadadonakis, Gözde Atağ and Yetkin Alıcı for helpful discussions, Ali Akbaba for assistance in sampling, Ian Hodder for allowing access to Çatalhöyük material, Swapan ‘Shop’ Mallick and David Reich for sharing Allen Ancient Genome Diversity Project shotgun genomes, and Joachim Burger and Jens Blöcher for sharing neutralome data from Aktopraklık and Barcın.

This work was supported by the H2020 ERC Consolidator grant (no. 772390 NEOGENE to M.S); the H2020-WIDESPREAD-05-2020 TWINNING grant (no. 952317 NEOMATRIX to M.S.); Wenner-Gren Foundation Dissertation Fieldwork grant (no. 9573 to D. Koptekin), EMBO Scientific Exchange grant (no. 8883 to D. Koptekin), and Swiss National Science Foundation (SNSF) project grant (no. PCEGP3_181251 to A.S.M). CJK‘s participation in this research benefited from the scientific framework of the University of Bordeaux’s IdEx (Initiative d’Excellence) ‘Investments for the Future’ program/GPR (Grands Programmes de Recherche) ‘Human Past’.

Computations were performed on NEOGENE Clusters at Middle East Technical University.

## Supplementary Information

Supplemental Document includes descriptions of archaeological sites and supplemental figures (Figures S1-36)

**Table S1:** Sequencing data information for newly generated ancient individuals and kinship results. The “Date (BCE/CE)” column shows either calibrated C14 dates directly obtained from the samples (with the prefix “cal”), or approximate dates based on archaeological content. Sheet 1 shows results for libraries merged per individual, Sheet 2 shows results for single libraries, and Sheet 2 shows READv2 results for kinship estimates.

**Table S2:** Information on all individuals used in the analyses. Abbreviations are as follows: HG, Hunter-Gatherer; N, Neolithic; CP, 1240K captured data; NC, neutralome captured data; C, Central; NW, Northwest; CW, Centralwest; SW, Southwest. Relative and outlier individuals of the site labelled with the “*_rel*” and “_o” suffix respectively.

**Table S3:** Results of f_4_-tests.

**Table S4:** Results of qpAdm models

**Table S5:** ancIBD analysis results

**Table S6:** Material culture dataset. The table contains presence/absence information on 58 cultural traits across SW Asian / SE European archaeological sites across three Early Holocene time periods, the coordinates of the sites, and the literature used.

