## Supplementary Information for "Out-of-Anatolia: cultural and genetic interactions during the Neolithic expansion in the Aegean"

This file contains “Description of archaeological sites and archaeological material” and Supplemental Figures S1-36

**Description of archaeological sites and archaeological material**

In the table below we summarise information on the West Anatolian Early Holocene settlements included in this study, their archaeological layers, and relevant literature. We then provide descriptions of those archaeological sites from which we produced genomic data..

| **Sites** | **Layers (Dates)** | **Dates of genetically sampled individuals (C14, 2σ)** | **Literature** |
|---|---|---|---|
| **Girmeler** | **Late Pleistosene deposits:**<br>12,000 - 10,600 BCE<br><br>**Early Holocene deposits:**<br>9200 - 8800 BCE<br><br>**Late 9th-8th millennium deposits:**<br>8200 - 7600 BCE | 7738 - 7597 calBCE | (Erdoğu, 2022; Takaoğlu et al., 2014) |
| **Aktopraklık** | **Layer Aktopraklık C:**<br>c. 6600-6000 calBCE<br><br>**Individual AKT16**<br>6658-6578 calBCE | 6658-6578 calBCE<br>6498-6412 calBCE<br>6472-6383 calBCE<br>5632-5525 calBCE | (Budd et al., 2013; Karul & Avci, 2011) |
| **Barcın** | **Phases VIa-VIb:**<br>c. 6200-6000 calBCE<br><br>**Phases VId2-VId3 and VIc:**<br>c. 6400-6200 calBCE<br><br>**Phases VIe and VId1:**<br>c. 6600-6400 calBCE | 6434-6255 calBCE<br>6419-6238 calBCE<br>6419-6246 calBCE<br>6387-6111 calBCE<br>6386-6228 calBCE<br>6381-6088 calBCE<br>6228-6081 calBCE<br>6225-6065 calBCE<br>6224-6074 calBCE<br>6424-6233 calBCE<br>6223-6034 calBCE<br>6221-6028 calBCE<br>6217-6007 calBCE | (Gerritsen & Özbal, 2019; H. Özbal et al., 2024; R. Özbal & Gerritsen, 2019) |
| **Bahçelievler** | **Layer 6:**<br>6600-6400 calBCE<br><br>**Layer 5**:<br>6400-6200 calBCE | 6612-6471 calBCE<br>6238-6391 calBCE | (Fidan et al., 2022) |

| | | | |
|---|---|---|---|
| **Ilıpınar** | **Layer VI:**<br>5600-5460 calBCE<br><br>**Layer VII:**<br>5600 calBCE<br><br>**Layer VIII:**<br>5700-5600 calBCE<br><br>**Layer IX:**<br>5700 calBCE<br><br>**Layer X:**<br>5950/5900-5700 calBCE | N/A | (Roodenberg, 1993, 2013; Thissen & Roodenberg, 1995) |
| **Menteşe** | **Stratum 1:**<br>5800-5600 calBCE<br><br>**Stratum 2:**<br>5950-5800 calBCE<br><br>**Basal Menteşe (Stratum 3 Upper):**<br>6200-5950 calBCE<br><br>**Basal Menteşe (Stratum 3 Middle)**<br>6300-6200 calBCE<br><br>**Basal Menteşe (Stratum 3 Lower)**<br>6400-6400 calBCE | 6008-5835 calBCE | (Alpaslan-Roodenberg & Maat, 1999; Roodenberg, 1999; Roodenberg et al., 2003) |
| **Pendik** | **Archaic Fikirtepe and Classical Fikirtepe phases**<br>late 7$^{th}$ millennium | 6387-6230 calBCE | (Harmankaya, 1983; Kızıltan, 2013, 2022; Özdoğan, 1983, 2001, 2013, 2014; Pasinli et al., 1994; Ulaş, 2020) |
| **Bademağacı** | **ENII/4-ENII/2**<br>6500/6400-6200 calBCE<br><br>**ENI/9 - ENI/5**<br>7000-6800/6500 calBCE | 6389-6237 calBCE<br>6381-6222 calBCE | (Duru, 2012; Duru & Umurtak, 2019) |

**Girmeler**

Girmeler (Erdogu et al., 2021; Korkut, 2016; Takaoğlu et al., 2014) is currently the only known excavated settlement in the Aegean coastal region of Anatolia that predates the establishment of fully sedentary agricultural villages. It is situated atop a small limestone hill in the Eşen Valley, within the bounds of the ancient Lycian city of Tlos. The site comprises two caves with extensive galleries and an adjacent mound. Nearby lies a natural hot spring.

In the 1980s, approximately seven meters of the mound's height was removed to level the area between the hot spring and the cave entrances to construct a thermal centre. The earliest archaeological evidence of human activity at Girmeler dates to the Epipalaeolithic period, roughly between 12,000 and 10,600 BCE, with transitional layers to the early Holocene evident around 9200-8800 BCE.

Three distinct trenches at Girmeler have yielded archaeological evidence of early Holocene habitation. Trenches A and C were excavated in 2013, and trench D in 2020. Radiocarbon dating of samples from these trenches indicates they date to 8200–7600 BCE, corresponding to Girmeler's early Holocene strata. Preliminary excavations have uncovered substantial evidence of pre-pottery Neolithic activities, indicative of a semi-sedentary population engaged in intensive hunting, wild plant gathering, and nascent agricultural experimentation. Trench A revealed a section of a structure with a lime-plastered floor composed of small pebbles embedded in lime. Postholes at the floor's corners and surrounding building debris suggest the structure once supported a wattle-and-daub superstructure. The structure's circular sunken mud-plastered basins and hearths appear to have been regularly renewed, indicating prolonged and consistent use.

The chipped stone industry at Girmeler was mainly a flake-based one but aimed at producing blades. Blades are scarce or virtually absent in the Mesolithic Aegean Island assemblages, precluding direct comparisons with Girmeler's percussion blade production. Chaff barley, likely wild, is a prevalent find. Glume bases of glume wheat, possibly domesticated, are also found, albeit infrequently and in fragmented form. One glume base has been classified as a tetraploid wheat (*Triticum turgidum/timopheevii* type). The discovery of large grinding stones, domestic wheats, wild barley, and at least one sickle blade at Girmeler suggests the presence of food production and initial steps toward agriculture.

Faunal analysis has identified bones from various wild species, including goats, roe deer, fallow deer, wild boar, and hare, as well as caracal, red fox, marten, and hedgehog. The caprine remains are believed to be exclusively goats, as no sheep specimens have been definitively identified. No evidence of domesticated or managed animals has been found.

<u>Description of burial practices and skeletal material</u>
In front of the cave entrances, three graves have been uncovered. The most intact grave from this period contained the remains of an adult woman positioned in a flexed posture with a large stone near her head (Erdogu et al., 2021; Korkut, 2016; Takaoğlu et al., 2014).

DNA analysis was conducted on this individual [gir001 (SK1)]. Beyond the mound, five additional skeletal remains associated with painted Terrazzo floors, dating to the early 7th millennium BC, have been found. Unfortunately, their poor condition precluded DNA analysis from these.

**Ulucak Höyük**

Ulucak Höyük, situated 25 km east of İzmir in Central-West Turkey, is a modest-sized mound spanning approximately 1 hectare with an 11-meter stratigraphic sequence. The site's Neolithic presence, spanning Levels VI to IV and dating from 6850/30 to 5670 calBCE, marks it as one of the earliest known locations for agriculture and four-tiered animal husbandry [including domestic pigs, that are absent in sites such as Çatalhöyük] in western Anatolia (Çakırlar, 2012; Çevik & Abay, 2016; Çevik & Erdoğu, 2020; Çilingiroğlu et al., 2012). The initial phase (Level VI: 6850-6500 cal BCE) did not exhibit residential structures; instead, the rectangular buildings with lime-plastered and red-painted floors and walls likely served communal purposes. Over time, construction methods and house layouts evolved. Early single-room, rectangular dwellings with wattle-and-daub or mud slab walls gave way to robust structures with sun-dried mud brick walls atop stone foundations in the early sixth millennium BCE. Later phases introduced internal partitions and enclosed courtyards as standard features in some homes. The presence of individual cooking, food processing, and storage facilities within each building indicates economic independence among households from at least the mid-seventh millennium BCE, with domestic storage capacity significantly increasing by the period's end (Çevik, 2019; Çevik & Abay, 2016). Stone tool findings imply that obsidian was sourced from the Island of Melos throughout the Neolithic, with a minor presence of Central Anatolian obsidian in later strata (Guilbeau et al., 2019; Milić, 2014).

Description of burial practices and skeletal material
No burials were found within or between domestic structures during this era. However, several neonatal interments were discovered in open areas with fire installations during the initial phase, including one near a fireplace (ulu008 - B259/JDR) and another adjacent to a grinding stone (ulu009 - B260J60), with a fragment of a newborn (ulu007 - B101 GSH/GTC) unearthed within the fill of Building 43.

**Bahçelievler Höyük**

The Bahçelievler settlement, situated in the urban heart of the town of Bilecik southeast of the Marmara Sea, underwent three seasons of rescue excavations from 2019 to 2021. These excavations were conducted under the auspices of the Bilecik Museum Directorate and were overseen by Erkan Fidan (Fidan, 2020; Fidan et al., 2022; Kolankaya Bostancı & Fidan, 2021). Radiocarbon dating from seven Neolithic occupation layers (Fidan et al., 2022) suggests the inception of Neolithic characteristics in the area around 7100/7000 BCE. Levels 8 to 6 date from 7100/7000 to 6500 BCE, while levels 5 to 2 span from 6500 to 6000 BCE. The earliest settlement layer, beginning around 7100/7000 BCE, already exhibits agriculture, domestication, and pottery (Fidan et al., 2022; Sarıaltun et al., 2024).

The settlement's oldest structures, with round or oval floor plans, were constructed using a branch knitting technique (Fidan et al., 2022). The oldest phase (Level 8) features an oval house with a pit bottom carved into the native soil, whereas subsequent phases (Levels 7-3) have structures with flat bottoms. The interstitial spaces between dwellings served as workshops.

Description of burial practices and skeletal material
The human remains unearthed all pertain to individuals interred in a flexed position within courtyards. A total of 11 in situ human remains were discovered, predominantly from Level 6 [bbh001 (Bahçelievler'20 iskelet No:1), bbh009 (BAH21-B3-70/2-iskelet6), bbh010 (BAH21-C3-35-iskelet9), bbh011 (BAH21-C3-38-iskelet10), bbh012 (BAH21-iskelet4)] (6600-6500 BC). Additionally, two remains from the upper level date to 6500-6300 BCE [bbh007 (BAH21-A4/2-iskelet11, bbh008 (BAH21-B5-24-iskelet7)]. DNA analysis was conducted on seven of these individuals (Table 1). Carbon-14 results are available for two human remains from these strata, but DNA analysis was not feasible due to their poor condition. The dates for these levels are calibrated to 6399-6236 BCE for Level 5 and 6591-6446 BCE for Level 6 (Fidan et al., 2022)

**Bademağacı Höyüğü**

Excavations at Bademağacı Höyük (Duru & Umurtak, 2019), situated 52 km north of Antalya along the Antalya-Burdur highway, were conducted from 1993 to 2010 under the leadership of Prof. Dr. Refik Duru and Prof. Dr. Gülsün Umurtak. The earliest settlement phase, known as the Early Neolithic I (EN I) Period, comprises five levels that reflect a simple village settlement. The initial settlers are believed to have arrived around 7100 BCE, establishing their dwellings. The Neolithic Period persisted until approximately 6100 BCE, with the subsequent four levels indicating a more developed phase in the Neolithic lifestyle (EN II). The first settlement, Early Neolithic I/9 (EN 9), was founded on virgin soil. However, no architectural remains have been discovered from EN I/9. It is likely that the early inhabitants of Bademağacı employed wattle-and-daub construction techniques. Notably, in Level EN I/8, a floor crafted from limestone-based plaster (terrazzo), comparable in hardness to concrete, was unearthed, signifying a remarkable architectural advancement for the era.

From level EN II/4 onward, the construction of houses transitioned to sun-dried bricks. This architectural innovation led to rectangular houses with centrally placed doors on the longer walls. Horseshoe-shaped ovens were typically positioned at the base of the wall opposite the door. These single-room houses included sleeping platforms, clay fireboxes, grinding stones, and designated work areas for grinding. Despite occasional destruction by fire or other causes, the settlements were consistently rebuilt following the same fundamental design. The inhabitants commonly used pottery, bone kitchen tools, stone chisels, axes, and flint and obsidian implements over the 1500-year Neolithic occupation. Additionally, figurines, stamp seals, pintaderas, and beads for necklaces were also prevalent.

Description of burial practices and skeletal material

The burial practices at Bademağacı Höyük during the Early Neolithic Period included approximately 74 interments, comprising around 50 graves and assorted human bone clusters. The human burials found during the excavations between the years 1993-1999 were studied by Anthropologist Dr. Elisabeth Smits from Amsterdam University. The skeletal material found between 2000 and the final excavations year 2010 was studied by Prof. Dr. Yılmaz Selim Erdal.

Three adult burials were found in the lower levels of DA 2, the deep trench that was dug to investigate the early phases of the Neolithic; two of them probably belonged to EN I / 7 and one to settlement levels later than EN I. These three adults were buried under the floor in a flexed position, two placed on their right side and one on their left side.

A large number of burials, the majority of which are infant burials and groups of bones belonging to neonates and young children, were found in various places in Trench A which was excavated to investigate the later phases of the Early Neolithic (EN II) settlement (levels EN II / 4A, 4, 3, 2 and 1). In addition to these, bones belonging to 11 individuals, 10 infants and one adult, were found in an irregular, commingled state within two-roomed house no.8 of level EN II / 3 and another burial in flexed position was found just outside the door of the same house. In EN II, both infants and adults were placed sometimes on their right side and sometimes on their left side in flexed positions, and some were placed on their backs with lower limbs flexed at the knees and contracted on the torso with the upper limbs placed over the breast, also in a flexed position.

Burial offerings were not customary at Bademağacı Höyük, with only a handful of infant burials—and even fewer adult burials—containing beads or simple jewelry. The long duration of the Neolithic occupation at Bademağacı suggests a substantial population. However, the limited number of adult burials uncovered implies that most individuals were interred outside the settlement, possibly in a designated cemetery. It is hypothesized that infants who passed away at or shortly after birth were buried within the settlement, near their parents' homes."

bad017 (BH ENÇ II/2 izole 1): An isolated skeletal element was found in an isolated context. The genetic sex was determined as XX.

bad019 (BH'00 EN II/3 SK 13 A) is an old adult gracile male  (50-60 years old), with a slender structure, and most of the skeletal characteristics of a female. The genetic sex was identified as XY. SK 13 has a perimortem blunt-force traumatic injury on the right parietal bone.

bad022 (BH'00 EN II/3 SK 17): a 1.5-2 years old infant skeleton. The individual has slightly developed rickets with a curved leg and forearm bones. The genetic sex was determined as XY.

bad023 (BH'00 EN II/3 SK 19): a 9-12 months-old infant burial. There are no pathological lesions on the bones. The genetic sex was determined as “consistent with XY but not XX”.

bad024 (BH'02 EN 2 SK 2-4): a newborn. The genetic sex was determined as XX.

bad025 (BH'02 EN 24A House 2): a young adult male who died between 20-24 years old. The genetic sex was determined as XY.

bad026 (BH'03 EN II/3 SK 20): a newborn baby. The genetic sex was determined as XY.

bad030 (BH'04 EN II/3 House 7 SK 34): a middle-age adult male. There is a Colles' fracture of the right radius, possibly related to falling. The genetic sex was determined as XY.

bad033 (BH'02 EN 2 SK 1): a 12-13 year-old adolescent. The individual has a slightly developed porotic hyperostosis. The genetic sex was determined as XY.

bad034 (BH'06 EN II/1 “Orta alan”): a young adult female. The genetic sex was determined as XX.

**Pendik Höyük**

Located on the northeastern coast of the Marmara Sea, Pendik is situated approximately 50m from the shoreline, featuring a bay-like topography due to the protrusion of Cape Temenye into the sea. In close proximity to the site are two freshwater sources and a desiccated stream to the east. The area gained recognition with the advent of railroad construction in the early 20th century. The inaugural archaeological investigations near the railway were conducted in 1960 by Ş. A. Kansu. Subsequent rescue excavations were initiated in 1981 under the auspices of the General Directorate of Antiquities and Museums. Further excavations were undertaken by the Istanbul Archaeological Museum in 1992 and during the period of 2012–2014.

Pendik is emblematic of the nascent stages of the Fikirtepe culture, predominantly inhabited in the second half of the 7th millennium BCE. The archaeological stratum, roughly 3 meters in thickness, spans an extensive area, with its perimeters rendered indistinct by contemporary urban development. The stratigraphy comprises two basal layers attributed to the Archaic Fikirtepe phase and an uppermost layer corresponding to the Classical Fikirtepe phase, surmounted by Byzantine strata. The architectural vestiges predominantly pertain to the Archaic Fikirtepe period. Distinct settlement patterns identified in the Marmara region include agrarian communities and those that intensively exploit environmental resources, echoing a Mesolithic heritage (Karul, 2017; Özdoğan, 2013). Pendik, as a coastal habitation with accessible natural resources, aligns with the latter category.

The predominant architectural form in Pendik is the round or oval wattle-and-daub hut, occasionally featuring a partially subterranean floor. Hearth structures have been discovered within these dwellings and in communal spaces, alongside evidence of external workshops (Harmankaya, 1983; Özdoğan, 1983; Pasinli et al., 1994). Recent excavations have also revealed a rectangular edifice with an interior partitioned into 11 square units, measuring approximately 6 x 3 meters. Postholes were detected solely along the structure's longitudinal sides. The edifice had experienced a conflagration, with charred vessels, flint blades, and stone axes found within. Two adjacent hearths were located in the northeast quadrant of the building, where a unique andiron-like vessel and the skeletal remains of an infant were also uncovered (Kızıltan, 2022). The rectilinear form of this structure is consistent with contemporaneous Anatolian architecture, suggesting a confluence of two cohabiting communities (Özdoğan, 2014). Additionally, a defensive ditch encircling the settlement has been postulated based on the 1981 research (Harmankaya, 1983).

Subsistence in Pendik was multifaceted, encompassing agriculture, pastoralism, and marine resource exploitation. While faunal and malacological studies remain unpublished, preliminary archaeobotanical analyses have been disclosed. The paucity of botanical remains, comprising a mere seven flax seeds and a handful of cereal and legume seeds, may be attributable to sampling and preservation biases. Nonetheless, the limited diversity of species suggests that agriculture was not the principal economic activity, although this interpretation is tentative (Ulaş, 2020).

Description of burial practices and skeletal material
The 1981 and 1992 excavations yielded 32 burials, with recent excavations uncovering an additional 53 burials (Harmankaya, 1983; Pasinli et al., 1994; Yılmaz, 2021). These interments predominantly represent primary burials, though secondary burials have been inferred from skeletal repositioning within some graves (Yılmaz, 2021). The orientation of the interred individuals, often in contracted postures, varied, and a minority of graves contained singular artefacts such as bone spoons (Kızıltan, 2022).

pft003 (Pendik '81 2.5-10) comprised an almost intact primary interment of a middle-aged adult female, exhibiting some masculine cranial traits. The genetic sex was determined as XX.

pft004 (Pendik '81 123-6) is the complete skeleton of a 35-39-year-old female with a complete skeleton, interred in a simple pit, alongside an isolated bone fragment from a secondary burial. Notable dental wear was observed on the individual's anterior teeth. The genetic sex was determined as XX.

**Çatalhöyũk**

The UNESCO World Heritage Site of Çatalhöyük is situated 9 km to the south of Boncuklu Höyük on the Konya Plain in Central Anatolia. The site was discovered in 1959 by James Mellaart (British Institute of Archaeology at Ankara), who conducted the first excavations between 1961 and 1965. New excavations took place between 1993 and 2017 under the direction of Ian Hodder (Stanford University).

The site consists of two separate mounds or "tells". The larger East Mound, covering an area of 13 ha, has been dated to c. 7100-5950 cal BCE (Bayliss et al., 2015) and corresponds to the Ceramic Neolithic period. The smaller West Mound is dated to the Early Chalcolithic period and was occupied until the middle of the 6th millennium BCE (Orton et al., 2018). For much of its occupation, settlement on the Neolithic East Mound consisted of tightly clustered mudbrick domestic structures and open spaces used for waste disposal, animal penning, and other activities. To date, large, clearly identifiable communal structures have not been documented at Çatalhöyük. Instead, individual houses served as foci for domestic activities such as craft production, food storage and processing. Houses also served as sites for ritual behaviours such as subfloor burials, wall paintings and other symbolic elaborations such as the embedding of animal bones within walls, platforms and benches (Hodder & Cessford, 2004). There is ample evidence for cultivating domesticated cereal crops and keeping domesticated sheep and goats from the earliest occupation phases of the site (Bogaard et al., 2009; Russell et al., 2013). Wild animal species, including aurochs, also formed part of the diet, but in later occupation phases (6500-5950 calBC) there is evidence for the gradual introduction of domesticated cattle (Russell et al., 2013; Wolfhagen & Price, 2017).

Description of burial practices and skeletal material

During the 1993-2017 Hodder excavations, the skeletal remains of over 700 individuals were excavated from stratified Neolithic contexts at Çatalhöyük (Larsen et al., 2019). Primary interments (n = 471 individuals), typically placed under house floors, are the dominant method of burial at the site (Boz & Hager, 2013; S. D. Haddow et al., 2021). Individuals were typically buried in simple pits under the raised platforms of the central room, although very young individuals (i.e. prenates, neonates and infants) were also buried in storage rooms and near ovens and hearths (Boz et al., 2006; Boz & Hager, 2013; S. D. Haddow et al., 2021). Secondary burials of loose or partially articulated skeletal remains (n = 96) are also documented (S. D. Haddow et al., 2021; S. Haddow & Knüsel, 2017). Towards the end of the occupation on the Neolithic East Mound, intramural burials became increasingly rare (S. D. Haddow et al., 2021; Marciniak et al., 2015), and on the Chalcolithic West Mound they are almost entirely absent (Anvari et al., 2017; Biehl, 2012).

Of the 471 Çatalhöyük individuals from primary burial contexts (Knüsel et al., 2021), there are 178 adults (20+ years), 29 adolescents (12-20 years), 90 children (3-12 years), 67 infants (2 months-3 years), 85 neonates (0-2 months), and 22 prenates (> 38 weeks in utero). Among the adults and adolescents whose sex could be determined (n = 155), 89 individuals (57%) were assessed as females or possible females, while 66 individuals (43%) were assessed as males or possible males.

(cch153) sk.10527: Building 43, Level South L Primary burial of pre-term foetus (approx. 32 weeks in utero – based on long bone length) placed under the west platform of B.43, excavated in 2004 (DNA identifies this individual as XY). 7100-6700 BCE.

(cch245) sk.12876: Building 56, Level South R Secondary burial of a 5-year-old (+/- 1.5yrs) child (based on dental development) placed under the east platform of B.56, excavated in 2006 (DNA identifies this individual as XX). 6500-6300 BCE.

(cch251) sk.2197: Building 1, Level North G Primary burial of a pre-term foetus (5 months in utero - based on long bone length) placed in the southwest corner of B.1, excavated in 1997 (DNA identifies this individual as XX). 6700-6500 BCE.

(cch289) sk.1885: Building 50, Level South M Primary burial of a 7-year-old (+/- 2yrs) child (based on dental development) found in the southwest corner of B.50, excavated in 1995 (DNA identifies this individual as XY). Radiocarbon dating places this individual between 6905–6885 cal BCE (1%) or 6825–6635 cal BCE (92%) or 6625–6600 cal BCE (2%).

(cch294) sk.2779: Building 50, Level South M Primary burial of a neonate (0-2 months at death based on measurements of the basi-occipital bone), excavated in 1997 (DNA identifies this individual as XY).

(cch311) sk8587: Building 114, Level North G Primary burial of a neonate (0-2 months at death – based on long bone length) excavated in 2002 and located under the southeast platform of B.114 (DNA identifies this individual as XX).

(cch348) sk2141: Building 1, Level North G Primary burial of a 9-month-old (+/- 3 month) infant (based on dental development) under northwest platform of B.1, excavated in 1997 (DNA identifies this individual as XY). 6700-6500 BCE.

### Supplemental Figures

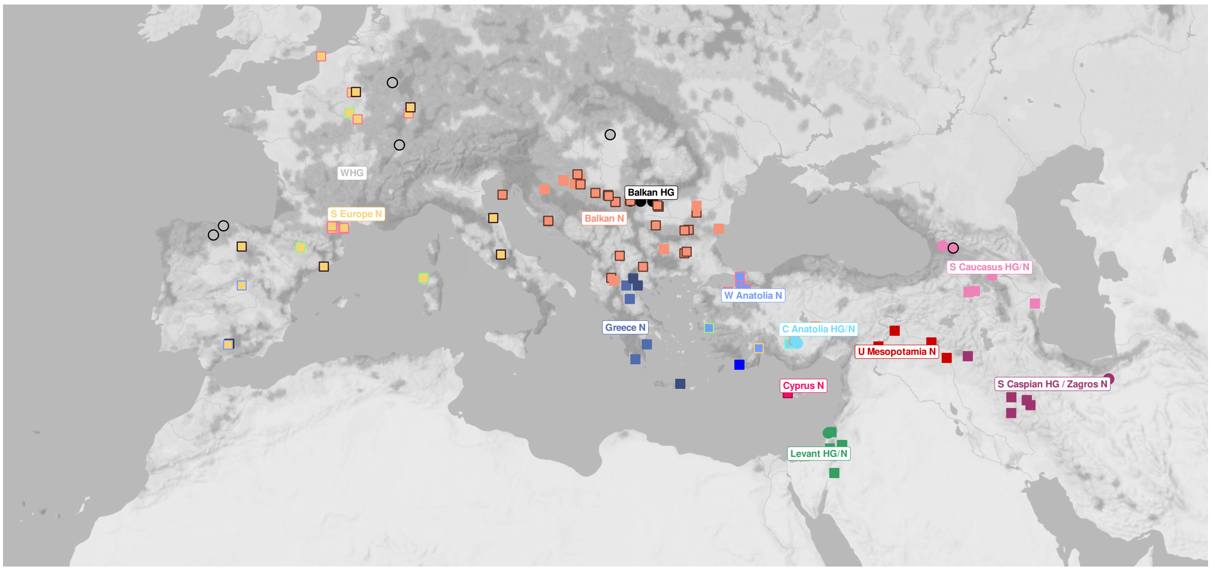


**Figure S1:** Map of W Eurasian sites with palaeogenomic data used in this study (see also **Table S2**).

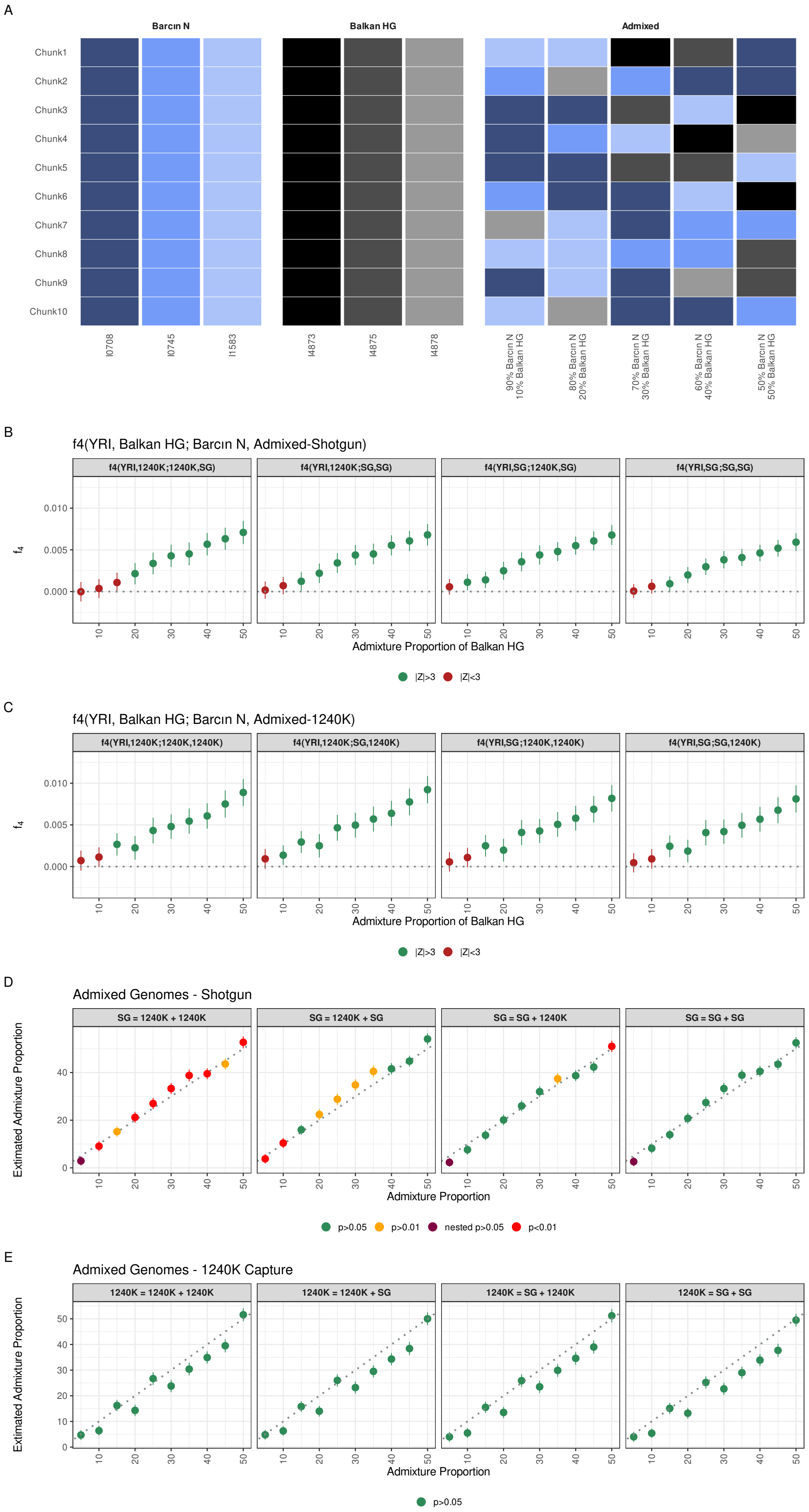
A
Barcın N
Balkan HG
Admixed
Chunk1
Chunk2
Chunk3
Chunk4
Chunk5
Chunk6
Chunk7
Chunk8
Chunk9
Chunk10
I0708
I0745
I1583
I4873
I4875
I4878
90% Barcın N 10% Balkan HG
80% Barcın N 20% Balkan HG
70% Barcın N 30% Balkan HG
60% Barcın N 40% Balkan HG
50% Barcın N 50% Balkan HG
B
f4(YRI, Balkan HG; Barcın N, Admixed-Shotgun)
f4(YRI,1240K;1240K,SG)
f4(YRI,1240K;SG,SG)
f4(YRI,SG;1240K,SG)
f4(YRI,SG;SG,SG)
Admixture Proportion of Balkan HG
|Z|>3
|Z|<3
C
f4(YRI, Balkan HG; Barcın N, Admixed-1240K)
f4(YRI,1240K;1240K,1240K)
f4(YRI,1240K;SG,1240K)
f4(YRI,SG;1240K,1240K)
f4(YRI,SG;SG,1240K)
Admixture Proportion of Balkan HG
|Z|>3
|Z|<3
D
Admixed Genomes - Shotgun
SG = 1240K + 1240K
SG = 1240K + SG
SG = SG + 1240K
SG = SG + SG
Extimated Admixture Proportion
Admixture Proportion
p>0.05
p>0.01
nested p>0.05
p<0.01
E
Admixed Genomes - 1240K Capture
1240K = 1240K + 1240K
1240K = 1240K + SG
1240K = SG + 1240K
1240K = SG + SG
Extimated Admixture Proportion
Admixture Proportion
p>0.05

**Figure S2:** The performance of qpAdm models and $f_4$-tests on genomic admixture simulations using the genoMIX algorithm and using Barcın (NW Anatolia) and Balkan HG genomes as sources. A: Demonstration of two sources, Barcın and Balkan HG genomes, and the resulting admixed genomes (on the right) produced by collating chunks from both sources. B and C: The performance of $f_4$-tests when the admixed genome is produced using shotgun data ("SG") or using 1240k capture data ("1240K"), respectively. The x-axis shows admixture proportions. Green points indicate significant $f_4$ test results. D and E: The performance of qpAdm models when the admixed genome is produced using shotgun data ("SG") and 1240k capture data ("1240K"), respectively. We varied the data type for source populations as well, which is indicated above each panel as follows: Data Type of Admixed Genome = Data Type of Source Population 1 + Data Type of Source Population 2. The x-axis shows admixture proportions. Green points indicate feasible models.

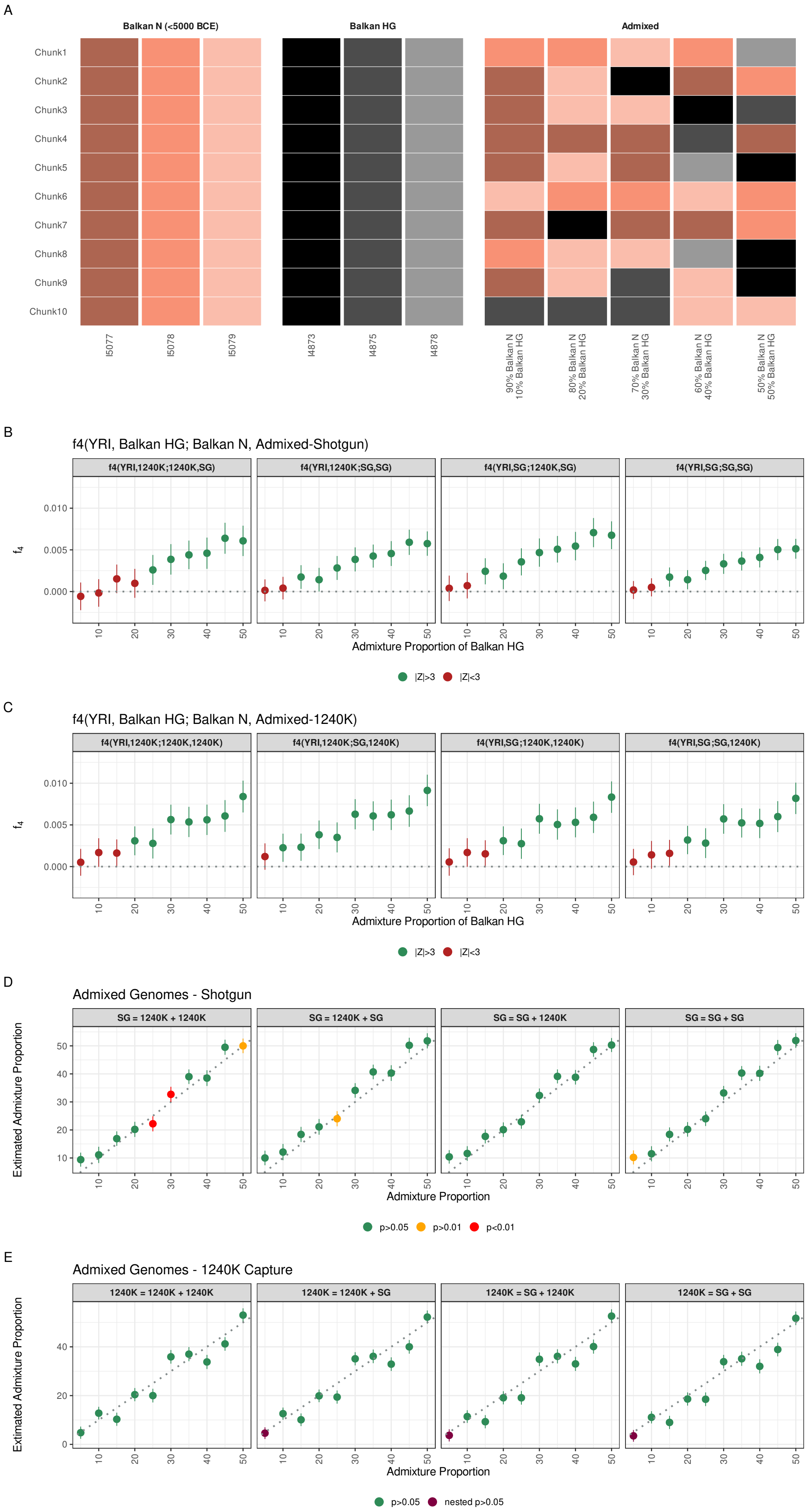

A
Balkan N (<5000 BCE)
Balkan HG
Admixed
Chunk1
Chunk2
Chunk3
Chunk4
Chunk5
Chunk6
Chunk7
Chunk8
Chunk9
Chunk10
I5077
I5078
I5079
I4873
I4875
I4878
90% Balkan N 10% Balkan HG
80% Balkan N 20% Balkan HG
70% Balkan N 30% Balkan HG
60% Balkan N 40% Balkan HG
50% Balkan N 50% Balkan HG
B
f4(YRI, Balkan HG; Balkan N, Admixed-Shotgun)
f4(YRI,1240K;1240K,SG)
f4(YRI,1240K;SG,SG)
f4(YRI,SG;1240K,SG)
f4(YRI,SG;SG,SG)
$f_4$
Admixture Proportion of Balkan HG
|Z|>3
|Z|<3
C
f4(YRI, Balkan HG; Balkan N, Admixed-1240K)
f4(YRI,1240K;1240K,1240K)
f4(YRI,1240K;SG,1240K)
f4(YRI,SG;1240K,1240K)
f4(YRI,SG;SG,1240K)
$f_4$
Admixture Proportion of Balkan HG
|Z|>3
|Z|<3
D
Admixed Genomes - Shotgun
SG = 1240K + 1240K
SG = 1240K + SG
SG = SG + 1240K
SG = SG + SG
Extimated Admixture Proportion
Admixture Proportion
p>0.05
p>0.01
p<0.01
E
Admixed Genomes - 1240K Capture
1240K = 1240K + 1240K
1240K = 1240K + SG
1240K = SG + 1240K
1240K = SG + SG
Extimated Admixture Proportion
Admixture Proportion
p>0.05
nested p>0.05

**Figure S3:** The performance of qpAdm models and $f_4$-tests on genomic mixture simulations using the genoMIX algorithm and using Balkan Neolithic and Balkan HG genomes as sources. The figure reports the same type of analysis as Figure S2 except for using Balkan Neolithic instead of Barcın as the source.

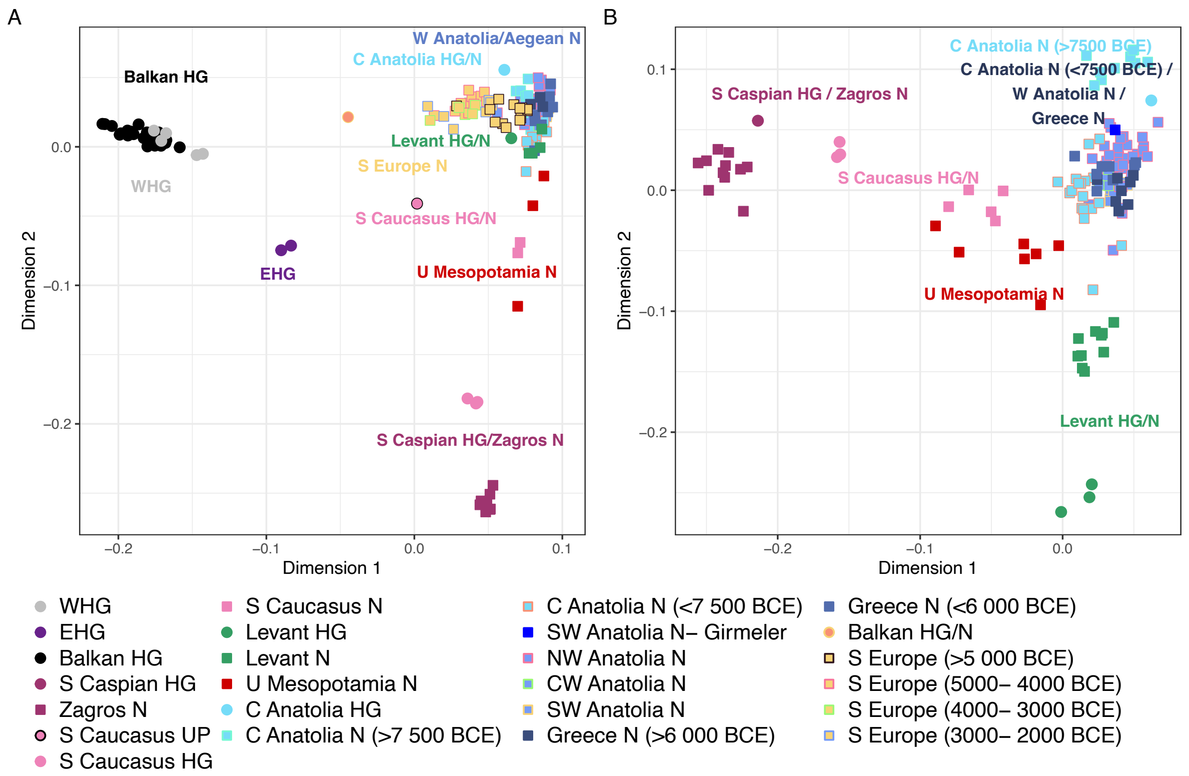


**Figure S4:** Multidimensional scaling plot of UP/EH palaeogenomes from (A) W Eurasia and (B) SW Asia. The plot summarizes $f_3$-based distances (Methods).

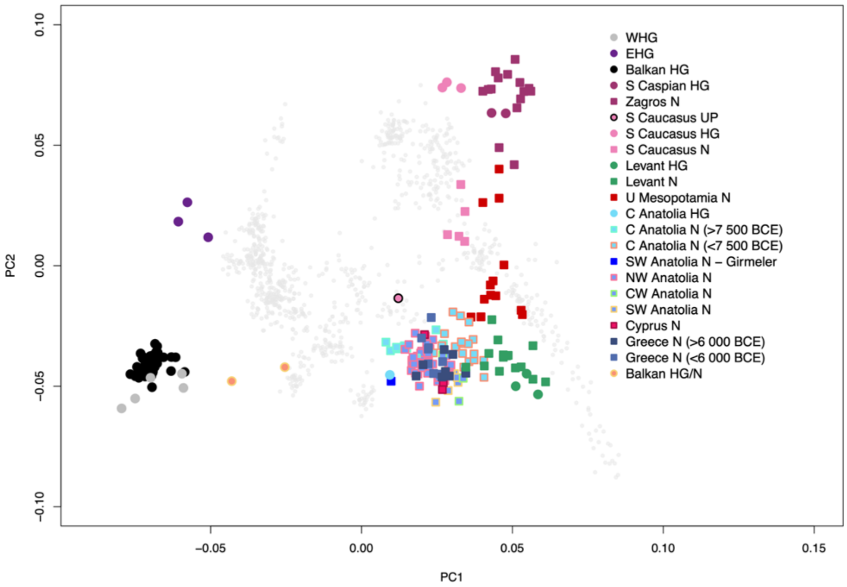


**Figure S5:** Principal components analysis (PCA) of W Eurasian modern-day and ancient genomes. UP/EH palaeogenomes from SW Asia and S Europe have been projected on a PCA space created using modern-day genomes from the Human Origins dataset.

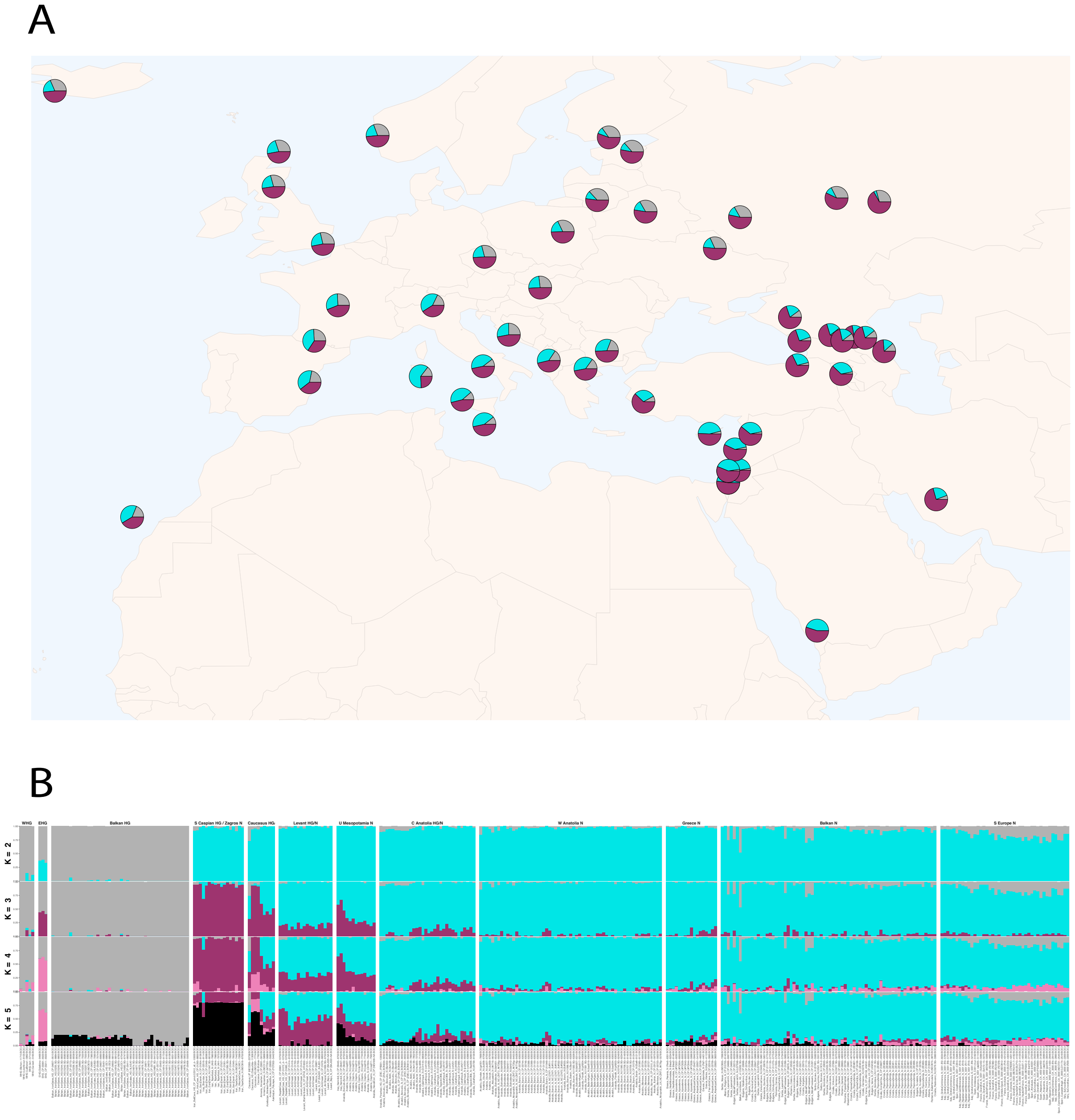


**Figure S6:** ADMIXTURE analysis of West Eurasian present-day (A) and ancient (B) genomes. K=3 yields the lowest cross-validation (CV) error. Panel A shows K=3 ADMIXTURE results for West Eurasian present-day genomes, where Panel B shows results from K=2 to K=5.

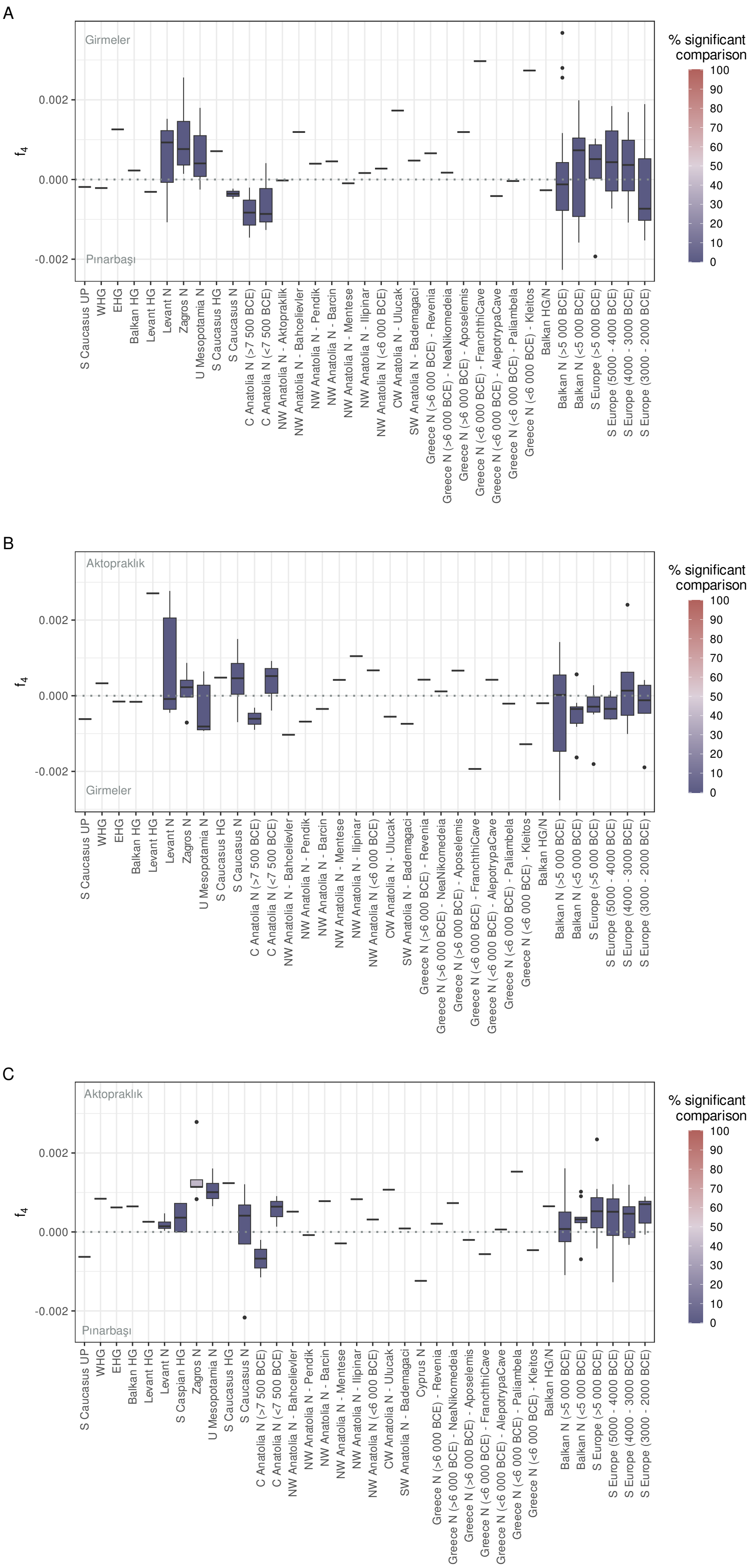
A
Girmeler
Pınarbaşı
f4
0.002
0.000
-0.002
% significant comparison
100
90
80
70
60
50
40
30
20
10
0
S Caucasus UP
WHG
EHG
Balkan HG
Levant HG
Levant N
Zagros N
U Mesopotamia N
S Caucasus HG
S Caucasus N
C Anatolia N (>7 500 BCE)
C Anatolia N (<7 500 BCE)
NW Anatolia N - Aktopraklik
NW Anatolia N - Bahcelievler
NW Anatolia N - Pendik
NW Anatolia N - Barcin
NW Anatolia N - Mentese
NW Anatolia N - Ilipinar
NW Anatolia N (<6 000 BCE)
CW Anatolia N - Ulucak
SW Anatolia N - Bademagaci
Greece N (>6 000 BCE) - Revenia
Greece N (>6 000 BCE) - NeaNikomedeia
Greece N (>6 000 BCE) - Aposelemis
Greece N (<6 000 BCE) - FranchthiCave
Greece N (<6 000 BCE) - AlepotrypaCave
Greece N (<6 000 BCE) - Paliambela
Greece N (<6 000 BCE) - Kleitos
Balkan HG/N
Balkan N (>5 000 BCE)
Balkan N (<5 000 BCE)
S Europe (>5 000 BCE)
S Europe (5000 - 4000 BCE)
S Europe (4000 - 3000 BCE)
S Europe (3000 - 2000 BCE)
B
Aktopraklık
Girmeler
f4
0.002
0.000
-0.002
% significant comparison
100
90
80
70
60
50
40
30
20
10
0
S Caucasus UP
WHG
EHG
Balkan HG
Levant HG
Levant N
Zagros N
U Mesopotamia N
S Caucasus HG
S Caucasus N
C Anatolia N (>7 500 BCE)
C Anatolia N (<7 500 BCE)
NW Anatolia N - Bahcelievler
NW Anatolia N - Pendik
NW Anatolia N - Barcin
NW Anatolia N - Mentese
NW Anatolia N - Ilipinar
NW Anatolia N (<6 000 BCE)
CW Anatolia N - Ulucak
SW Anatolia N - Bademagaci
Greece N (>6 000 BCE) - Revenia
Greece N (>6 000 BCE) - NeaNikomedeia
Greece N (>6 000 BCE) - Aposelemis
Greece N (<6 000 BCE) - FranchthiCave
Greece N (<6 000 BCE) - AlepotrypaCave
Greece N (<6 000 BCE) - Paliambela
Greece N (<6 000 BCE) - Kleitos
Balkan HG/N
Balkan N (>5 000 BCE)
Balkan N (<5 000 BCE)
S Europe (>5 000 BCE)
S Europe (5000 - 4000 BCE)
S Europe (4000 - 3000 BCE)
S Europe (3000 - 2000 BCE)
C
Aktopraklık
Pınarbaşı
f4
0.002
0.000
-0.002
% significant comparison
100
90
80
70
60
50
40
30
20
10
0
S Caucasus UP
WHG
EHG
Balkan HG
Levant HG
Levant N
S Caspian HG
Zagros N
U Mesopotamia N
S Caucasus HG
S Caucasus N
C Anatolia N (>7 500 BCE)
C Anatolia N (<7 500 BCE)
NW Anatolia N - Bahcelievler
NW Anatolia N - Pendik
NW Anatolia N - Barcin
NW Anatolia N - Mentese
NW Anatolia N - Ilipinar
NW Anatolia N (<6 000 BCE)
CW Anatolia N - Ulucak
SW Anatolia N - Bademagaci
Cyprus N
Greece N (>6 000 BCE) - Revenia
Greece N (>6 000 BCE) - NeaNikomedeia
Greece N (>6 000 BCE) - Aposelemis
Greece N (<6 000 BCE) - FranchthiCave
Greece N (<6 000 BCE) - AlepotrypaCave
Greece N (<6 000 BCE) - Paliambela
Greece N (<6 000 BCE) - Kleitos
Balkan HG/N
Balkan N (>5 000 BCE)
Balkan N (<5 000 BCE)
S Europe (>5 000 BCE)
S Europe (5000 - 4000 BCE)
S Europe (4000 - 3000 BCE)
S Europe (3000 - 2000 BCE)

**Figure S7**: $f_4$ tests with Pınarbaşı, Girmeler and Aktopraklık genomes as reference populations. The graphs show results of $f_4$ tests of the form $f_4$*(Yoruba, Test; A, B)*, where *Test* populations are shown in the x-axis, while *A* and *B* either of Pınarbaşı, Girmeler and Aktopraklık genomes indicated on the upper and lower left side axes of each panel. The colours indicate the proportion of nominally significant comparisons at $|Z|>3$. Among the comparisons involving single genomes shown by single lines, none were significant. The results indicate that third populations cannot distinguish between Pınarbaşı, Girmeler and Aktopraklık.

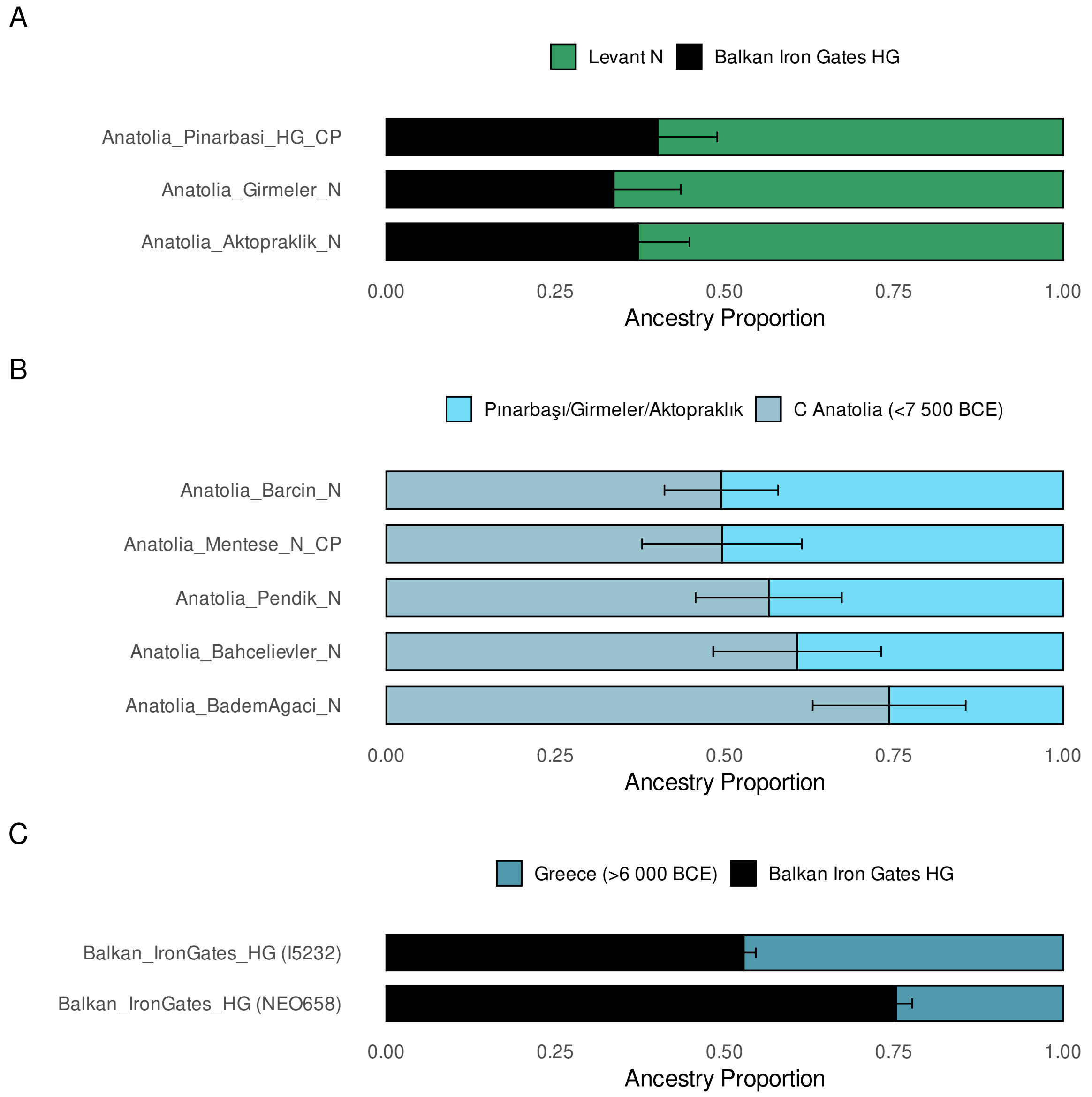


**Figure S8:** qpAdm models for Epipalaeolithic and Neolithic populations. A) Local populations of C and W Anatolia can be explained as admixture of Balkan HG and Levant N, B) The majority of W Anatolian Neolithic palaeogenomes can be explained as admixed between immigrants from C Anatolia Neolithic (Çatalhöyük) and local groups (Pınarbaşı/Girmeler/Aktopraklık), C) Two individuals from Balkan HG groups can be explained as an admixture of Balkan HG and Greece N. The bars represent the coefficients of source populations (shown in the x-axis). The source populations used in qpAdm models are color-coded and shown above each figure, and error bars show one standard error.

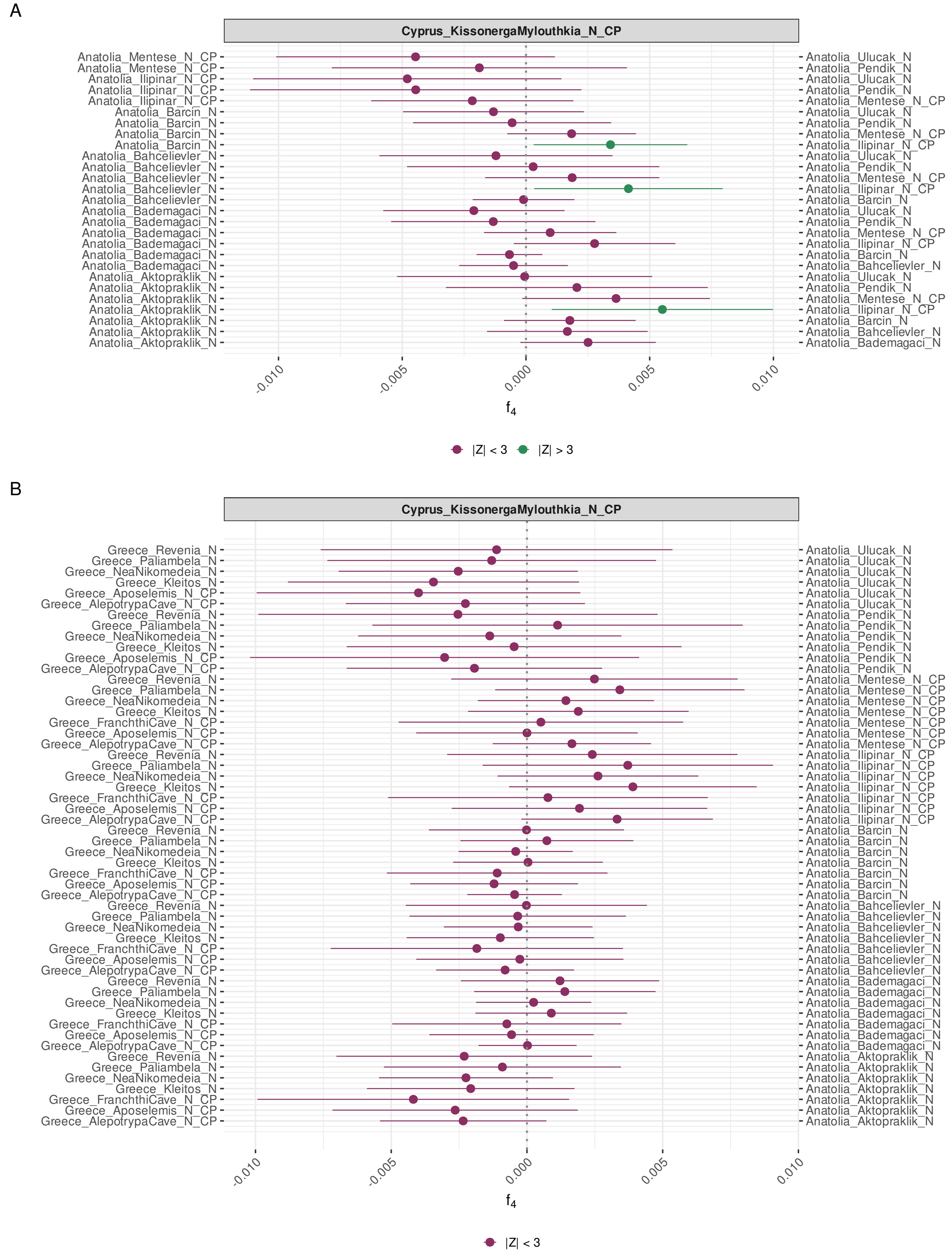


**Figure S9**: The affinities of Cyprus Kissonerga Myiouthkia Neolithic to different sited from (A) W Anatolia and (B) Greece analysed using $f_4$-tests. Colour coding indicates the proportion of nominally significant tests out of all comparisons (at $|Z|>3$). The fact that Cyprus shows affinity to Ilıpınar over other NW Anatolian sites may be driven by technical biases, namely both Cyprus and Ilıpınar were 1240k capture data.

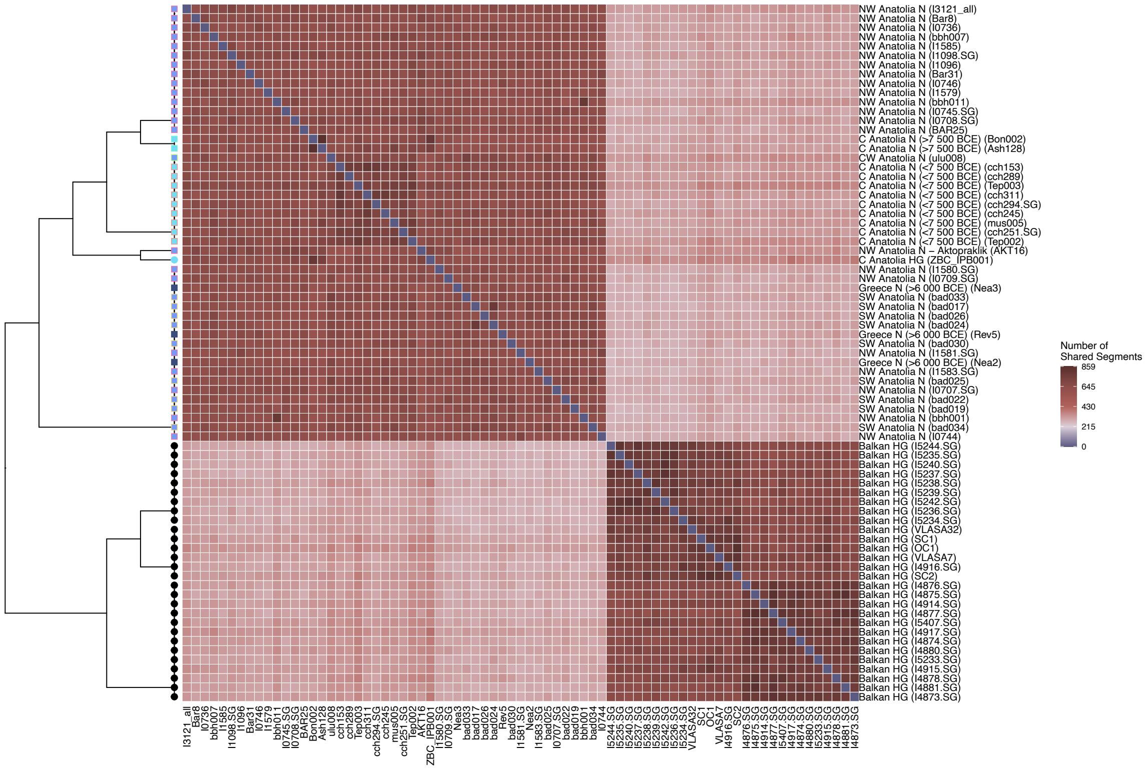


**Figure S10:** Pairwise haplotype counts shared between individuals from Anatolia Neolithic, Greece Neolithic and Balkan HG populations. The clustering on the left-side was performed with FineSTRUCTURE on shared autosomal haplotype counts. The clustering between C Anatolia HG (Pınarbaşı) and Aktopraklik (AKT16) is notable. The Girmeler genome could not be included in this analysis due to its low coverage, which did not allow imputation.

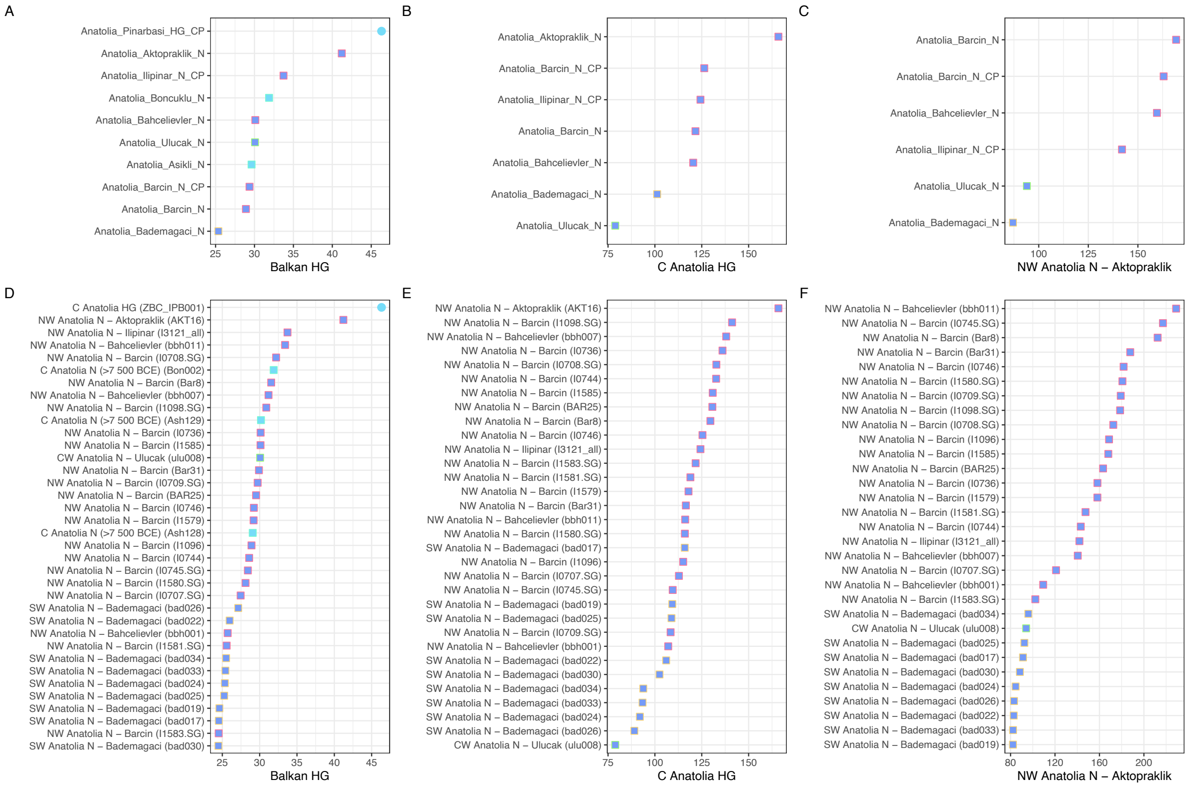


**Figure S11:** Haplotype-sharing statistics between Aegean/Anatolian populations and Balkan HG (Panle A and D), C Anatolia HG (Pınarbaşı) (Panel B and E), Aktopraklık (Panel C and F). Upper panels (A,B,C) are population means whereas lower panels (D,E,F) indicate individual-level computations. Shared chunk lengths were estimated in cM by ChromoPainter for each pair of individuals. The Girmeler genome could not be included in this analysis due to its low coverage, which did not allow imputation. The "CP" suffix stands for aDNA data produced using capture technologies (instead of shotgun).

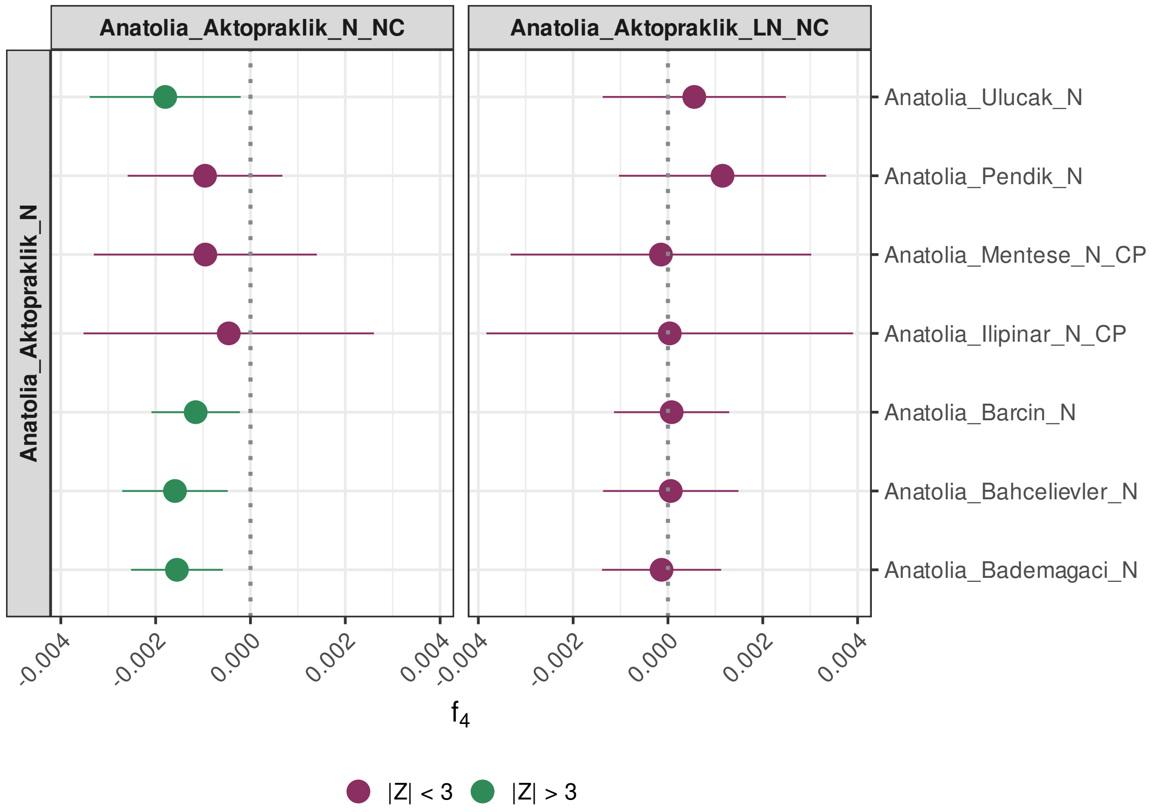


**Figure S12:** The affinities of Aktopraklık neutrolome capture genomes to AKT16 versus W Anatolia analysed using $f_4$-tests. Left panel: Early Neolithic (>6000 BCE); right panel: Late Neolithic (<6000 BCE) genomes from Aktopraklık. The left population is the Aktopraklık shotgun genome (AKT16) and the right populations are W Anatolian groups. Colour coding indicates the proportion of nominally significant tests out of all comparisons (at |Z|>3).

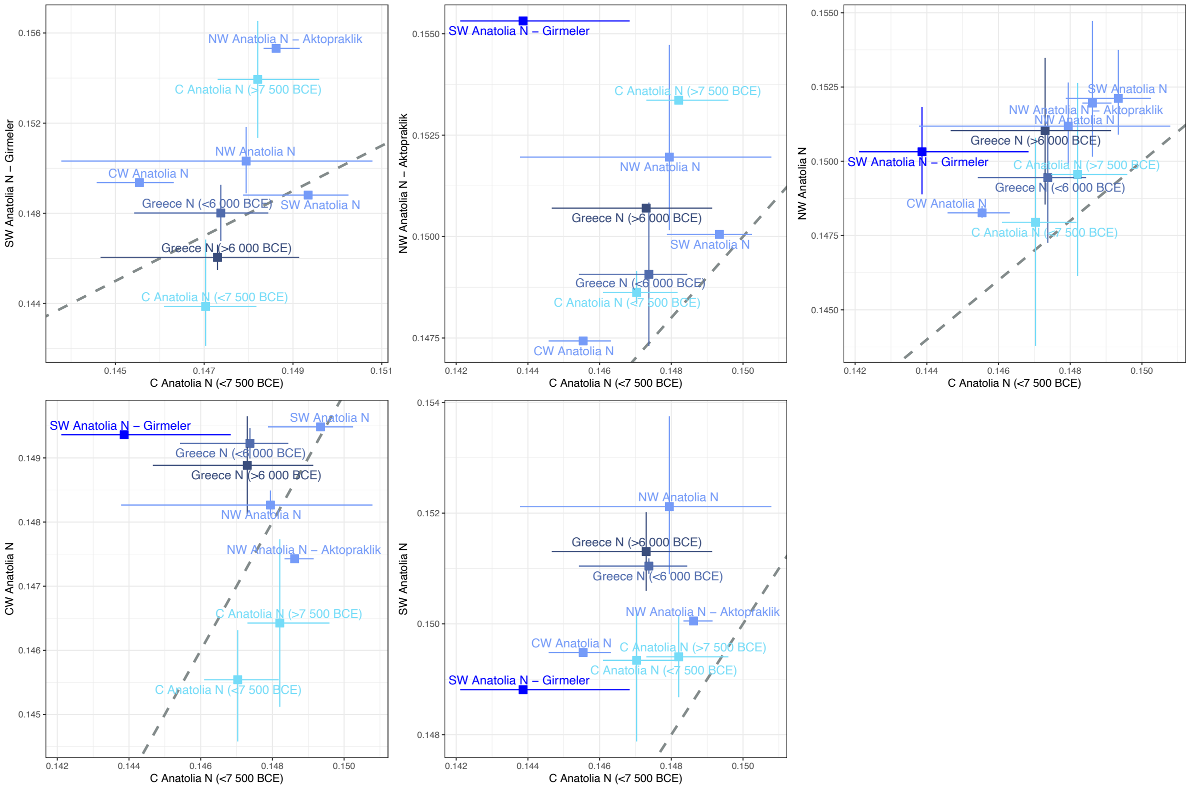


**Figure S13:** Outgroup $f_3$ statistics values of form *$f_3$(Yoruba; C Anatolia N (<7 500 BCE), Anatolia/Greece)* and *$f_3$(Yoruba; W Anatolia N, Anatolia/Greece)* plotted against each other. Points show the mean $f_3$value of each region, whereas lines show each region's upper and lower bounds.

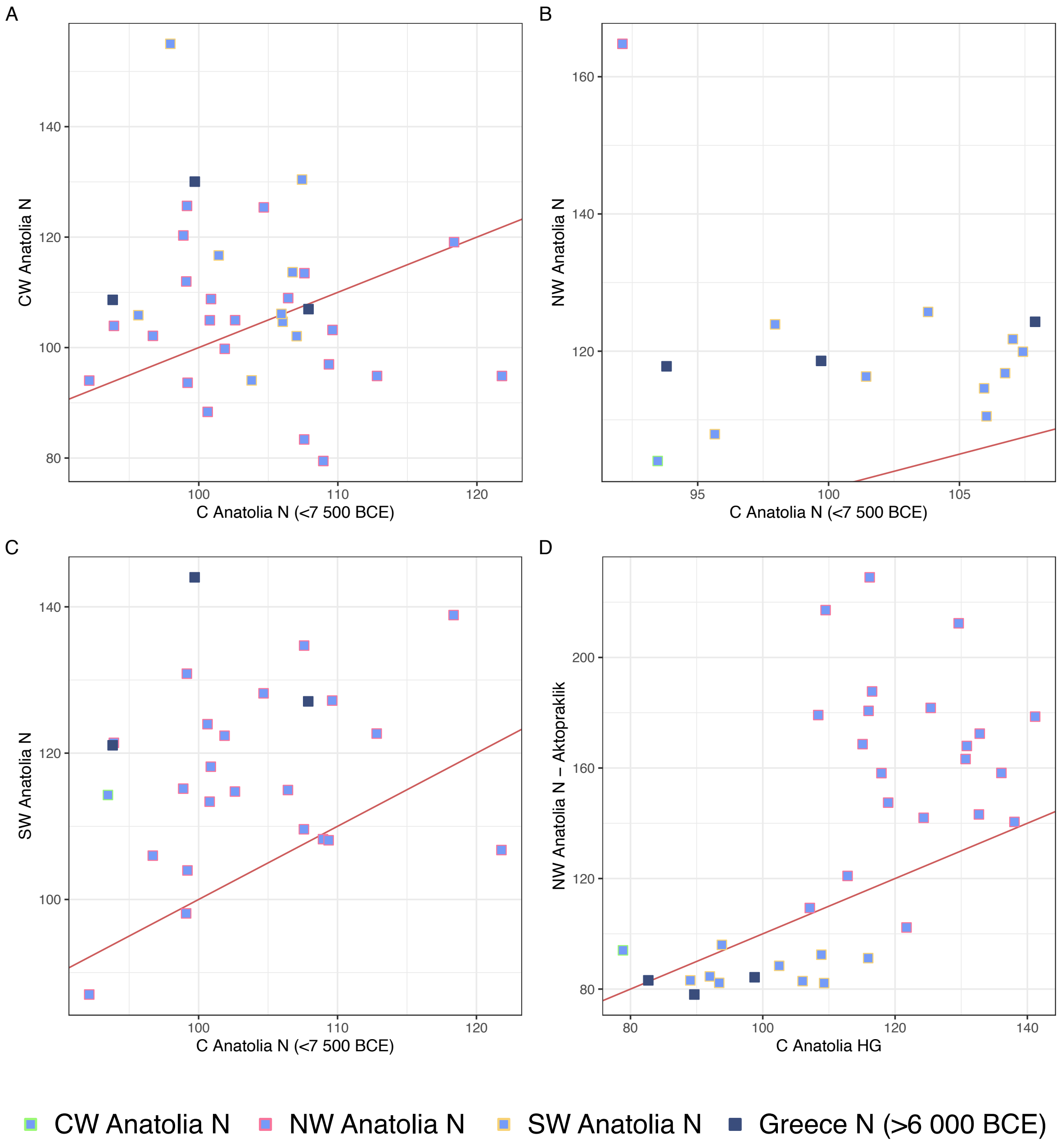


**Figure S14:** Two-dimensional haplotype sharing statistics of relevant individuals. Panels A, B and C shows C Anatolia N (<7500 BCE) on x-axes and CW Anatolia N, NW Anatolia N, SW Anatolia N on y-axis, respectively, while Panel D infers C Anatolia HG and Aktopraklık on x and y-axes. Each point indicates one ancient genome from NW Anatolia and Greece.

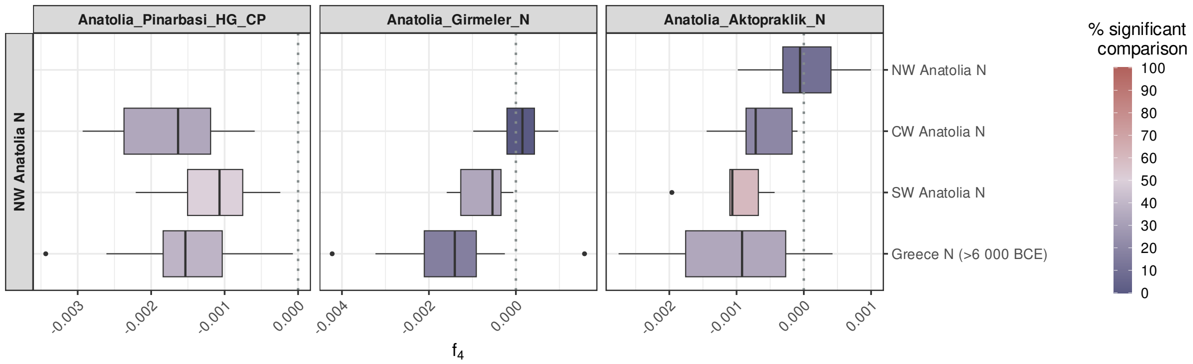


**Figure S15:** The local genetic profile of W Anatolia shows a higher affinity to NW Anatolia than other regions of the Aegean. The panels show results of $f_4$ tests in the form of $f_4$*(Yoruba, Pınarbaşı/Girmeler/Aktopraklık; NW Anatolia N, X)*, where *X* represents any site within an Aegean regional/temporal group indicated on the right side, and *Pınarbaşı/ Girmeler/Aktopraklık* represent the genetic profile of local W Anatolians. The tests were performed at the population (site) level. The colours indicate the proportion of nominally significant comparisons at |Z|>3.

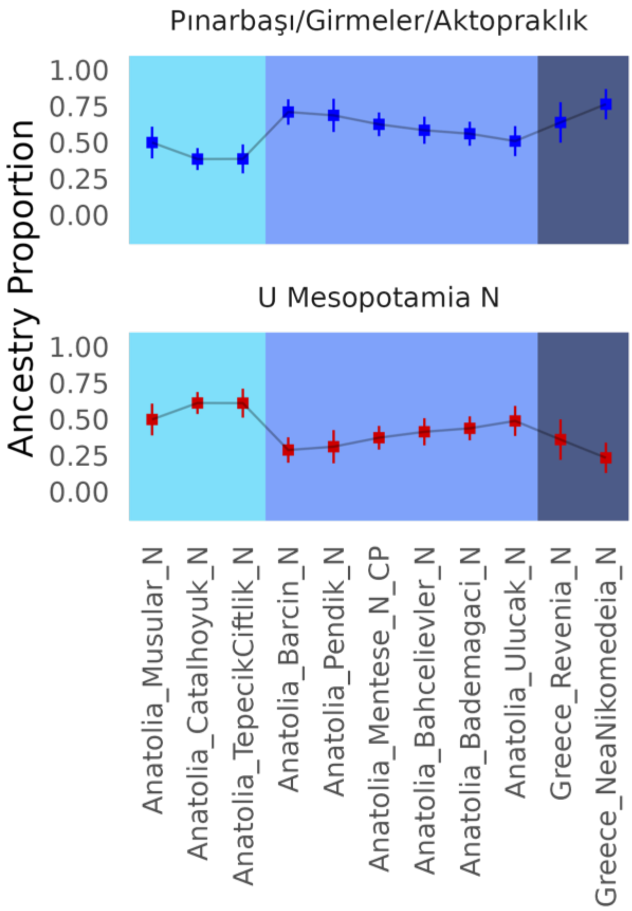


**Figure S16:** Pınarbaşı/Girmeler/Aktopraklık- and U Mesopotamia-related ancestry proportions in C Anatolian and Aegean genomes from archaeological sites dating <7500 BCE. The proportions are based on feasible qpAdm models performed on site-based genomic samples, shown on the x-axis (Methods). From left to right different colours show sites from C Anatolia (<7500 BCE), W Anatolia and Greece. The top and lower panels show Pınarbaşı/Girmeler/Aktopraklık and U Mesopotamia proportions, respectively. We must note that modeling Çatalhöyük with Girmeler does not necessarily imply that W Anatolian indigenous populations contributed directly to the Çatalhöyük gene pool (indeed, models with Aşıklı/Boncuklu (C Anatolia >7500 BCE) and Çayönü (U Mesopotamia) as source can also explain Çatalhöyük (C Anatolia <7500 BCE)]. Likewise, modelling Barcın (NW Anatolia) with Çayönü does not imply that U Mesopotamian populations contributed directly to the Barcın gene pool (indeed, models with Girmeler and Çatalhöyük as source can also explain Barcın).

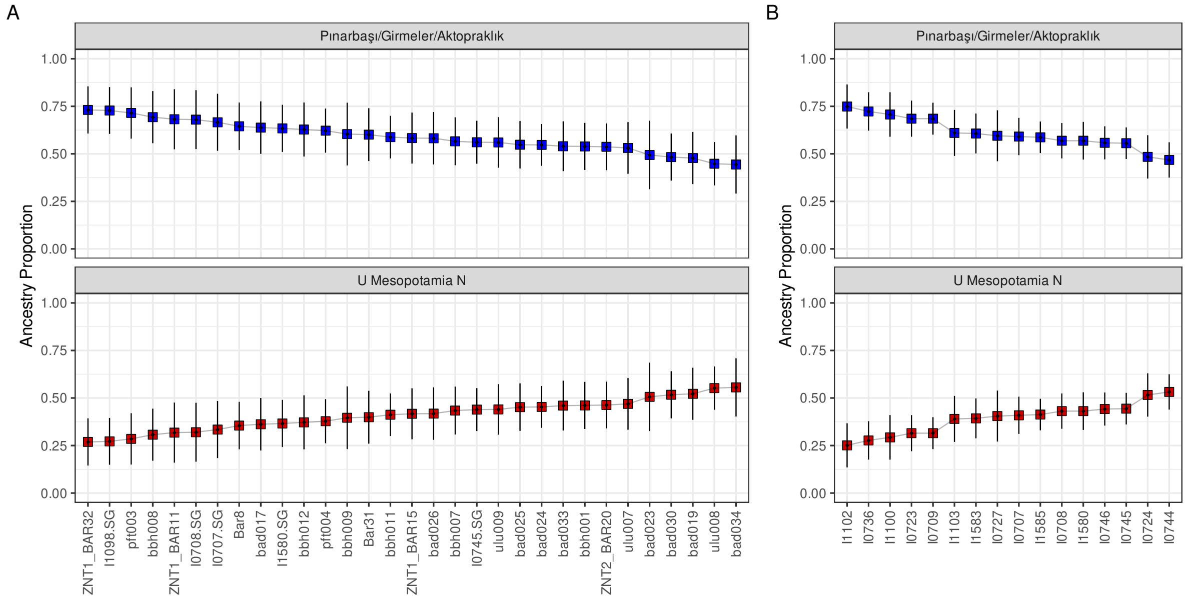


**Figure S17:** Pınarbaşı/Girmeler/Aktopraklık- and Çayönü-related ancestry (U Mesopotamia) proportions in W Anatolian individual A) shotgun-sequenced or genome-wide captured B) 1240K captured genomes from <7500 BCE. Pınarbaşı/Girmeler/Aktopraklık- and Çayönü-related ancestry proportion estimates in W Anatolian genomes from archaeological sites dating >6000 BCE. The graph is based on qpAdm modelling and contains the same information as Figure S16 but has been performed on individual genomes instead of all usable samples per site.

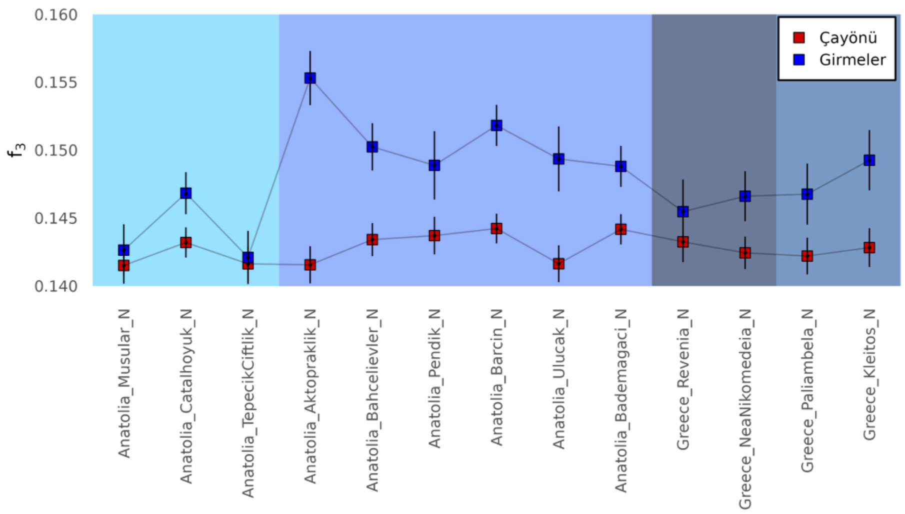


**Figure S18:** Shared genetic drift between Girmeler (SW Anatolia) or Çayönü (U Mesopotamia) and W Anatolian/Greece populations calculated by outgroup-$f_3$ statistics in forms of *$f_3$(Yoruba, Girmeler|Çayönü, W Anatolia/Greece)*. From left to right colours indicate sites from C Anatolia (<7500 BCE), W Anatolia, Greece (>6000 BCE) and Greece (<6000 BCE).

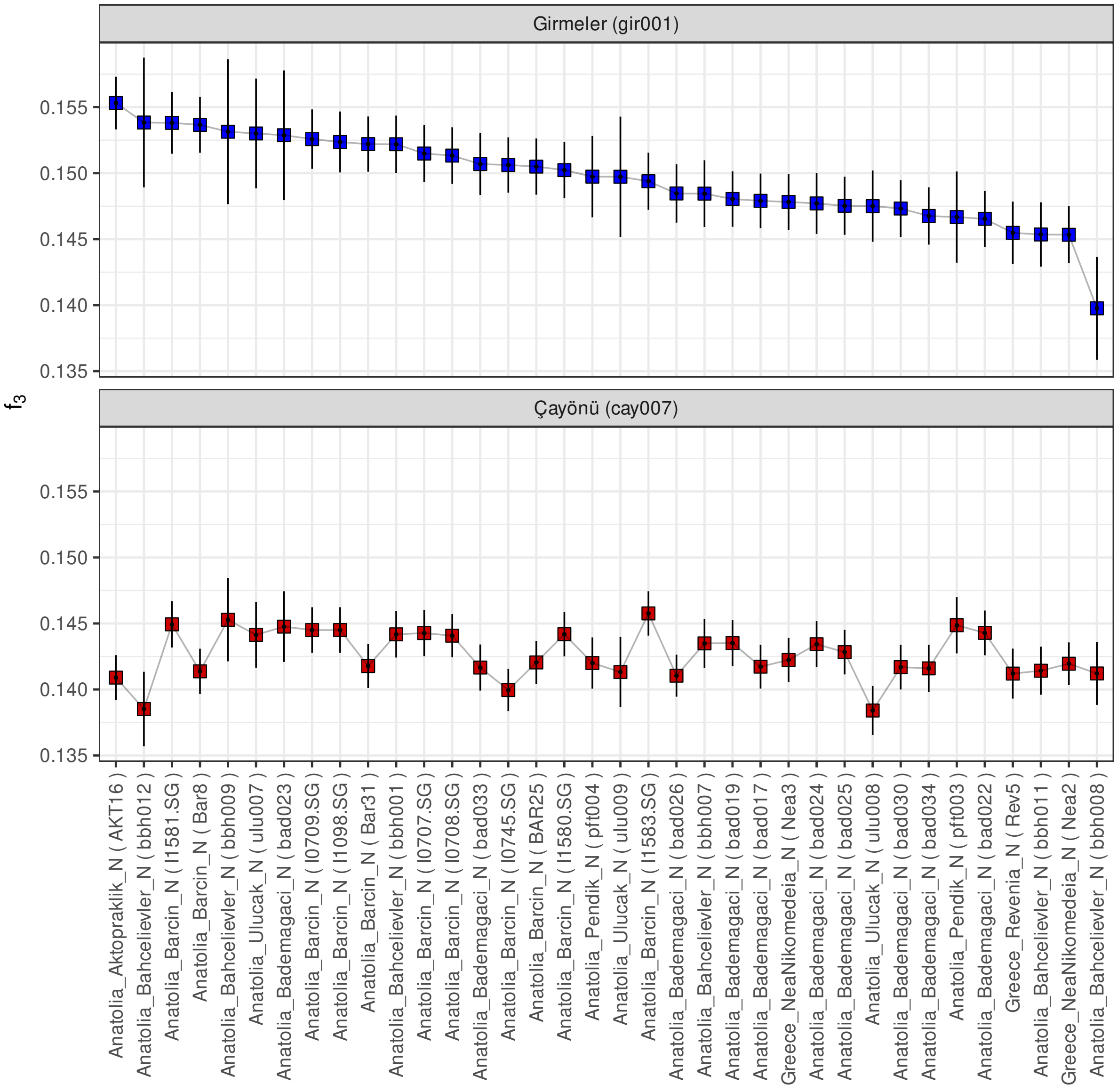


**Figure S19:** Shared genetic drift between Girmeler (SW Anatolia) or Çayönü (U Mesopotamia) and W Anatolian/Greece individuals calculated by outgroup-$f_3$ statistics in forms of $f_3$*(Yoruba, Girmeler|Çayönü, W Anatolia/Greece)*.

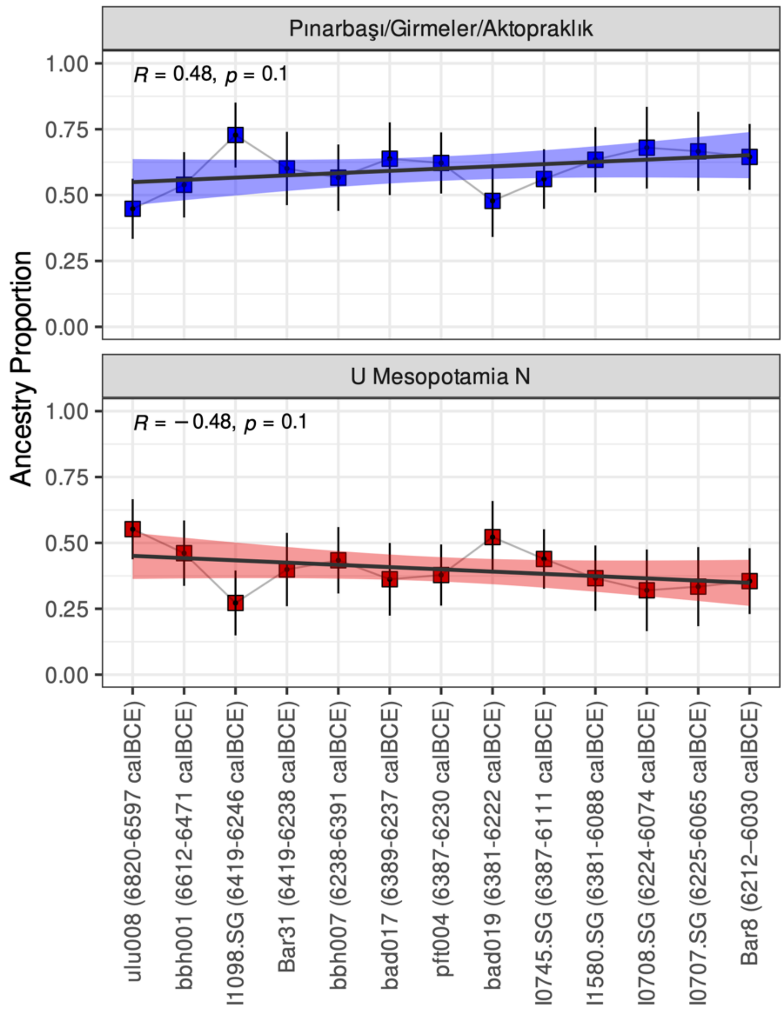


**Figure S20:** Pınarbaşı/Girmeler/Aktopraklık- and U Mesopotamia (Çayönü)-related ancestry proportions in W Anatolian individual genomes from pre-6000 BCE. The graph is based on qpAdm modeling and contains the same information as Figure S17 except it only includes genomes that have C14 dates and are ordered from earliest to the latest.

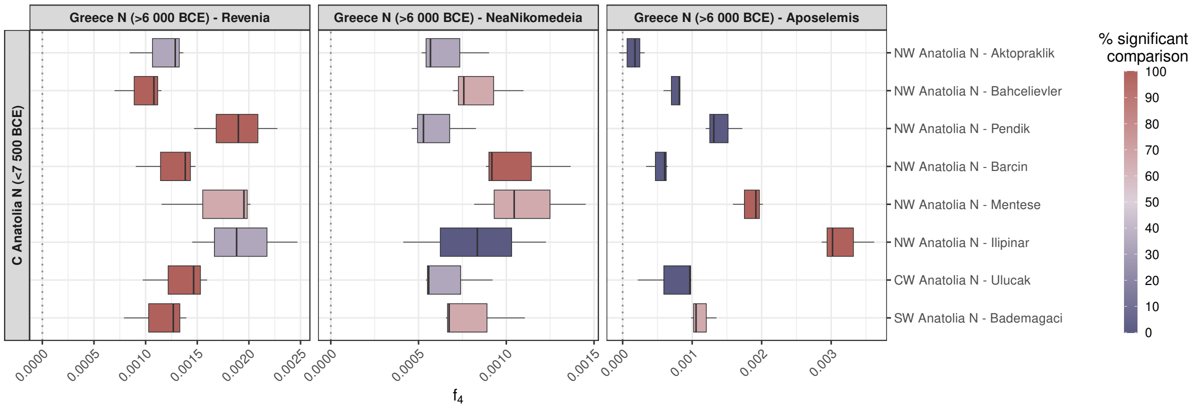


**Figure S21:** Greece N (> 6 000 BCE) populations show a higher affinity to W Anatolia than contemporary C Anatolian populations. The panels show results of $f_4$ tests in the form of $f_4$*(Yoruba, Greece N (> 6 000 BCE); C Anatolia N (<7 500 BCE), W Anatolia N)*. The tests are performed on the population (site) level. The colours indicate the proportion of nominally significant comparisons at |Z|>3.

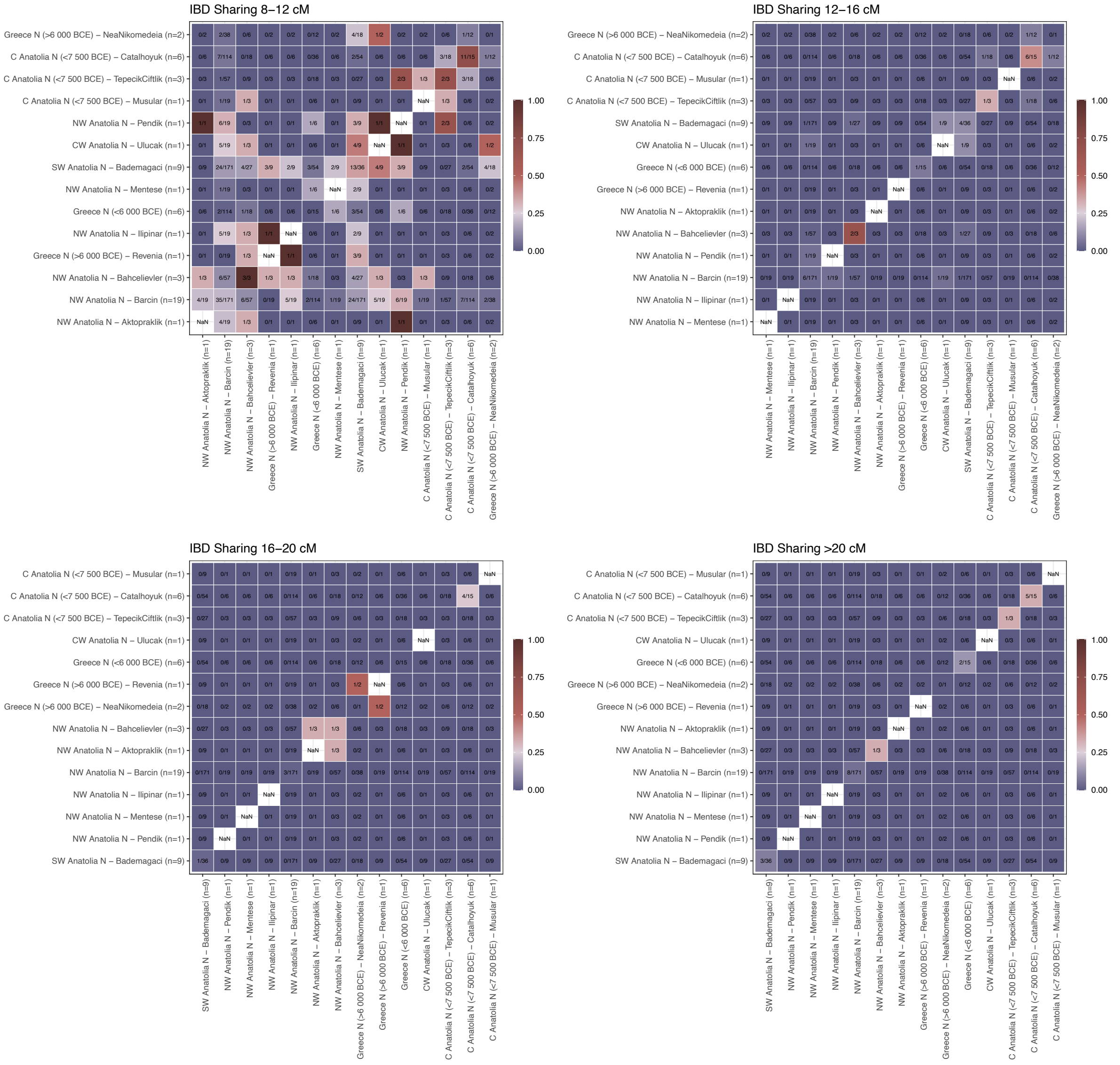


**Figure S22:** Heatmaps showing IBD-sharing among imputed genomes from the study region in the Early Holocene. Each row and column represent a set of genomes from a specific region and period. Each matrix was ordered using hierarchical clustering (with the *hclust* function in R). The colour coding indicates the proportion of sharing, as shown as a fraction within each cell. For instance, the number 24/171 between NW Anatolian Barcin and SW Anatolian Bademagaci in the first panel indicates that NW Anatolian Barcin, represented by imputed 19 genomes, was compared with SW Anatolian Bademagaci represented by 9 imputed genomes, and out of 171 comparisons 24 pairs had at least one IBD segment shared of length 8-12 cM.

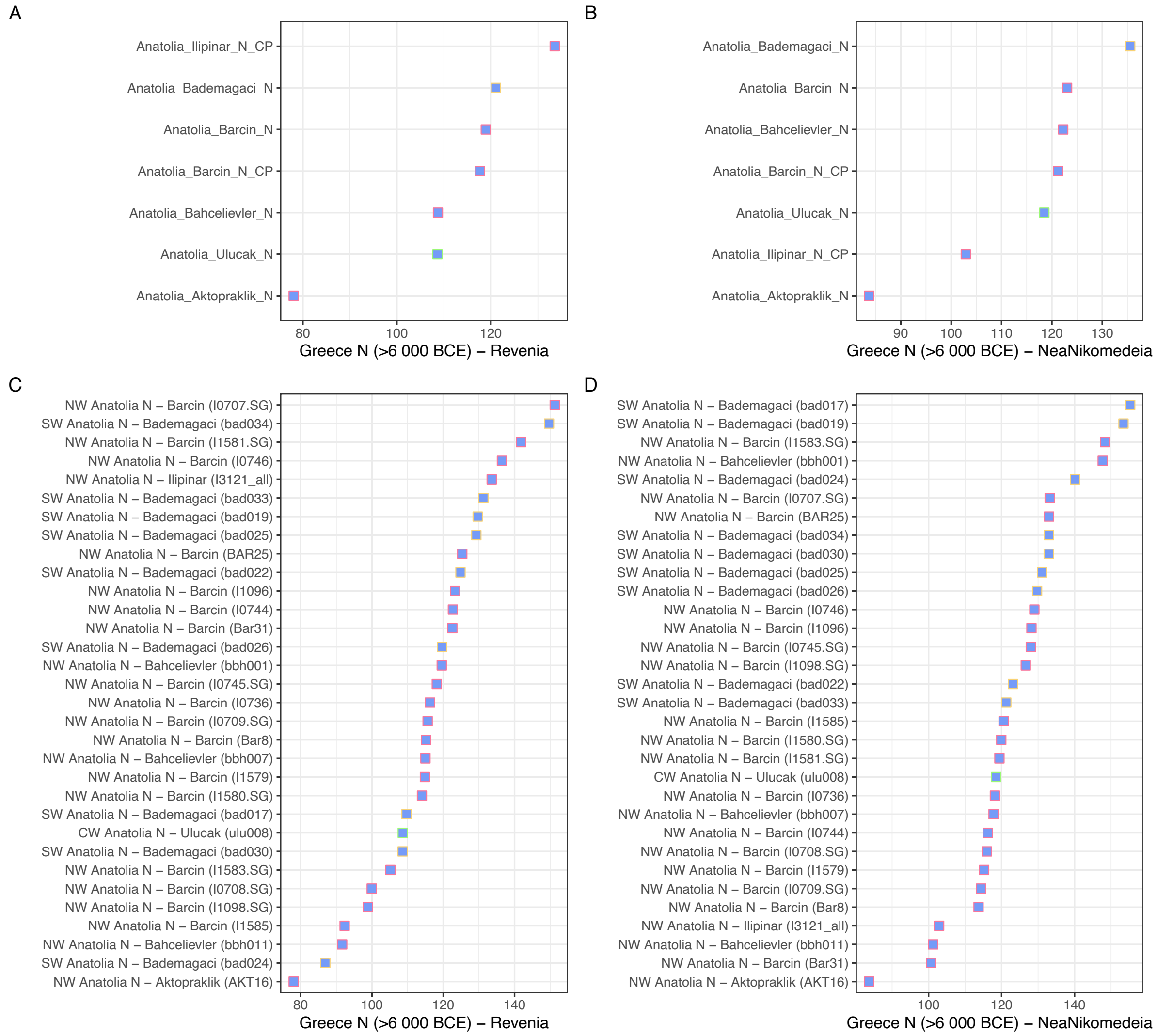


**Figure S23:** Haplotype-sharing statistics between Aegean/Anatolian populations and Revenia (A, C) or Nea Nikomedeia (B, D). Upper panels (A, B) are population means whereas lower panels (C, D) indicate individual-level computations. Shared chunk lengths were estimated in cM by ChromoPainter for each pair of individuals. The Girmeler genome could not be included in this analysis due to its low coverage, which did not allow imputation.

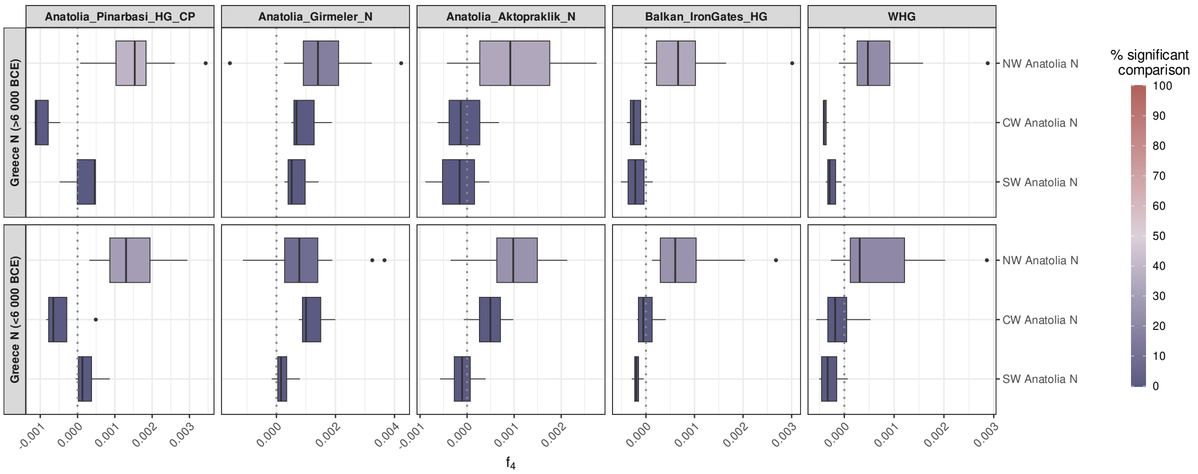


**Figure S24:** The Greece N (> 6 000 BCE or < 6 000 BCE) populations do not show a higher affinity to European Hunter-Gatherers (Balkan HG or WHG) than W Anatolian populations do. The panels show results of $f_4$-tests in the form of $f_4$*(Yoruba, Balkan HG|WHG; Greece N; W Anatolia N)*. The tests are performed on the population (site) level. The colours indicate the proportion of nominally significant comparisons at |Z|>3.

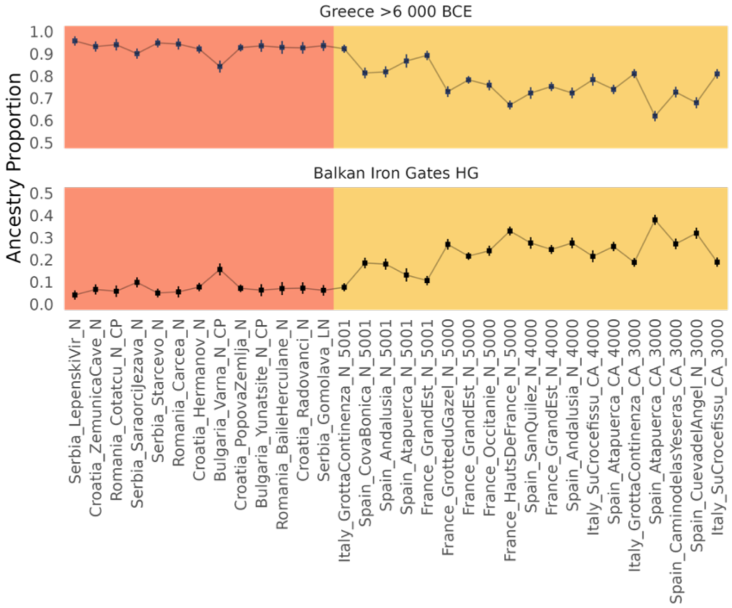


**Figure S25:** The proportion of early Neolithic (>6000 BCE) Greece-related (Revenia) and European Mesolithic-related (Balkan HG) ancestry in European Neolithic communities. The colours indicate sites from Balkan (salmon) and S Europe (yellow). The proportions are based on feasible qpAdm models performed on site-based genomic samples, shown on the x-axis (Methods). The top and lower panels show Revenia and Balkan HG proportions, respectively.

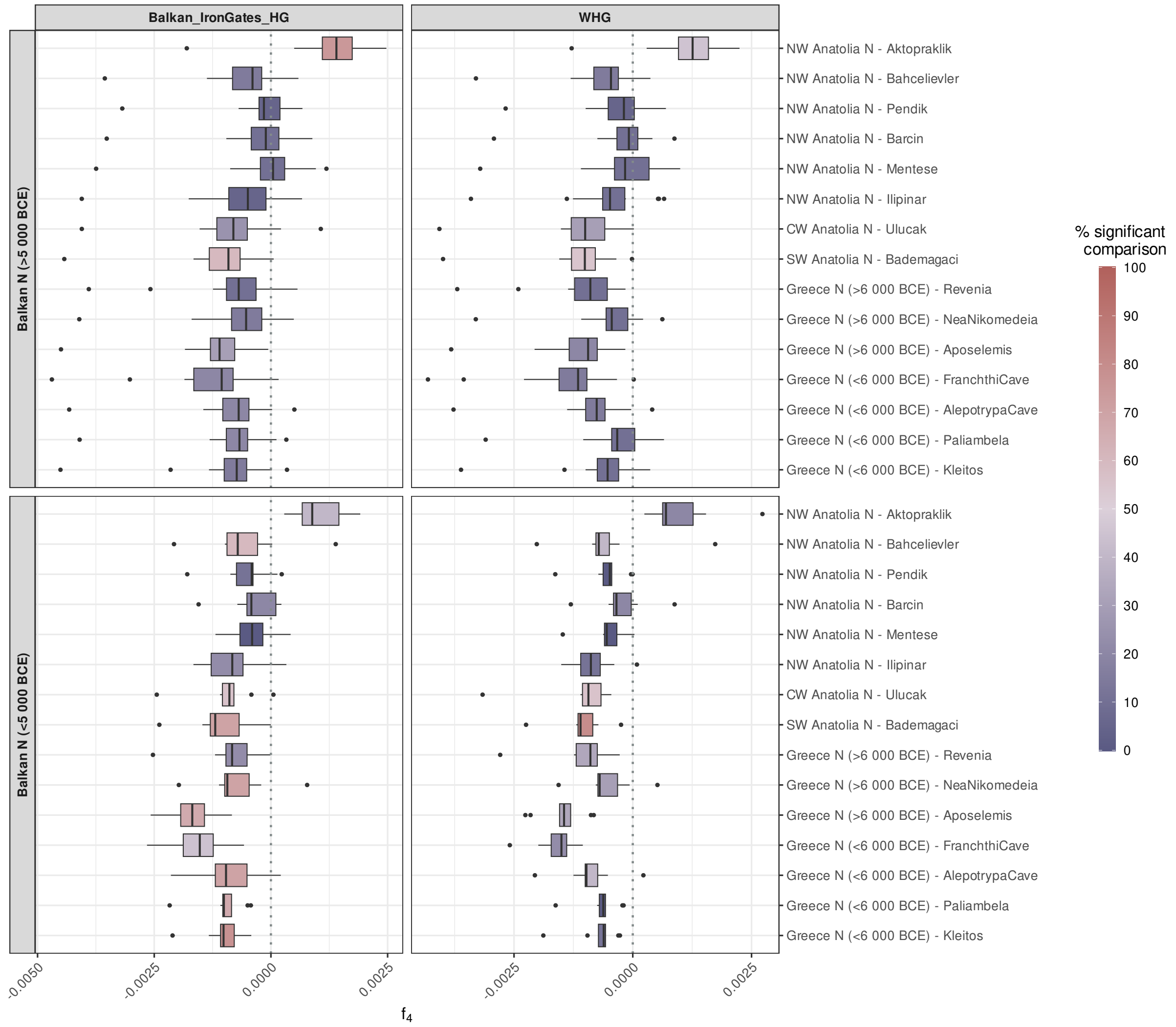


**Figure S26:** The Balkan N populations show a higher affinity to European Hunter-Gatherers (Balkan HG or WHG) than the Aegean populations. The panels show results of $f_4$-tests in the form of $f_4$*(Yoruba, Balkan HG|WHG; Balkan N; Aegean N)*. The tests are performed on the population (site) level. The colours indicate the proportion of nominally significant comparisons at |Z|>3.

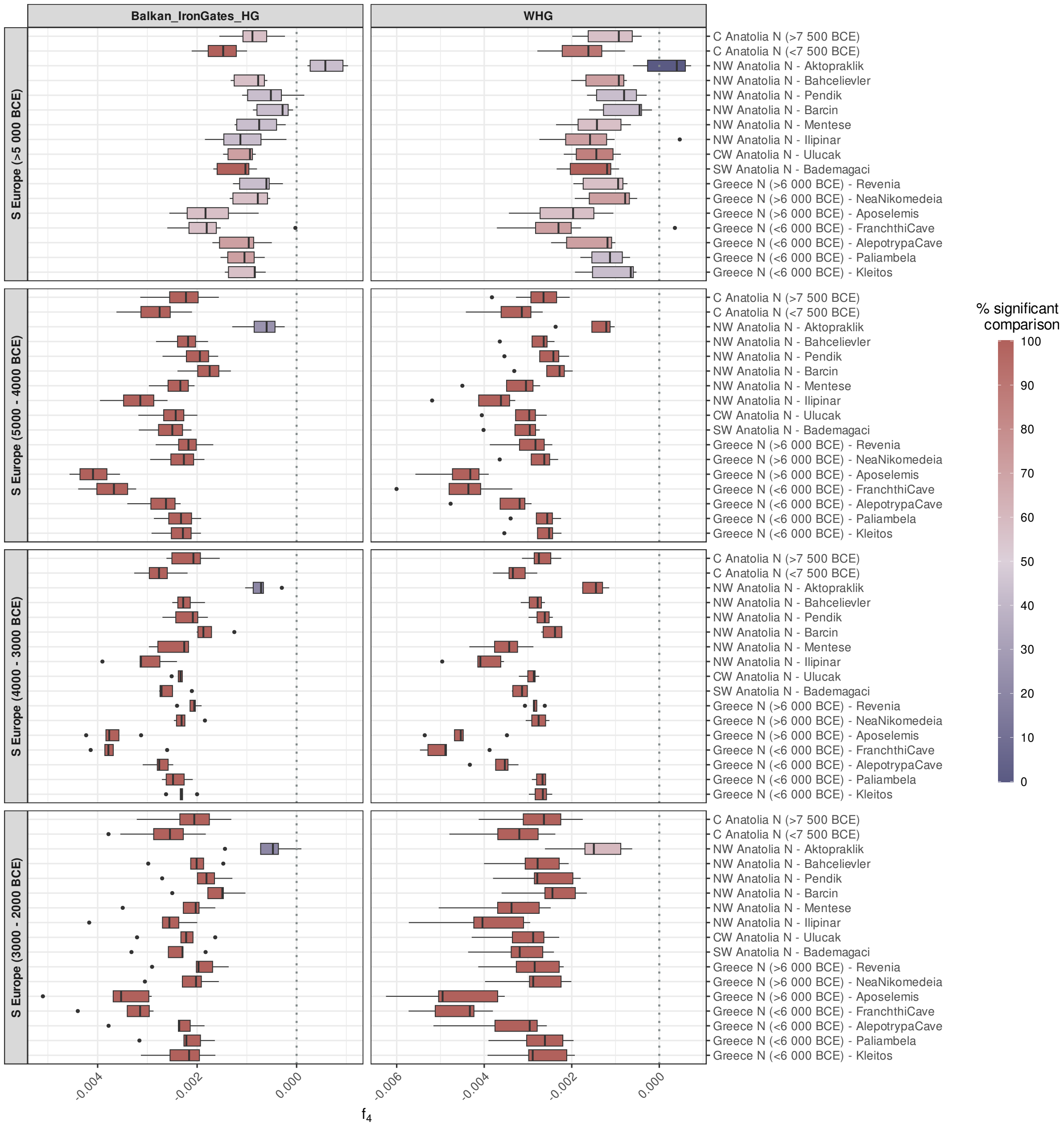


**Figure S27:** The S European N populations show a higher affinity to European Hunter-Gatherers (Balkan HG or WHG) than the Aegean populations. The panels show results of $f_4$-tests in the form of $f_4$*(Yoruba, Balkan HG|WHG; S Europe N; Aegean N)*. The tests are performed on the population (site) level. The colours indicate the proportion of nominally significant comparisons at |Z|>3.

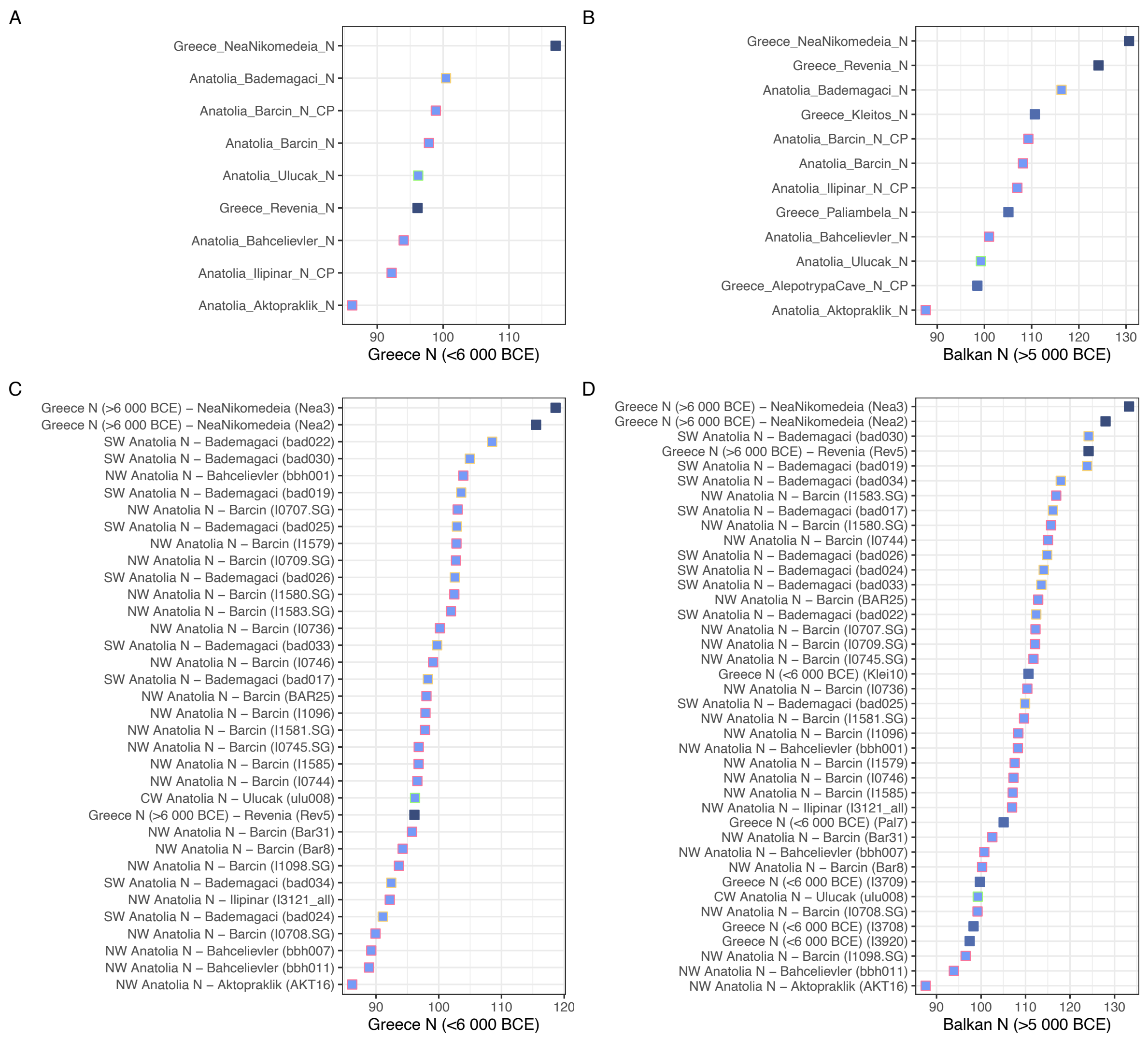


**Figure S28:** Haplotype-sharing statistics between Aegean/Anatolian populations and Greece N (A, C), Balkan N (B, D). Upper panels (A,B) are population means whereas lower panels (C, D) indicate individual-level computations. Shared chunk lengths were estimated in cM by ChromoPainter for each pair of individuals. The Girmeler genome could not be included in this analysis due to its low coverage, which did not allow imputation.

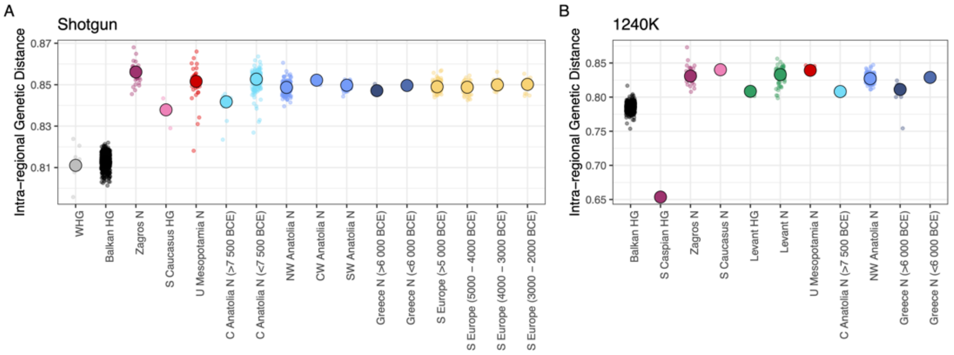


**Figure S29:** $f_3$-based genetic diversity among individuals within a region/time unit. To avoid known batch effects between shotgun and 1240k capture data we performed comparisons on each data type independently.

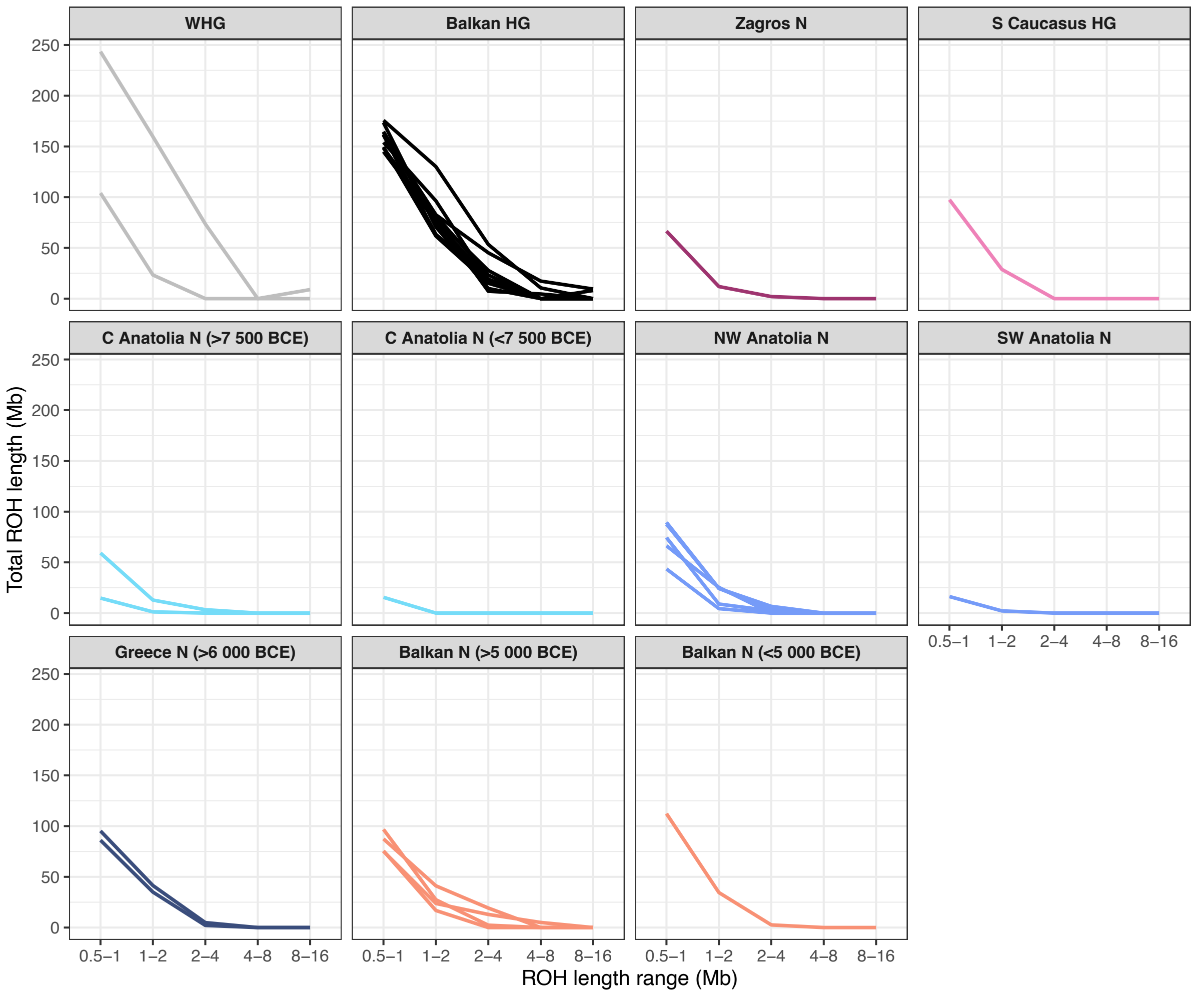


**Figure S30:** Runs of homozygosity (ROH), a measure of inbreeding, calculated using PLINK on W Eurasian genomes if any were >5x.

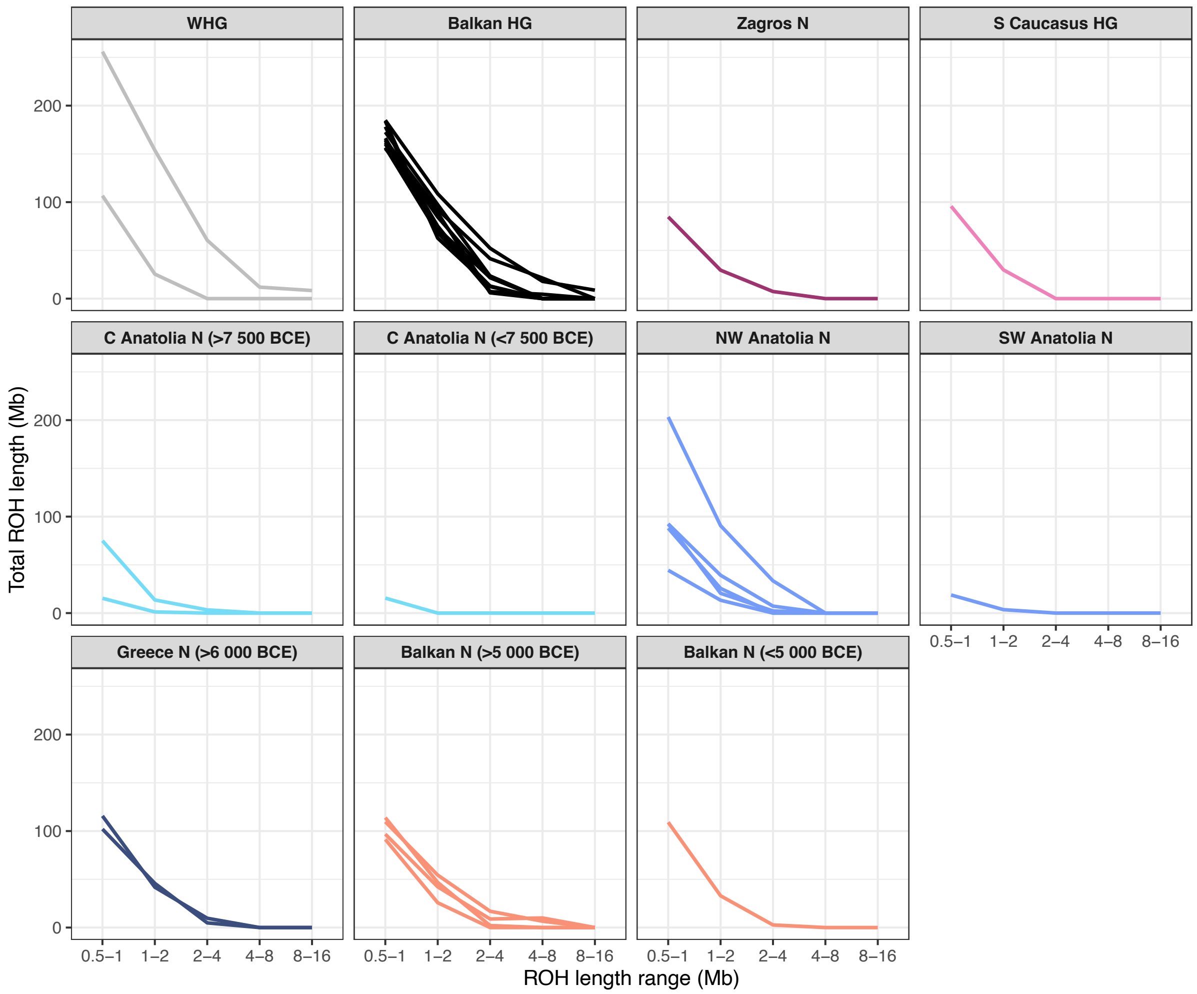


**Figure S31:** Runs of homozygosity (ROH), a measure of inbreeding, calculated using PLINK on W Eurasian genomes. This presents the same information as Figure S30 except that here all genomes were downsampled to 5x.

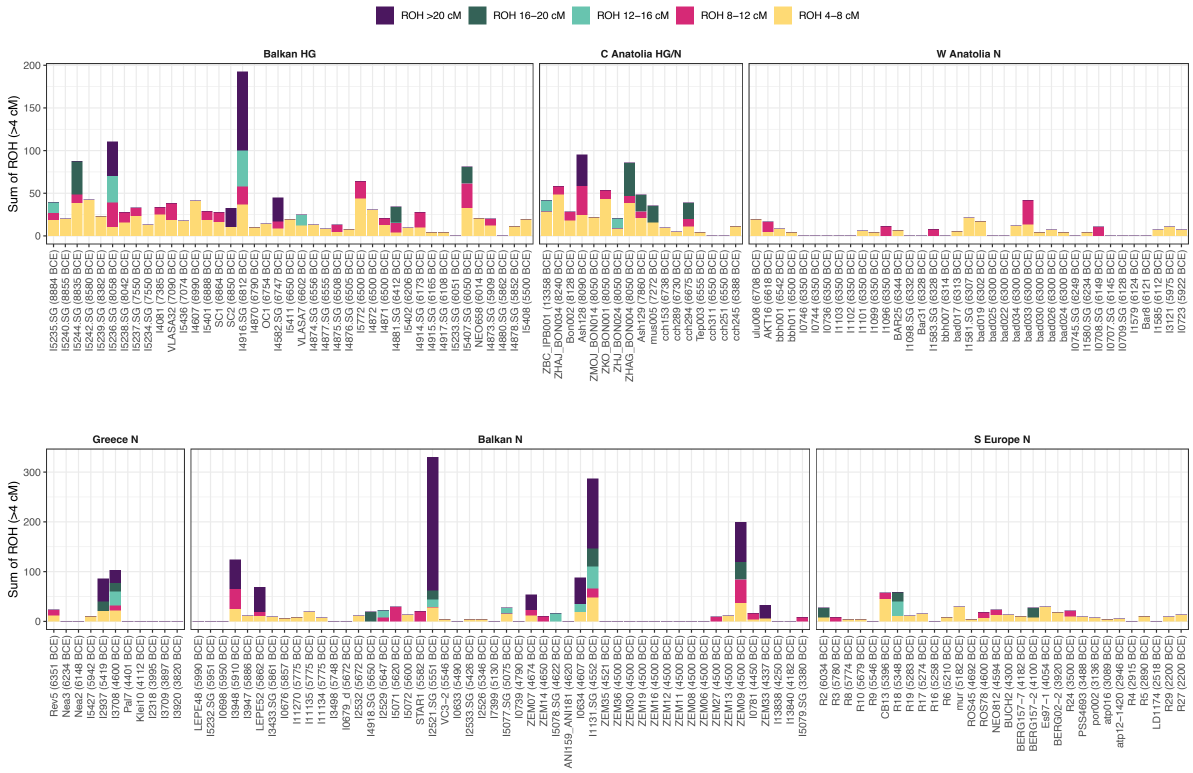


**Figure S32:** Runs of homozygosity (ROH), a measure of inbreeding, calculated using hapROH. Results are plotted by region and ordered by average date of individuals.

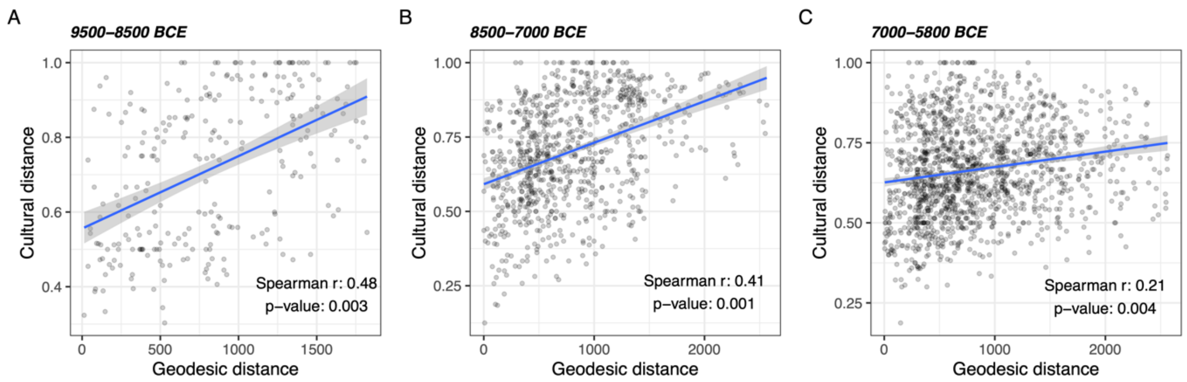


**Figure S33:** Correlations between pairwise cultural (Jaccard dissimilarity) and geographic (geodesic) distances in SW Asia / SE Europe in three periods: 9500-8500 BCE (A), 8500-7000 BCE (B) and 7000-5800 (C). The numbers of archaeological sites used here were 23, 42, and 55, respectively (**Table S6**). The Spearman correlation coefficient and the Mantel test p-values are shown in the plots.

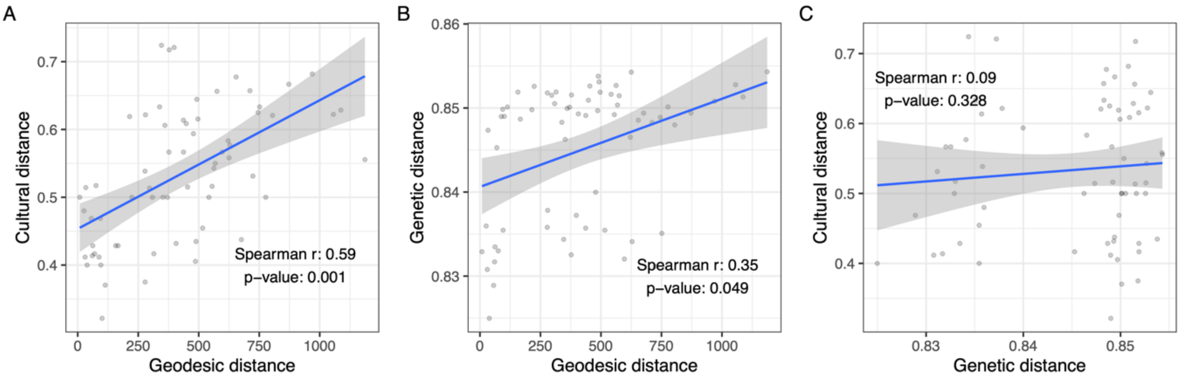


**Figure S34:** Correlations between pairwise (A) cultural-geographic, (B) genetic-geographic and (C) cultural-genetic distances across 12 sites in Greece and Anatolia in the period 7000-5800 BCE. The result of a partial Mantel test examining the relationship between material culture and genetic distances while controlling the confounding effect of geography was not significant (r = -0.15, p = 0.778).

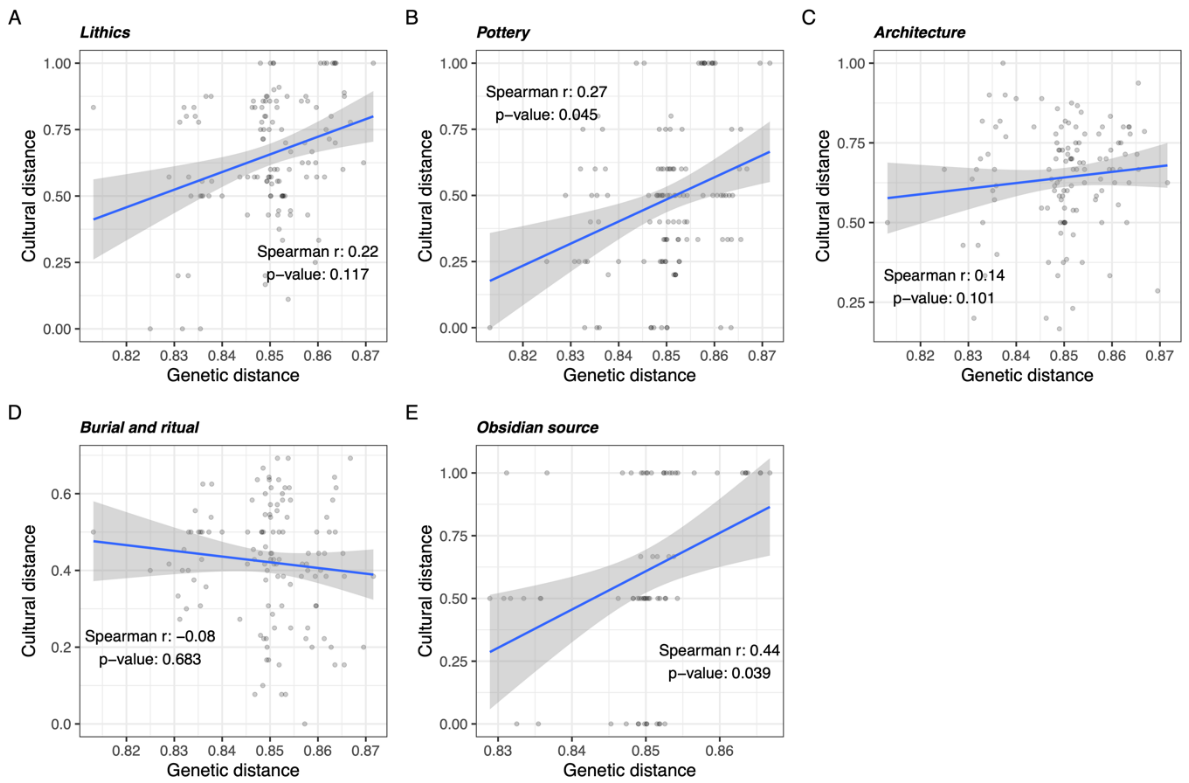


**Figure S35:** Correlations between cultural features and genetic distance in the period 7000-5800 BCE. The features are selected based on the categories (see Figure 6A). The Spearman correlation coefficient and partial Mantel test p-values (controlling for the confounding effect of geography) were as follows: (A) Lithics; r = -0.015, p = 0.538, (B) Pottery; r = -0.08, p = 0.708, (C) Architecture; r = -0.08, p = 0.734, (D) Burial and ritual; r = 0.11, p = 0.261, (E) Obsidian source; r = 0.08, p = 0.382. We removed Ain Ghazal, Menteşe, Nea Nikomedeia and Padina for the comparison based on obsidian source due to the lack of sufficient data.

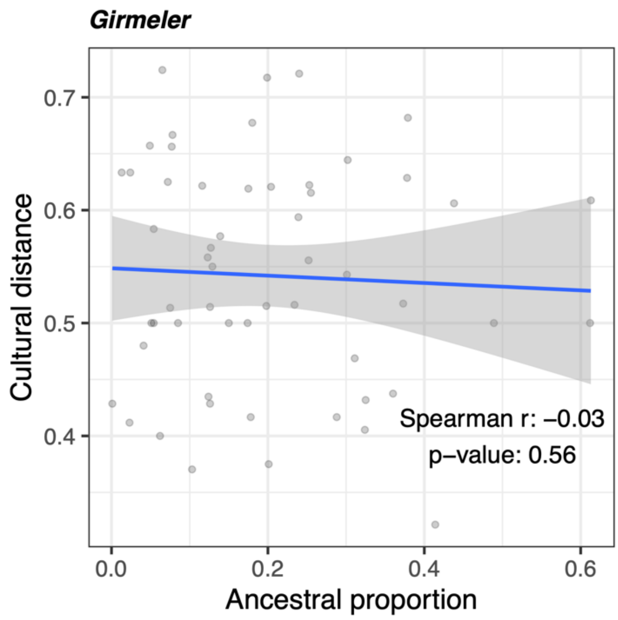


**Figure S36:** Correlation between pairwise cultural distance and difference of Girmeler ancestral proportion across 12 sites in Greece and Anatolia. The result of a partial Mantel test examining the relationship between material culture and genetic distances while controlling the confounding effect of geography was not significant (r = -0.15, p = 0.772).
